# A single-cell 3D spatiotemporal multi-omics atlas from *Drosophila* embryogenesis to metamorphosis

**DOI:** 10.1101/2024.02.06.577903

**Authors:** Mingyue Wang, Qinan Hu, Zhencheng Tu, Lingshi Kong, Jiajun Yao, Rong Xiang, Zhan Chen, Yan Zhao, Yanfei Zhou, Tengxiang Yu, Yuetian Wang, Zihan Jia, Kang Ouyang, Xianzhe Wang, Yinqi Bai, Mingwei Lian, Zhenyu Yang, Tao Yang, Jing Chen, Yunting Huang, Ni Yin, Wenyuan Mo, Wenfu Liang, Chang Liu, Xiumei Lin, Chuanyu Liu, Ying Gu, Wei Chen, Longqi Liu, Xun Xu, Yuhui Hu

## Abstract

The development of a multicellular organism is a highly intricate process tightly regulated by numerous genes and pathways in both spatial and temporal manners. Here, we present Flysta3D, a comprehensive multi-omics atlas of the model organism *Drosophila*, spanning its developmental lifespan from embryo to pupa. Our datasets encompass 3D single-cell spatial transcriptomic, single-cell transcriptomic, and single-cell chromatin accessibility information. By integrating these multi-dimensional data, we constructed cell state trajectories that uncover the detailed profiles of tissue development. With a focus on the central nervous system (CNS) and midgut, we dissected the spatiotemporal dynamics of gene regulatory networks, cell type diversity, and morphological changes from a multi-omics perspective. This extensive atlas provides an unprecedentedly rich resource and serves as a systematic platform for studying *Drosophila* development with integrated single-cell data at an ultra-high spatiotemporal resolution.

## INTRODUCTION

The advances in single-cell multi-omics technologies have revolutionized our understanding of biological processes, revealing cell-specific functional heterogeneities that underlie the complex physiologies of development, aging, and diseases. To date, the functional profile of a single cell can be characterized across multiple dimensions, including its cell surface epitopes, transcriptome, epigenome, and proteome (reviewed in Ref^1^). The development of spatial multi-omics techniques further added spatial context to these dimensions of information (reviewed in Ref^2^), and progress has been made in integrating these multi-modal data to construct a panoramic profile of context-specific functions of single cells and their communications with one another (reviewed in Ref^3^).

*Drosophila melanogaster* has long been a fundamental model organism for genetics and developmental biology research. Recent single-cell multi-omics studies have highlighted the versatility of *Drosophila* in characterizing transcriptomic and epigenomic dynamics of individual cells during embryogenesis^4,5^, tissue development^6–8^, tissue regeneration^9^, and systemic aging^10^. These studies generated rich resources for dissecting the multi-omics profiles of various tissues at single cell precision across developmental stages. Nevertheless, the spatial context of such single-cell omics data is crucial to understanding their biological relevance but is often lost during standard single-cell sequencing procedures.

Embryogenesis is an intricately regulated process that transforms a totipotent zygote into a fully formed embryo with functional organs. Over the past several decades, research into *Drosophila* embryogenesis has yielded invaluable insights into this meticulous process and many of its features that are conserved in mammals (reviewed in Ref^11^). Recently, a few studies have addressed *Drosophila* embryogenesis from the perspective of single-cell multi-omics^4,5^, but focused only on a few developmental time points or are limited in genome coverage. Until recently, genome-wide spatial transcriptomic profiling of developing *Drosophila* was lacking due to the miniature sizes of *Drosophila* samples and resolution limit of spatial transcriptomic techniques. Previously, we utilized spatial enhanced resolution omics sequencing (Stereo-seq)^12^, a sequencing- and patterned DNA nanoball (DNB) array-based spatial transcriptomic platform with high spatial resolution and sensitivity, to address this gap. Using Stereo-seq, we generated 3D spatiotemporal transcriptomic maps of *Drosophila* late-stage embryos and larvae and analyzed the development of tissues within their actual 3D spatial context^13^.

Here, we expanded our previous spatiotemporal transcriptomic atlas of *Drosophila* to cover its developmental lifespan from embryo to pupa. Using Stereo-seq and *Spateo*, a computational pipeline designed to analyze single-cell multi-modal spatial transcriptomic data^14^, we reconstructed 3D transcriptomes at single cell spatial resolution. We further complemented embryo single-cell Stereo-seq (scStereo-seq) data with single-cell RNA sequencing (scRNA-seq) and single-cell assay for transposase-accessible chromatin using sequencing (scATAC-seq) data to create a multi-omics atlas of *Drosophila* embryos that includes transcriptomic and epigenomic information within an ultra-high-resolution spatial context. The data in this single-cell spatiotemporal multi-omics atlas of *Drosophila* development are curated in our database, Flysta3D, for easy access.

Based on the unprecedentedly rich data resource, we established multi-omics cell state trajectories of tissue development. Along these trajectories, we systematically characterized the spatiotemporal dynamics of cell differentiation potential, signaling pathways, and transcription factor (TF) regulatory networks. Focusing on two widely studied *Drosophila* tissues, central nervous system (CNS) and midgut, we delved into their cell type diversification, gene regulatory networks, and morphological changes from a multi-omics perspective. Given that we have produced extensive multi-omics datasets for the embryonic stages, the major focus of this paper will be the analysis of embryogenesis from a multi-omics perspective. The scStereo-seq data for the larval and pupal stages are not discussed extensively here but will be accessible via Flysta3D database. Flysta3D hosts all the datasets generated in this study and provides interactive 3D visualization of gene expression patterns, TF regulatory networks, signaling pathway activities, etc. in these datasets. Our database can facilitate systematic research on *Drosophila* development with its comprehensive information and broad range of applications.

## RESULTS

### Single-cell 3D spatial transcriptomes of *Drosophila* from embryogenesis to metamorphosis

To construct multi-omics atlas of *Drosophila* development, we started off by expanding and enhancing the 3D spatial transcriptomes of *Drosophila* development based on our previous work. Developing embryos were collected at 0.5 to 2 h intervals throughout the ∼24 h course of embryogenesis (hereafter termed based on computationally inferred developmental age, see below). Larva samples were collected at early or late time points for each of the three larval stages (hereafter termed L1 to L3 early/late). Pupa samples were collected at 12 h intervals starting from pupation (hereafter termed P12 to P72) **(Figure 1A and Table S1)**. Cryosection was performed for each sample to obtain their sagittal sections of 7 or 8 μm thickness, and all available sections of each sample were subjected to *in situ* mRNA capture, library preparation, and sequencing at the Stereo-seq platform.

**Figure 1.**
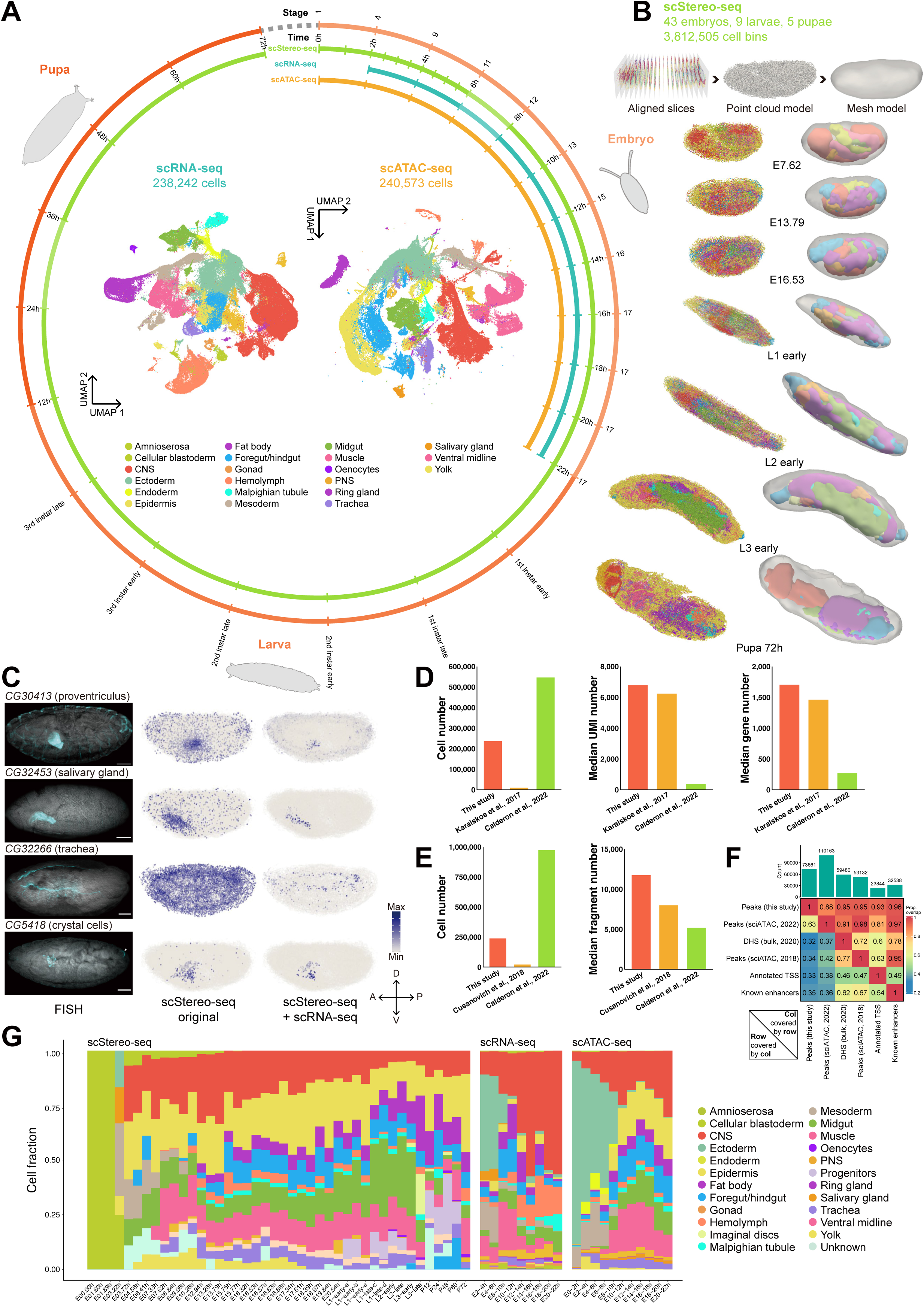
A single-cell spatiotemporal multi-omics atlas of developing Drosophila. **(A)** Samples covered in this study. The outer rim indicates sample collection windows for three omics datasets, with each arc segment represents a collection window. Time points indicate hours after egg laying in embryos and hours after pupation in pupa. The inner panel shows UMAP plots of aggregated scRNA-seq and scATAC-seq data, color coded by tissue annotation. We did not obtain quality P36 scStereo-seq and E6-8h scRNA-seq data. CNS: central nervous system; PNS: peripheral nervous system. **(B)** 3D modeling of representative scStereo-seq samples using *Spateo*, showing point cloud (left) and mesh (right) models for the entire animal over developing stages. Models of epidermis, trachea, hemolymph, and muscle are not displayed in some samples for better visualization of internal organs. Tissue color codes are the same as **(A)**. Samples are not on the same scale. **(C)** FISH validation of representative genes from the list of genes without reported spatial expression patterns **(Table S3)**. For each gene, representative FISH images were obtained from stage 11-16 embryos from lateral or near-lateral view. Cyan: gene-specific RNA probes; grey: nuclei stained with DAPI. Arrowheads indicate structures with autofluorescence (e.g., trachea). Scale bars = 50 μm. All scStereo-seq samples are shown in lateral or near-lateral view. A-P: anterior-posterior; D-V: dorsal-ventral. Spatial expression patterns generated from original scStereo-seq or integrated scStereo-seq and scRNA-seq data are also from representative stage 13-17 embryos, projected along the Z-axis. See additional examples in **Figure S1C**. **(D)** Quality benchmark of scRNA-seq dataset in this study, showing cell number, median UMI number per cell, and median gene number per cell in datasets from this study and previous *Drosophila* embryo scRNA-seq studies. **(E)** Quality benchmark of scATAC-seq dataset in this study, showing cell number and median fragment number per cell from this study and previous *Drosophila* embryo scATAC-seq studies. **(F)** Heatmap showing proportion of scATAC-seq peaks in this study overlapping peaks in two previous *Drosophila* embryo scATAC-seq/scATAC-seq studies, bulk DHS peaks, and peaks in known TSSs and enhancers. **(G)** Bar plot showing cell type composition of data from scStereo-seq (some low-quality samples are filtered), scRNA-seq, and scATAC-seq over sample collection time. The y-axes are fraction of cell types annotated in each dataset. The x-axes are sample collection time points/windows (*RAPToR* inferred developmental age for embryo scStereo-seq data).

The subcellular spatial resolution achieved by Stereo-seq (∼500 nm distance between DNBs) was not fully utilized in our previous dataset due to the lack of cell location information. Here, to address this, we attempted to achieve single-cell spatial resolution by nucleus staining and imaging of each Stereo-seq chip before library preparation. Cell segmentation was then performed based on the location of each nucleus. After sequencing and mapping, 2D spatial gene expression matrices were aligned with segmented images. Each DNB was then assigned to a cell bin, allowing for more precise single-cell transcriptome analysis **(Figure S1A)**. We then combined cell bins from all sections of individual samples, performed unsupervised clustering based on both gene expression profiles and spatial locations **(Data S1)**, and manually annotated the clusters according to marker gene expression and spatial morphology **(Table S2)**. Utilizing the Stereo-seq platform, we generated organism-wide single-cell spatial transcriptomes for 43 embryo, 9 larva, and 5 pupa samples throughout *Drosophila* development, with a total of 3,812,505 cell bins **(Table S1)**.

With the 2D single-cell spatial transcriptomic datasets, we reconstructed the spatial transcriptomes in 3D leveraging point cloud-based modeling method in *Spateo* package, which were optimized for cell bins. This approach offered enhanced structural details compared to our previous 3D modeling results and allowed the alignment of 3D models from different time points for morphometric analysis (see below). The 3D modeling of cell bin spatial transcriptomic data effectively captured the anatomical morphology of tissues with finer details than our previous 3D models **(Figure 1B and Figure S1B)**. In our previous study on limited embryo and larva samples, we demonstrated that Stereo-seq data reproducibly captured spatial gene expression patterns that largely overlapped with established *in situ* databases^15,16^, as well as those that were absent in these databases. Based on this more comprehensive spatiotemporal transcriptomic dataset, we further identified a list of 338 genes without reported spatial expression patterns in embryos and reconstructed their patterns in 3D **(Table S3)**. We selected 9 genes to validate their spatial expression patterns with fluorescence *in situ* hybridization (FISH) and found high consistency with scStereo-seq data in terms of spatial gene expression patterns and tissue enrichment **(Figure 1C and Figure S1C)**. These results further substantiated the power of Stereo-seq in recapitulating spatial gene expression profiles and guiding *in vivo* validation.

### A single-cell spatiotemporal multi-omics atlas of *Drosophila* embryogenesis

Despite its ability to provide a better representation of single-cell spatial transcriptomes, scStereo-seq had a higher dropout rate compared to droplet-based scRNA-seq due to a reduced number of DNBs assigned to each cell bin. This limitation curtailed the ability of scStereo-seq to detect genes that express at a lower level. To overcome this drawback and to augment our single-cell 3D spatial transcriptomic data with deeper transcriptomic and epigenomic information, we collected samples at 2-hour intervals across embryogenesis and performed droplet-based scRNA-seq and scATAC-seq **(Figure 1A)**. Following quality control, we obtained 238,242 single-cell transcriptomes with scRNA-seq, with a median of 6,841 unique molecular identifiers (UMIs) and 1,707 genes per cell **(Table S1)**. These quality control statistics in our scRNA-seq data were comparable to or better than previous *Drosophila* embryo scRNA-seq studies^4,5^ **(Figure 1D)**. Additionally, we obtained 240,573 single-cell chromatin accessibility profiles with scATAC-seq, with a median of 11,772 fragments per cell **(Table S1)**. The number of fragments captured per cell and other quality control statistics in our scATAC-seq data were also comparable to or better than previous *Drosophila* scATAC-seq datasets^5,17^ **(Figure 1E and Figure S1D-F)** and achieved high coverage of previously reported scATAC-seq datasets^5,17^, DNase I hypersensitive sites (DHS)^18^, annotated transcription start sites (TSS)^19^, and known enhancer sites^20–22^ **(Figure 1F)**.

With the aggregated scRNA-seq data collected across embryogenesis, we performed an initial round of coarse unsupervised clustering and generated 45 cell clusters in the uniform manifold approximation and projection (UMAP) plot **(Figure S1G)**. We annotated these clusters and classified annotations at three levels (cell type/tissue substructure - tissue - germ layer, e.g., gastric caecum - midgut - endoderm) based on marker gene expression **(Figure 1A and Table S2)**. Similarly, we performed coarse unsupervised clustering in aggregated scATAC-seq data, generating 40 distinct clusters in the UMAP plot **(Figure S1H)**. Each cluster was also annotated through inspection of marker genes **(Figure 1A and Table S2)**. The data we collected achieved extensive coverage of major tissues, as reflected by the proportion of cells representing each tissue and their dynamics over developmental stages **(Figure 1G)**.

Given the deep genome coverage of our data, we further profiled tissue cell type heterogeneity by subclustering and annotating tissue clusters from scRNA-seq and scATAC-seq data. The resolution of subclustering for each tissue was determined based on previously reported cell type complexities. Detailed cell types were annotated based on marker gene expression and literature search **(Table S4)**. Through subclustering of the scRNA-seq data, we were able to examine the specific composition of embryo tissue cell types. For example, the subclustering of the peripheral nervous system (PNS) cluster allowed for the distinct identification of neurons and glia from external sensory^23^ and chordotonal organs^24,25^ **(Figure S2)**. These subclustering results indicated that we were able to extensively characterize the major cell types in the embryo and identify several rare ones, such as adult midgut progenitors (AMPs)^26^ and entero-endocrine cells (EEs)^27^ in the midgut (see below). We also identified subclusters representing most of these detailed cell types in the scATAC-seq data **(Figure S3)**. While we annotated tissue subclusters to the best of our knowledge, there could still be instances where clusters were not assigned their optimal annotations. We annotated some of the ambiguous or unknown cell clusters with reference to cell types they resembled most based on marker genes, such as “neuron-like” **(Figure S2 and S3)**. Therefore, community efforts are welcome to help further specify the annotations of tissue cell types. To verify the well annotated subclusters we identified in both datasets, we compiled a list of common tissue substructure/cell type markers, which are identified in both datasets **(Table S5)** and validated the expression specificity of 3 previously unreported cell type markers using FISH **(Figure 2A)**.

**Figure 2.**
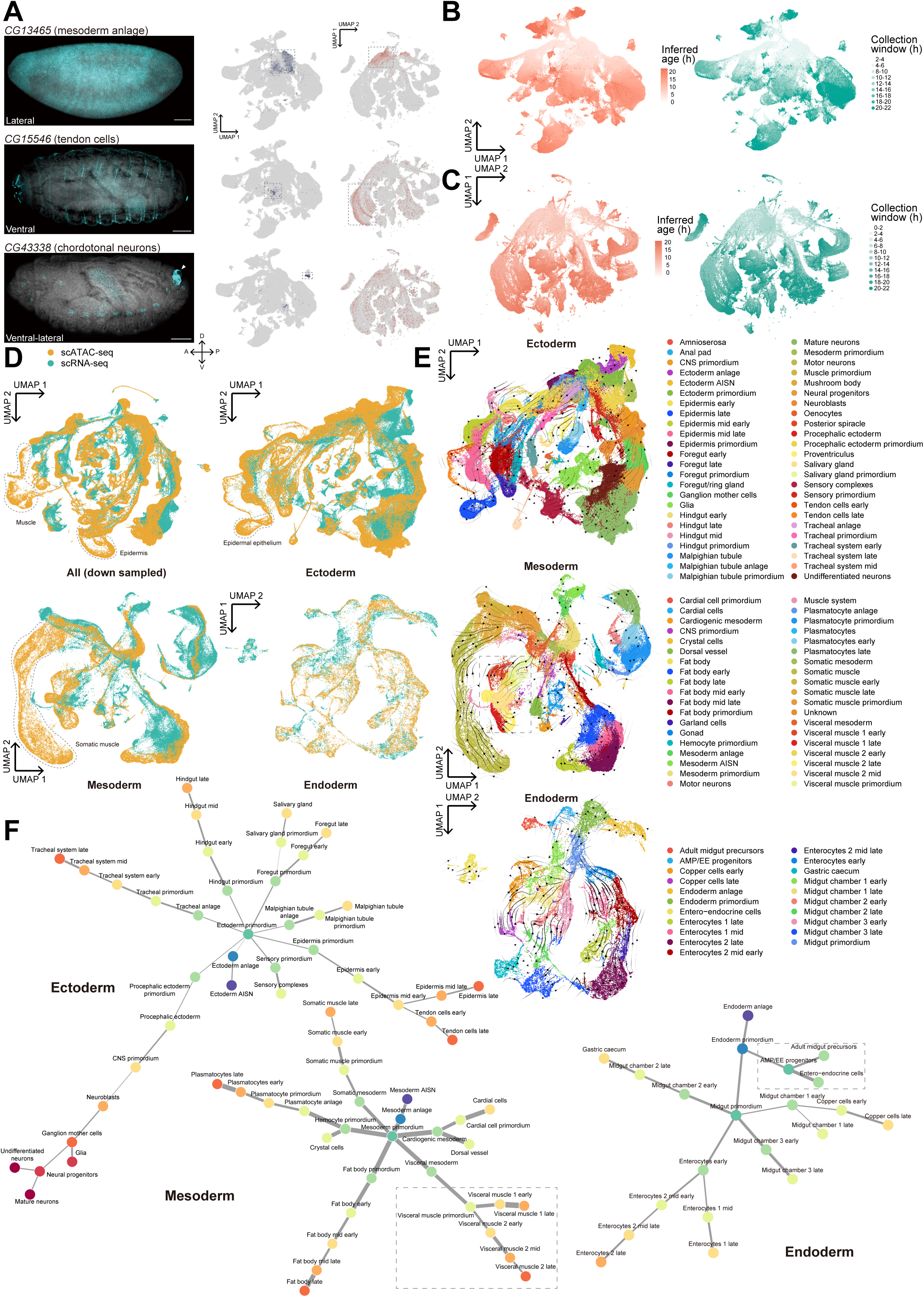
Integration of scRNA-seq and scATAC-seq data for construction of tissue development trajectories. **(A)** FISH validation of representative genes in the list of common tissue substructure/cell types **(Table S5)**. Left: representative FISH images of corresponding stages of gene expression enrichment, with sample viewpoints labeled. Cyan: gene-specific RNA probes; grey: nuclei stained with DAPI. Arrowheads indicate structures with autofluorescence (e.g., trachea). A-P: anterior-posterior; D-V: dorsal-ventral. Scale bars = 50 μm; Right: UMAP plots of marker gene expression specificity in aggregated scRNA-seq and scATAC-seq data. Cells with enriched marker gene expression/peak accessibility are highlighted in dashed rectangles. **(B)** UMAP plots of aggregated scRNA-seq data, color coded with *RAPToR* inferred developmental age (left) and actual sample collection window (right). **(C)** UMAP plots of aggregated scATAC-seq data, color coded with neural network model inferred developmental age (left) and actual sample collection window (right). **(D)** UMAP plots of co-embedded scRNA-seq and scATAC-seq data of all cells (down sampled) and three germ layers. Dashed lines mark cell clusters in scATAC-seq data that miss corresponding cells in scRNA-seq data, with their scATAC-seq annotations labeled. **(E)** Velocity fields of co-embedded UMAP plots of three germ layers in **(D)**, color coded with re-annotated cell types based on clustering of integrated data. Velocity trajectories point backward from chronologically older to younger cells. The dashed rectangle indicates visceral muscle groups discussed in the following analyses. AISN: anlage in statu nascendi. **(F)** Tissue development trajectories based on cluster phylogeny inferred from **(E)** for major tissues of three germ layers. Within each germ layer, widths of lines connecting subcluster annotations indicate gene expression similarities. Dashed rectangles indicate visceral muscle trajectories discussed in GRN analysis and AMP/EE trajectories discussed in midgut cell type identification.

In summary, we generated a compendium of scStereo-seq, scRNA-seq, and scATAC-seq datasets throughout *Drosophila* embryogenesis. The high granularity and temporal continuity of our multi-omics data opened the possibility of cell type- and developmental age-dependent integration of these multi-omics data.

### Developmental age-matched integration of multi-omics data

Integration of multi-omics data offers more comprehensive perspectives when searching for key molecular factors regulating tissue development. Our multi-omics data were obtained from multiple batches of samples using different techniques. Considering the rapid spatiotemporal gene expression changes during embryogenesis, it is crucial to confirm that the developmental ages of samples were matched before integrating multi-omics data. Age matching was also necessitated by the fact that mated female flies might retain embryos in their reproductive tract for some time between fertilization and egg laying (“egg retention”)^28^, leading to possible deviations of the actual developmental age from the sample collection windows in individual scStereo-seq samples.

To precisely stage our samples, we applied *RAPToR*, a predictive model inferring the developmental age of biological samples based on transcriptomic profiles^29^, on both embryo scStereo-seq and scRNA-seq data. The developmental age of embryo scStereo-seq samples was determined by *RAPToR* inference with the entire embryo as a pseudo-bulk input **(Table S1)**. The ages inferred by *RAPToR* aligned well with the collection window for the majority of samples. However, in a few instances, there was a significant discrepancy, with *RAPToR* inferring an age notably older than that derived from the collection window, indicating the presence of female egg retention **(Table S1)**. We further validated the age of these samples by inspecting their nuclear staining morphologies and found better agreement with *RAPToR* inference than collection window. Consequently, these samples are denoted by their *RAPToR*-inferred developmental age rather than the actual sample collection time window (e.g., E15.75 refers to an embryo sample with the inferred developmental age of 15.75 h). The resulting set of 43 embryo scStereo-seq samples we collected comprehensively covered *Drosophila* embryogenesis **(Figure S4A)**. The *RAPToR*-inferred developmental age of single cells from scRNA-seq data showed overall good agreement with their actual sample collection window **(Figure 2B and Figure S4B)**, but with significant tissue-dependent variations within each stage **(Figure S4C)**, likely because *RAPToR*, a model trained on bulk RNA-seq data, lacks cell type specificity for scRNA-seq data. Similar variations were also observed in age inference of individual cell bins of scStereo-seq samples **(Figure S4D)**. To infer the developmental age of cells from scATAC-seq data, we employed a previously described neural network model^5^. The resulting cell developmental age was also largely consistent with the sample collection window **(Figure 2C and Figure S4E)**.

Subsequently, we selected cells in the scRNA-seq data with a *RAPToR* inferred developmental age difference of 1 hour for integration with scStereo-seq data using *NovoSpaRc*^30^. The integrated data enabled the imputation of spatial gene expression patterns with higher genome coverage, yielding markedly reduced signal background, enhanced tissue enrichment, and improved spatial gene expression patterns that exhibited greater resemblance to FISH validation results **(Figure 1C and Figure S1C)**.

We acquired scRNA-seq and scATAC-seq data from several hundred embryos per sample batch. Sample developmental age matched collection window for the majority of embryos, and the substantial sample size mitigated the influence of female egg retention. This was reinforced by the consistency between the model-predicted age and the actual collection window in both datasets **(Figure 2B-C)**. Additionally, the developmental ages of scRNA-seq and scATAC-seq were inferred with different models and might not be comparable. Consequently, we directly used sample collection window to integrate the scRNA-seq and scATAC-seq data for downstream analysis.

Together, our three multi-omics datasets exhibited coherence during embryo stages and can be integrated in a developmental age-specific manner in downstream analyses.

### Construction of multi-omics tissue development trajectories

To delve into the detailed dynamics of developing tissues, we aimed to chronologically organize the tissue cell types in scRNA-seq and scATAC-seq data into continuous tissue developmental trajectories. Upon examining the subclustered and annotated data, we noticed that certain developmentally transitional cell types were categorized into different tissue clusters between assays, possibly due to differences in assay techniques, genome coverage, or clustering resolution (e.g., “muscle primordium” was annotated in the “mesoderm” cluster of the scATAC-seq dataset but in the “muscle” cluster of the scRNA-seq dataset) **(Figure S2 and Figure S3)**. To resolve this issue, instead of focusing on individual tissues, we included all cells of the same germ layer for collective and continuous analysis.

We first integrated the scRNA-seq and scATAC-seq data by finding integrated anchors for label transfer, imputing gene expression matrix from peak matrix of scATAC-seq data, and co-embedding them in the same UMAP space **(Figure 2D)**. Subsequently, unsupervised clustering was performed on the integrated germ layer data **(Figure S4F)**. Notably, a substantial number of late-stage cells annotated as “muscle” and “epidermis” in the scATAC-seq data did not correspond to any cell clusters in the scRNA-seq data **(Figure 2D)**. This discrepancy likely stemmed from technical limitations in capturing late-stage muscle (possibly due to their syncytial characteristics) and epidermal cells with our droplet-based scRNA-seq procedure. This was supported by a significant decrease in the fraction of these two cell types in late-stage scRNA-seq data **(Figure 1G)**.

To construct tissue developmental trajectories within a germ layer, we applied *PhyloVelo*^31^ to the integrated data to establish velocity vector fields for three germ layers and re-annotated cell clusters based on marker genes and their chronological order along the velocity trajectories **(Figure 2E and Table S4)**. Due to their complexities, the trends of cell type differentiation are better visualized in 3D UMAP plots **(Data S2)**. With these velocity vector fields, we delineated multi-omics tissue development trajectories for all three germ layers **(Figure 2F)**.

Multi-omics tissue development trajectories allowed continuous and systematic tracing of various aspects of tissue- and cell type-specific dynamics during embryo development. To assess the differentiation dynamics of single cells, we employed *CytoTRACE*^32^, which leveraged the number of detectably expressed genes as a robust indicator of differentiation potential. *CytoTRACE* analysis revealed diverse trends in differentiation dynamics across tissues during organ specification and maturation **(Figure 3A-B and Figure S4G)**. In general, mesodermal and endodermal tissues exhibited a slower decrease in differentiation potential compared to ectodermal ones. As anticipated, gonad cells maintained a consistently high level of potential throughout embryogenesis. Notably, the nervous system displayed the most rapid decline in differentiation potential throughout development, indicating its relatively faster pace towards terminal differentiation **(Figure 3A)**. Within each tissue, different cell types also exhibited varying rates of reduction in differentiation potential during development **(Figure 3B and Figure S4G)**. Genes whose expression level showed the strongest positive correlation with *CytoTRACE* scores were enriched in cell differentiation-related and ribosome protein genes **(Figure S4H-I and Table S6)**. The latter has been previously reported as indicators of both differentiation potential (reviewed in Ref^33^) and aging^10^. Conversely, genes whose expression level most negatively correlated with *CytoTRACE* scores included specific markers of differentiated tissues (e.g., GABAergic neuron-specific marker *Rdl*^34^ and hemocyte-specific marker *Ppn*^35^) **(Figure S4H)**.

**Figure 3.**
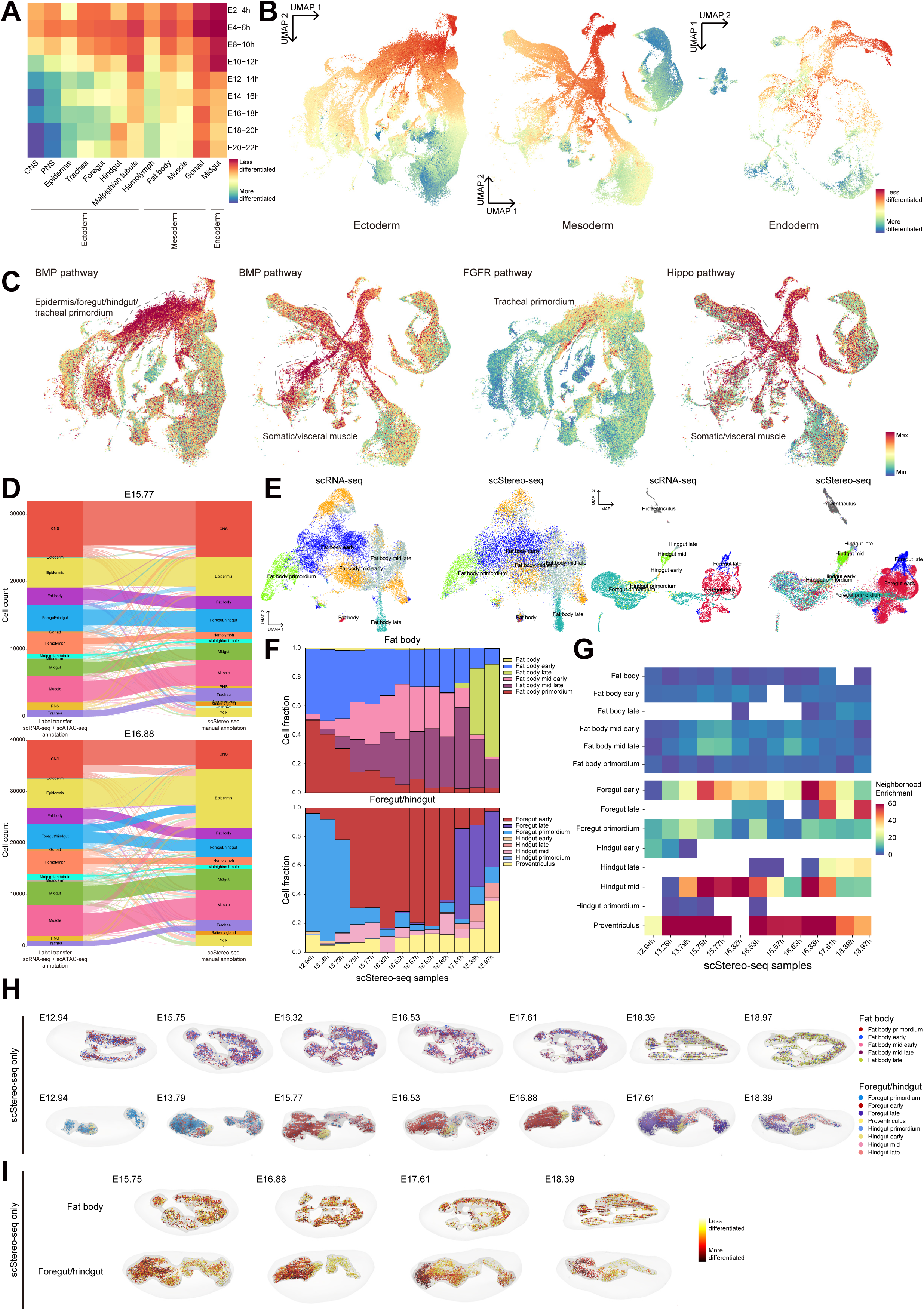
Spatiotemporal dynamics along multi-omics tissue developmental trajectories. **(A)** Heatmap showing median tissue *CytoTRACE* scores based on scRNA-seq data along tissue development trajectories. *CytoTRACE* scores are scaled across all cells. **(B)** UMAP plots of scRNA-seq cells in the co-embedded UMAP space in Figure 2E, color coded with *CytoTRACE* scores. *CytoTRACE* scores are scaled within each germ layer. **(C)** Same as **(B)** but color coded with gene activity scores of core components of signaling pathways. Representative tissues enriched in signaling pathway activities are labeled. **(D)** Sankey plots showing agreement between scStereo-seq tissue manual annotations and transferred labels from integrated scRNA-seq and scATAC-seq data in representative scStereo-seq samples. **(E)** Co-embedding of fat body and foregut/hindgut cells from scRNA-seq and scStereo-seq (pooled samples) data in the same UMAP plots, labeled with original scRNA-seq annotations or transferred annotations. **(F)** Bar plots showing cell type composition of fat body and foregut/hindgut in scStereo-seq samples. Cell types are label transferred from scRNA-seq data. **(G)** Heatmaps showing neighborhood enrichment scores of fat body and foregut/hindgut cell types across scStereo-seq samples. Blank cells indicate absence of label transferred cell types or lack of enrichment in corresponding samples. **(H)** 3D tissue models across representative embryo scStereo-seq samples, showing spatial distribution of label transferred cell types, mesh models for fat body or foregut/hindgut, and mesh models of the entire embryo. Due to high homology, some hindgut cells are annotated as foregut ones by label transfer. **(I)** Spatial distribution of cell bin *CytoTRACE* scores in representative fat body and foregut/hindgut models in **(H)**.

To characterize the activities of signaling pathways along tissue trajectories, we utilized 7 signaling pathway gene sets from *FlyphoneDB*^36^ and examined the expression dynamics of core pathway component genes across tissue developmental trajectories **(Figure S5A-B)**. Throughout the trajectories, we observed the up-regulation of multiple tissue-specific signaling pathways that are well-documented in the literature. The BMP signaling pathway, known for its integral role in ectoderm dorsal-ventral patterning (reviewed in Ref^37^), and in the regulation of neuromuscular junctions (NMJ, reviewed in Ref^38^), demonstrated the highest level of activity in early ectoderm and muscles. Meanwhile, the FGFR signaling pathway, which has been widely recognized for its role in trachea branching morphogenesis (reviewed in Ref^39^), showed maximum activity during the early stages of tracheal development. Lastly, the Hippo signaling pathway, well-established for its contribution to myogenesis (reviewed in Ref^40^), was most active in early muscle clusters **(Figure 3C)**. Thus, our multi-omics tissue developmental trajectories could serve as a systematic framework for exploring cell-cell communication networks.

### Spatiotemporal cell type succession during tissue development

Next, we aimed to visualize the spatiotemporal dynamics of the identified cell types along the multi-omics tissue development trajectories. Using the marker genes associated with cell types as a reference, we applied the label transfer method from *Seurat* to annotate scStereo-seq cell bins with the cell types identified in the multiomics tissue development trajectories. At the tissue level, the transferred labels demonstrated good agreement with manually annotated scStereo-seq cell bin clusters **(Figure 3D)**. Considering their relatively defined and regular morphology, we selected fat body and foregut/hindgut (both of ectodermal origin^41^) as models and aligned their cell types with embryo scStereo-seq samples. Within these tissues, the distribution of cell bins from label-transferred scStereo-seq and cells from scRNA-seq data exhibited a coherent pattern when plotted in the same UMAP space **(Figure 3E)**. Additionally, the top marker genes of each label-transferred cell type in scStereo-seq data were consistent with their counterparts in the integrated scRNA-seq and scATAC-seq data **(Figure S5C)**. These results suggested a precise mapping of cell types to their spatial locations in scStereo-seq data.

Within tissues, at the cell type level, the succession of different stages of cell types can be traced through their proportional changes over development **(Figure 3F)**. The spatial distribution of each cell type can be quantified by neighborhood enrichment, where a higher score indicates a greater level of spatial clustering **(Figure 3G)**. We observed significantly higher neighborhood enrichment in foregut/hindgut cell types compared to those in the fat body. When mapped to their spatial locations, different stages of foregut/hindgut cell types formed more aggregated clusters, while fat body cell types were more scattered and mixed **(Figure 3H)**. These observations suggested that these two tissues employ different cell differentiation strategies. In the fat body, differentiating cells are dispersed across the entire tissue, resulting in the mixing of cell types at different stages. In the foregut/hindgut, spatially defined “differentiation hubs” exist to continuously give rise to new cells, while cells outside the hubs do not contribute much to differentiation and proliferation. Consequently, cell types at different developmental stages occupy more distinct spatial locations. This hypothesis was further supported by the spatial distribution of cell bin *CytoTRACE* scores of scStereo-seq data. Cells with higher differentiation potential were more spatially aggregated in foregut/hindgut than in fat body **(Figure 3I)**. It is established that fat body cells originate from precursors arranged in segments that extend throughout the entire tissue^42^. This arrangement could account for the widespread dispersion of differentiating cells we observed here. On the other hand, the role of spatially clustered potential foregut/hindgut differentiation hubs might be associated with previously identified niches of digestive tract stem cells, where two defined groups of stem cells give rise to the adult foregut and hindgut, respectively^43^.

Therefore, through label transfer, we were able to map cell types along tissue development trajectories to their spatial locations in scStereo-seq samples, allowing us to track their spatiotemporal dynamics. In the following analyses, we extended this approach to more complex CNS and midgut cell types to uncover their dynamics during development.

### Transcription factor regulatory networks along tissue development trajectories

Transcription factors (TFs) play a pivotal role in orchestrating the proper formation and growth of tissues. To unravel the regulatory networks governed by TFs during tissue development and differentiation, we scrutinized the top marker genes of each cell type and investigated the enrichment of TF binding motifs in their promoter/TSS regions in scATAC-seq data. Motif enrichment analysis unveiled the regulatory TFs guiding the differentiation paths from each germ layer **(Table S7)**, encompassing both well-established cell type-specific regulators as well as potentially novel and uncharacterized ones.

Throughout the developmental trajectories, we pinpointed multiple well-characterized TFs that exhibited stage- and tissue-specific regulatory functions. We then traced the temporal dynamics of their regulatory activities along tissue development trajectories. Exemplary findings include motif enrichment of *GATAe* in Malpighian tubules^44^, *Rfx* in both PNS and CNS^45^, and *sage* in the salivary gland^46^ within the ectoderm. In the mesoderm, we identified motif enrichment of *Mef2* in somatic muscle^47^, *bin* in visceral muscle^48^, and *srp* in fat body^49^ and hemocytes^50^. The endoderm displayed motif enrichment of *CrebB* in early endoderm formation^51^, along with *fkh*, *GATAe*^52^, and other GATA family TFs (reviewed in Ref^53^) regulating late-stage endoderm specification **(Figure S6A-B)**. Additionally, we uncovered several previously uncharacterized TFs with potential spatiotemporally specific functions during embryogenesis. Notably, *CG34367*, a TF featuring a Homeobox (Hox) domain, exhibited significant and specific motif enrichment in the early primordium of all three germ layers, suggesting a ubiquitous role in early developmental regulation. Mammalian orthologs of *CG34367*, SHOX/SHOX2, are implicated in early organogenesis and their mutations are associated with genetic disorders including Turner syndrome^54,55^. The TF *crp*, ubiquitously expressed in multiple tissues and known for specifying terminal cells in tracheal tubes^56^, demonstrated potential regulatory functions in the mesodermal fat body and hemolymph, as indicated by our analysis. Moreover, we observed significant motif enrichment of *CG9727* and *CG12219* in nervous systems, *CG7368* in cardiac mesoderm, and *CG12236* and *CG4360* in early endoderm **(Figure 4A-B)**, indicating their specific functions in these tissues.

**Figure 4.**
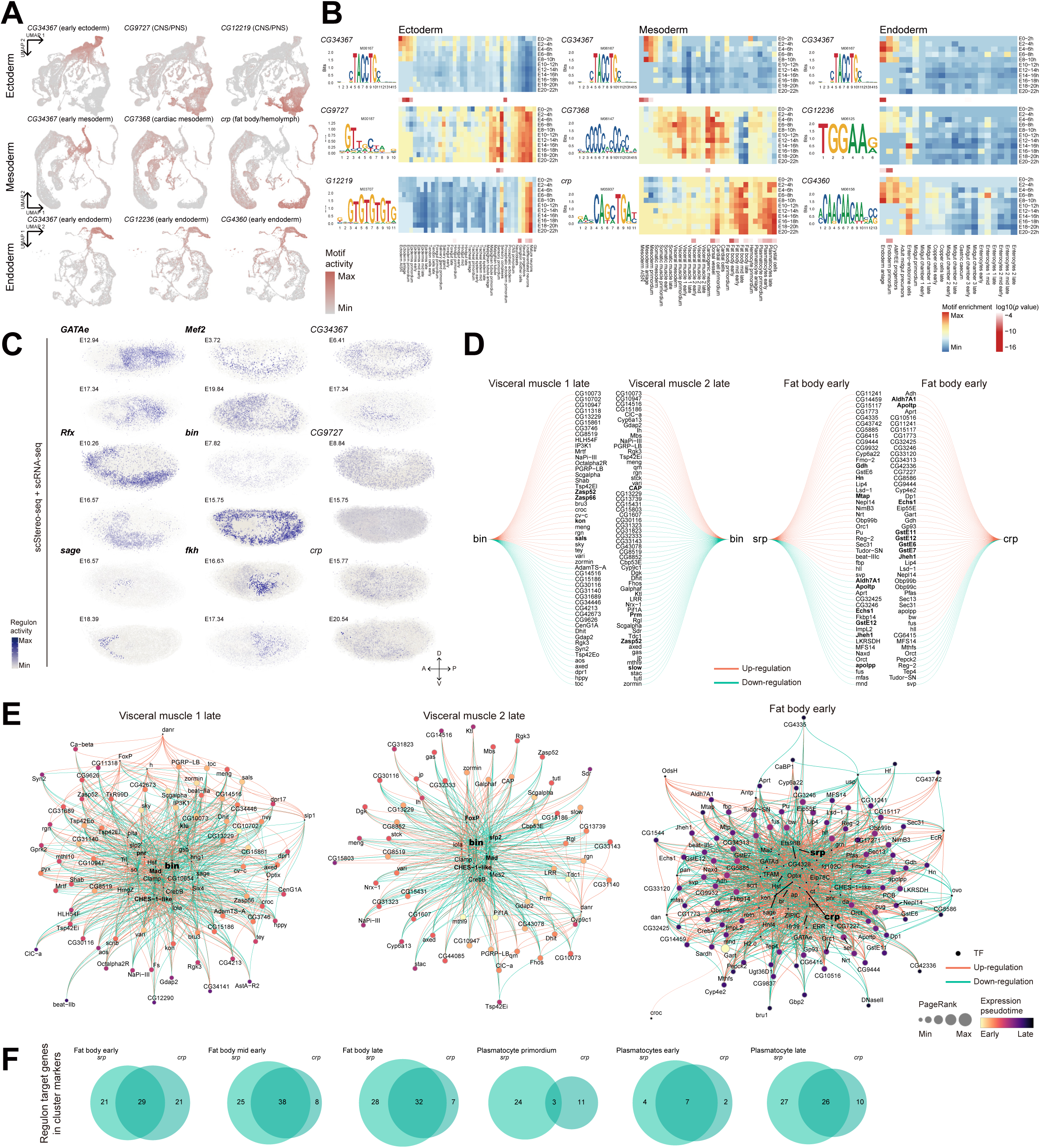
Transcription factor regulatory networks along multi-omics tissue development trajectories. **(A)** The same UMAP plots as Figure 2E but only show scATAC-seq cells, color coded with motif activities of less-characterized TFs. **(B)** TF motif enrichment along tissue development trajectories, showing less-characterized TF genes in **(A)**, their binding motifs (left), motif enrichment heatmap (upper right), and enrichment *p* value heatmap (lower right) across tissue types and developmental stages in cells from three germ layers in scATAC-seq data. **(C)** Visualization of *SCENIC* regulon activity of some of TFs in **(A)** in representative samples from integrated scStereo-seq and scRNA-seq data, projected along the Z-axis. Previously reported tissue-specific TFs are in bold. All scStereo-seq samples are shown in lateral or near-lateral view. A-P: anterior-posterior; D-V: dorsal-ventral. **(D)** *Pando* identified regulons of TF *bin* in visceral muscle 1 late and visceral muscle 2 late, and those of TF *srp* and *crp* in fat body early. Genes in bold are discussed in detail in the main text. **(E)** *Pando* identified GRNs of TFs (highlighted in bold) and cell types in **(D)**. Other TFs in bold are discussed in detail in the main text. **(F)** Venn diagrams showing overlap between target genes in regulons of *srp* and *crp* along developmental trajectories of fat body and plasmatocytes.

To further explore the spatial regulon activities of these TFs, we applied *SCENIC*^57^ to the integrated scStereo-seq and scRNA-seq data, revealing that the spatial patterns of regulon activities for both known and uncharacterized TFs were consistent with the motif enrichment analysis in terms of tissue specificity **(Figure 4C)**. The spatial expression patterns of these less-characterized TFs were also probed by BDGP *in situ* database and all of them exhibited weak signal or ubiquitous expression patterns in stages of their inferred functions **(Figure S6C)**. The lack of staining can be explained by poor probe efficiency or low expression levels of these TFs, while ubiquitously expressed TFs could perform regulatory functions in a tissue-specific manner. The ambiguous *in situ* staining results underscored the advantages of our multi-omics data in facilitating the elucidation of tissue-specific TF functions.

Subsequently, we employed *Pando*^58^ on the integrated scRNA-seq and scATAC-seq data to delve into the detailed regulons of identified TFs. Notably, visceral muscles exhibited segregation into two distinct groups in our multi-omics trajectories (dashed rectangles in **Figure 2E-F**, also see **Figure S6D and Data S2**). Upon scrutinizing the regulons of *bin* in these two groups, we observed that gene modules that *bin* regulated varied between them **(Figure 4D)**. In the visceral muscle 1 group, *bin* activity was positively correlated with expression levels of muscle assembly genes (*Zasp52*, *sals*, *Zasp66*, and *kon*). Conversely, in the visceral muscle 2 group, *bin* activity was negatively correlated with expression of genes with similar functions (*Prm*, *Zasp52*, *slow*, and *CAP*). Intriguingly, *bin* activity appeared to be partially opposite in regulating muscle structure assembly in these two groups. Supporting this result, the expression level of *Zasp52*, a core component of indirect flight muscles^59^, was significantly lower in scRNA-seq cells from visceral muscle 2 late cluster than those from visceral muscle 1 late cluster **(Figure S6E)**. It is known that *bin* is a cell fate determinant of transformation between somatic and visceral muscle through the BMP signaling pathway^48,60^. The contrasting effects *bin* exerted on some target genes in different visceral muscle cell groups may reflect its fine-tuning functions among muscle lineages. We further visualized the gene regulatory networks (GRNs) in which *bin* participated in these two groups **(Figure 4E)**. Inspection of GRNs in two visceral muscle groups uncovered several shared known muscle co-regulators of *bin*, such as *Mad*, a BMP pathway regulating TF functional at NMJ^61^, and *CHES-1-like*, also a BMP pathway regulator^62^. *bin* also co-regulated with different nervous system-related TFs in the two lineages, including *klu* and *pnr* in visceral muscle 1, and *slp2* and *FoxP* in visceral muscle 2. *klu* had reported functions in motor neurons^63^ while *FoxP* is important for motor coordination^64^. The organization of these GRNs highlighted the coordinated and cell type-specific co-regulation between nervous and muscle systems.

To further characterize the fat body- and hemolymph-specific regulon activities of *crp* identified above, we visualized its GRN in early fat body and discovered that *crp* co-regulated with *srp* **(Figure 4E)**. Upon inspecting their regulons, we found that *srp* activity was negatively correlated with lipid metabolism pathway genes (*Apoltp*, *Aldh7A1*, *apolpp*, *Jheh1*, and *Echs1*), while *crp* acted in a contrasting fashion. Additionally, *srp* positively regulated amino acid metabolism genes *Mtap*, *Hn*, and *Gdh*, while *crp* positively regulated glutathione metabolism genes *GstE6*, *GstE7*, *GstE11*, and *GstE12* **(Figure 4D)**. Target genes in the regulons of *srp* and *crp* largely overlapped in fat body and plasmatocytes, and this overlap increased along developmental trajectories of both tissues **(Figure 4F)**. The regulons of *srp* we identified were consistent with its role in inducing fat cell formation starting from early fat body development^49,65^ and *crp* is known to affect cell growth and tissue size control^56^. Our analysis suggested an increasingly coordinated role of *crp* and *srp* within the same GRN during fat body and plasmatocyte development.

In tracing tissue development trajectories, we successfully identified both previously reported and potential TFs, uncovering their tissue specificity and regulatory networks. It is worth noting that TFs and their binding motifs were linked based on *CIS-BP* database^66^. While the motifs we mentioned here were indeed enriched in specific cell types, it remained possible that their actual binding TFs differ from database inference, or there are additional unknown regulators that could bind these motifs. Algorithms like *Pando* used correlation between gene expression levels of TFs and their target genes to infer up- or down-regulation effects of these TFs, which could be susceptible to capture sensitivity of current single-cell sequencing techniques. Thus, additional experiment validation is required to elucidate these tissue-specific regulatory networks.

### Multi-omics dissection of gene regulation during embryonic CNS development

The *Drosophila* nervous system serves as a prominent model for investigating neuron development and functions. Thus, we examined the development of CNS from a multi-omics perspective based on our data. The subclustering results of the CNS scRNA-seq data identified most major CNS cell types, including neuroblasts (marked by *mira* and *wor*), ganglion mother cells (GMC, marked by *tap*), neural progenitors (marked by *insb* and *nerfin-1*), glioblasts (marked by *gcm* and *repo*), and various types of glial cells **(Table S4)**. The UMAP plot of CNS cells provided an intuitive representation of differentiation paths of neurons and glia **(Figure 5A)**. Subclustering scATAC-seq data also identified most of these CNS cell types **(Figure 5B)**. Integration of scRNA-seq and scATAC-seq data allowed detailed annotation of various mature neuron cell types by generating more distinct cell type specific markers **(Figure 5C-D and Table S4)**. In light of the significantly higher complexity of mature neuron cell types, we chose a higher resolution for their clustering and annotation. Each mature neuron cell type was annotated based on expression of neurotransmitters **(Figure 5D)**. Within mature neuron groups expressing the same neurotransmitters, cell subtypes were distinguished by a list of largely uncharacterized marker genes **(Figure 5E and Table S4)**. The complex trends of CNS cell differentiation are better visualized in 3D UMAP plots **(Data S3)**.

**Figure 5.**
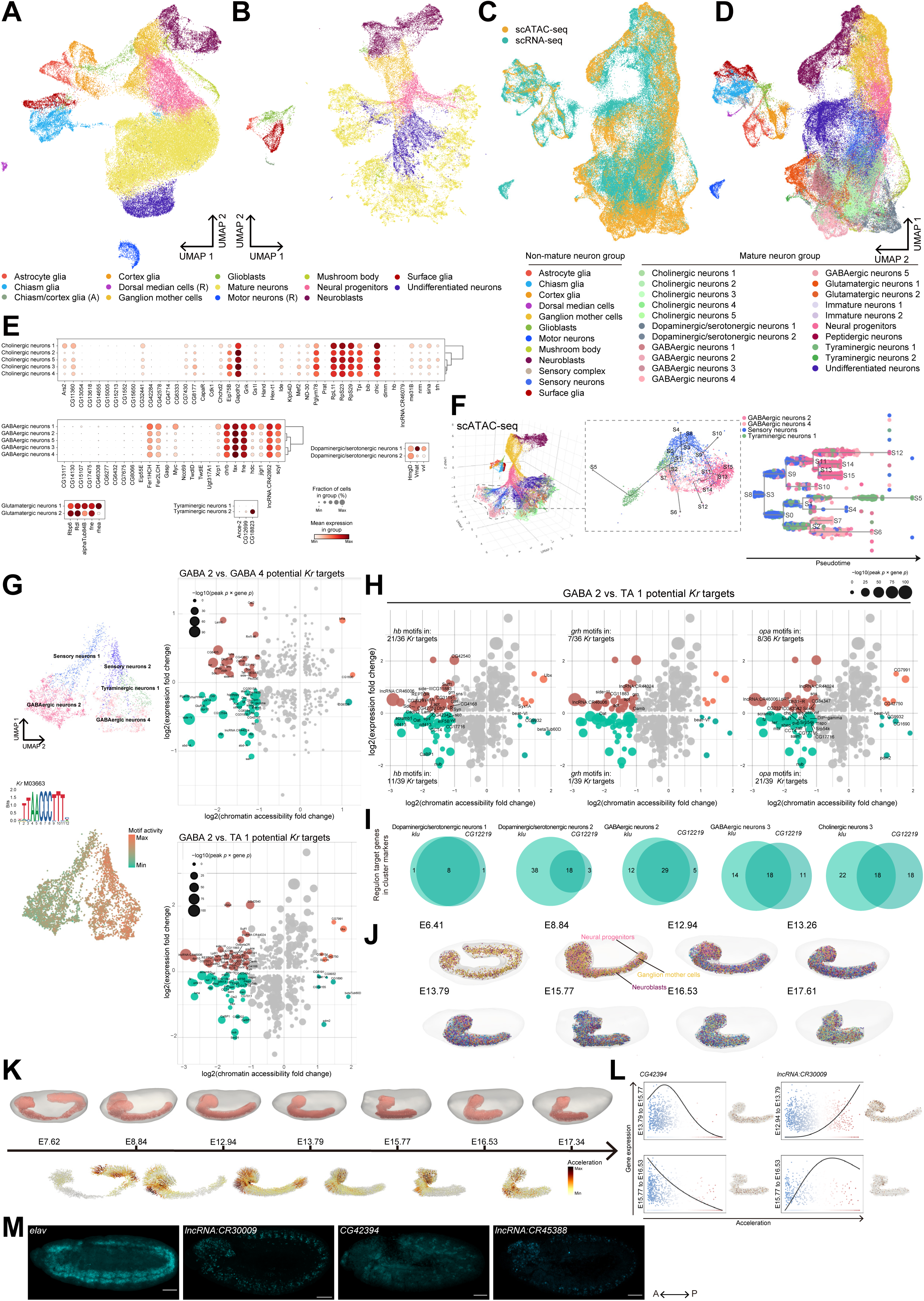
Gene regulation and morphometric dynamics in embryonic CNS. **(A)** UMAP plot showing subclustering and annotation of CNS cells from scRNA-seq data. **(B)** Same as **(A)**, but for scATAC-seq data. Annotations with (R) or (A) indicate clusters identified only in scRNA-seq data or only in scATAC-seq data, respectively. **(C)** Co-embedding of CNS cells from scRNA-seq and scATAC-seq data in the same UMAP plot. **(D)** Same as **(C)**, but re-clustered and re-annotated. **(E)** Bubble plots showing expression level and enrichment of top marker genes of mature neuron cell types in **(D)**. **(F)** Left: 3D UMAP plot of scATAC-seq data (also see **Data S3**) and differentiation trajectories of selected cell clusters, with S0 through S15 denoting branching points of differentiation. S8 was set as the origin of differentiation; right: subway map plot showing differentiation trajectories and branching points of the same cell clusters. Each dot represents one cell from scATAC-seq data subset. **(G)** Upper left: UMAP plot showing subclustering of sensory neurons and their differentiation paths; lower left: the same UMAP plot color coded with *Kr* motif activity. *Kr* has 5 known motifs with highly similar sequence compositions. The composition and activity of representative motif M03663 are shown; right: scatter plot showing the genes associated with DA peaks and DE genes, comparing GABAergic neurons 2 with GABAergic neurons 4, and GABAergic neurons 2 with tyraminergic neurons 1. Nearest genes of *Kr* binding motifs are labeled. The size of each dot corresponds to the product of *p* values for DA peaks and DE genes. **(H)** The same scatter plots as **(G)**, comparing GABAergic neurons 2 with tyraminergic neurons 1 and labeled with nearest genes of binding motifs of *hb*, *grh*, and *opa*. **(I)** Venn diagrams showing overlap between target genes in regulons of *klu* and *CG12219* among representative mature neuron cell types. **(J)** 3D CNS models across representative embryo scStereo-seq samples, showing spatial distribution of cell types, mesh models of CNS, and mesh models of the entire embryo. Cell type color codes are the same as **(D)**. **(K)** 3D models of CNS, CNS cell migration trajectories, and acceleration scores across 7 scStereo-seq samples of developmental age between 7 and 18 h. **(L)** General linear model-based correlation between acceleration scores and expression levels of *CG42394* and *lncRNA:CR30009* in transitions between representative scStereo-seq samples. Spatial gene expression patterns in CNS 3D models are shown on the right of each plot. **(M)** FISH validation in stage 11-16 embryos of gene candidates identified in CNS morphometric analysis. Representative images of pan-neuronal marker gene *elav* and candidate genes *CG42394*, *lncRNA: CR30009*, and *lncRNA:CR45388* are shown. All samples are shown in lateral view. A-P: anterior-posterior; D-V: dorsal-ventral. Scale bars = 50 μm.

Remarkably, mature neurons displayed a significantly more striking diversity in the UMAP plot derived from scATAC-seq data compared to that from scRNA-seq data **(Figure 5A-B and Data S3)**. This observation suggested the possibility that mature neurons appear similar in their transcriptomic profiles during late embryogenesis, but various types of neurons are under highly distinct epigenetic regulations, likely in preparation for more complex neural differentiation during larval stages. Notably, co-embedding of scRNA-seq and scATAC-seq CNS cells in the same UMAP plot showed that mature neurons in scATAC-seq data displayed an overall distribution shift from those in scRNA-seq data **(Figure 5C and S7A)**. This shift was not observed in non-mature neuron cell types **(Figure S7A)**. Similarly, we also noted a temporal mismatch in the distribution of mature neuron subtypes between scATAC-seq and scRNA-seq data **(Figure S7B)**. This further reflected the potential divergence between transcriptomic and epigenomic profiles among mature neurons.

### Differentiation trajectories of mature neurons revealed by scATAC-seq

To dissect the epigenetic regulation of mature neuron and identify potential regulators of cell subtype differentiation, we explored the 3D UMAP plot of scATAC-seq data and focused on a cell subset, in which a cell cluster expressing sensory neuron markers (e.g., *ct*, *lov*, and *robo3*) appeared to differentiate into three mature neuron clusters: GABAergic (GABA) neurons 2 & 4 and tyraminergic (TA) neurons 1 **(Figure 5F and Data S3)**. We employed *STREAM*^67^ to map the differentiation trajectories of these cell clusters, which were then projected onto the 3D UMAP space. This enabled us to identify the branching events within the differentiation trajectories **(Figure 5F)**. Leveraging these trajectories, we further subclustered sensory neurons into two distinct groups according to their chromatin accessibility and differentiation outcomes **(Figure 5G)**. Focusing on the top DA peaks between sensory 1 & 2, as well as those among GABA 2, GABA 4, and TA 1, we conducted a TF motif enrichment analysis. This revealed *Kr* as a principal regulator of this differentiation process, with significant motif activity contrast between the differentiation branches **(Figure 5G and Figure S7C)**. *Kr* is a well-established transcription repressor and temporal determinant of neuron fate^68,69^. The gene *Kr* itself exhibited significantly higher chromatin accessibility and expression level in GABA 2 compared to GABA 4/TA 1 **(Figure S7D)**, suggesting a more active regulatory activity in GABA 2. On the contrary, the binding motifs of *Kr* are significantly less enriched in GABA 2 and most of its potential target genes (nearest genes of *Kr* binding motifs) displayed reduced chromatin accessibility and expression level in GABA 2 compared to GABA 4/TA 1 **(Figure 5G)**. This observation supports a working model based on previous knowledge that *Kr* performs its transcription repressor functions through local quenching of transcription activators^70,71^, likely through closing up the proximal chromatin. We carefully examined the genes that showed significant changes in both chromatin accessibility and expression level among the neuron subtypes. Pathway enrichment of these genes showed that these *Kr* target genes were functionally enriched in processes including axon guidance and glycosylation **(Figure S7E)**.

While most potential targets of *Kr* showed decreased chromatin accessibility, their expression changes varied across target genes. This variability could be due to the impact from transcriptional co-factors of *Kr*. To identify co-regulators that influenced the expression levels of genes repressed by *Kr*, we conducted a motif enrichment analysis within *Kr* peaks **(Figure S7F)**. This revealed several previously characterized neuron differentiation regulators, including *hb*, *grh*, and *opa*, through the comparison between GABA 2 and TA 1. **(Figure 5H)**. It is well established that the sequential activities of *hb*, *Kr*, and *grh* determine the temporal fate of several neuroblast lineages during differentiation (reviewed in Ref^72^). *opa* is previously reported as a regulator of *Kr* activity during early embryogenesis^73^, as well as a regulator of temporal patterning of neural progenitors that acts in coordination with *grh*^74^. Our observations suggested that the synergy of these regulators persist in more differentiated neuron subtypes. Both *hb* and *grh* are known to function as either transcription activators or repressors^75–77^. In the differentiation process we investigated here, the motif activities of *hb* and *grh* were mostly in up-regulated *Kr* target genes, even in the presence of repressive effect of *Kr*. Conversely, the motif activities of *opa* were enriched in down-regulated *Kr* target genes **(Figure 5H)**. We then examined the peaks around the binding motifs of these co-regulators in the chromosomal regions of their mutual target genes using the scATAC-seq data. We observed a dramatic overall increase in chromatin accessibility along the differentiation track from sensory 2 to GABA 4/TA 1, compared to the subtle changes during the transition from sensory 1 to GABA 2. This seems to be a general phenomenon regardless of the expression change between GABA 2 and GABA 4/TA 1, suggesting diverse and complex regulatory consequences depending on the cooperating TFs and the targets. This was further supported by the long distance of *Kr*/co-regulator peaks from the TSS of the nearest genes, which could be a few kb in length and mostly downstream of gene targets. As examples, the peaks and chromosomal regions of two genes with the most significant expression level changes, *side-III* (potentially co-regulated by *Kr*, *hb*, and *grh*) and *fz* (potentially co-regulated by *Kr* and *opa*), are plotted to demonstrate their coordinated regulatory roles outside of promoter regions (**Figure S7G**).

Overall, we observed a high clustering resolution in scATAC-seq data when characterizing mature neuron subtypes, which was able to facilitate the discovery of transcription regulators and their co-factors that govern the refined developmental trajectories.

### Mutual and diverse GRNs among CNS cell subtypes

In pursuit of potential regulators of the diverse neuron cell types, we conducted motif enrichment analysis and pinpointed TF regulators across various stages of neural development **(Table S7)**. Among these, previously reported TFs, such as *seq*, governing dendrite and axon outgrowth^78^, exhibited the highest motif activity in neuron progenitors. Additionally, *klu*, known to specify the identity of a specific group of neuroblasts^79^, displayed sustained activity in several types of mature neurons **(Figure S8A)**. Our analysis also revealed cell type-specific activity for several less characterized TFs, including *BEAF-32*^80^ and above-mentioned potential nervous system-specific regulator *CG12219* **(Figure S8A)**. *Pando* visualization of their regulons showed that in neuroblasts, *BEAF-32* and *seq* co-regulated multiple cell cycle regulator genes (e.g., *PolE2*, *fzy*, *mad2*, and *Mcm2*) and neuroblast determinants (e.g., *mira* and *CycE*) **(Figure S8B-C)**. In mature neuron clusters, *CG12219* and *klu* displayed similar activity patterns across cell types **(Figure S8A)** and their regulons largely coincided in mature neuron cell types **(Figure 5I)**. For example, in dopaminergic/serotonergic neurons 2, *CG12219* and *klu* co-regulated the same group of signal transduction genes (e.g., *Pkc53E*, *Oct1R*, *Syngr*, *Sytα*, and *CG34393*) in the same GRN; In cholinergic neurons 3, *klu* and *CG12219* co-regulated glucose metabolism (e.g., *Pgi* and *Pgm1*) gene groups in the same GRN **(Figure S8B-C)**.

Our findings strongly supported the existence of cell subtype-specific regulons for the same TF, as well as the cooperative actions of different TFs that are finely tuned for neuronal differentiation. These regulatory mechanisms may play a role in orchestrating the precise development of neurons.

### Refined spatiotemporal CNS cell subtypes during embryogenesis

We subsequently applied label transfer to project identified CNS cell types onto scStereo-seq samples **(Figure 5J)**. As expected, there was a discernible shift in cell count fraction from undifferentiated neuroblasts and GMCs to differentiated neuron and glia cell types from early to late-stage samples **(Figure S8D)**. Co-embedding embryo scStereo-seq data with scRNA-seq data in the same UMAP space demonstrated high coherence **(Figure S8E)**. Among the transferred cell types, as expected from their anatomical distribution, neuroblasts and glia cell types exhibited the highest level of spatial aggregation, whereas mature neurons were largely dispersed **(Figure S8F)**. Upon inspecting their spatial loci, the distribution of CNS cell types aligned well with their stratified anatomical structures in early-stage samples, with less differentiated cell types occupying the outer layers of the CNS and more differentiated ones in the inner layers **(Figure 5J)**. These findings supported the precision of the label transfer method in identifying CNS cell subtypes in scStereo-seq samples, thereby facilitating the exploration of neuron functions within their spatial context.

### Gene expression dynamics during CNS morphometric changes

The *Drosophila* CNS undergoes profound morphological transformations throughout embryogenesis, influenced by intrinsic factors such as cell proliferation and differentiation, as well as external cues like inter-organ communication (reviewed in Refs^72,81,82^). Leveraging our 3D spatial transcriptomes generated with scStereo-seq, we delved into transcriptomic dynamics during morphogenesis by simultaneously tracking changes in tissue morphology and gene expression. Employing morphometric analysis from the *Spateo* package, we were able to align the 3D point-cloud models of two time points in spatial coordinates. Subsequently, we linked cell bins between the two samples based on spatial adjacency and transcriptomic similarity **(Figure S9A)**. This enabled the generation of 3D vectors, concurrently characterizing cell migration paths and transcriptomic changes over continuous developmental stages **(Figure 5K)**. Finally, we computed morphometric parameters describing cell migration paths and correlated them with transcriptomic changes.

We characterized the morphometric changes in the CNS across 3D models of seven scStereo-seq samples, spanning developmental ages from 7 to 18 h. These changes were represented by parameters such as the acceleration of cell migration (proportional to the distance cells migrated given the same migration time between two samples) **(Figure 5K and Movie S1)**, curvature (bending of cell migration paths) **(Figure S9B)**, curl (rotation of paths) **(Figure S9C)**, and torsion (curve twisting of paths) **(Figure S9D)**. Throughout CNS development, we observed a shift in regions with the highest acceleration from the posterior end of the ventral nerve cord (VNC) to the anterior end of the brain **(Figure 5K)**. The decline in acceleration and curl scores in the VNC was likely linked to the completion of germ band retraction, indicating that the shortening of the VNC during early development primarily relied on the migration of posterior cells toward the anterior end. Conversely, the increase in acceleration and curl scores in the anterior brain region might reflect active cell organization in brain lobes during late embryogenesis **(Figure 5K and Figure S9C)**. As anticipated from CNS morphology, regions with the highest curvature and curl scores concentrated around the curved joint between the VNC and the brain **(Figure S9B-C)**.

The morphometric analysis yielded a set of genes exhibiting spatiotemporal expression changes relevant to CNS morphometric dynamics **(Table S8)**. Gene ontology (GO) enrichment revealed that genes linked to CNS morphometric changes were highly enriched in cell fate specification and pattern formation **(Figure S9E and Table S8)**. Additionally, gene group enrichment analysis highlighted the significance of Hox family transcription factors, such as *Antp*, *Ubx*, *abd-A*, and *Abd-B*, consistent with their critical roles in specifying CNS patterns and segment identity^83^. The spatial expression patterns of these Hox family genes in our 3D CNS models aligned with BDGP *in situ* results **(Figure S9F)**. Notably, the expression levels of these Hox genes were mostly negatively correlated with acceleration scores across developmental stages **(Figure S9G)**, suggesting that their expression is associated with the inhibition of CNS cell migration. It is reported that Hox genes’ roles include repressing neuroblast formation and entry into neuroblast quiescence in embryonic CNS (reviewed in Ref^84^). It is possible that Hox genes inhibit CNS cell migration through repression of neuroblast differentiation.

Associations were also observed between CNS morphometric scores and known CNS development regulators (e.g., *mira*, *tll*, and *toy*) as well as several uncharacterized factors **(Table S8)**. For example, the expression level of *CG42394* was negatively correlated with acceleration, while that of *lncRNA:CR30009* displayed a positive correlation **(Figure 5L)**. We validated the CNS-specific expression of these potential regulators with FISH **(Figure 5M)**. Notably, this list includes multiple long non-coding RNA (lncRNA) genes besides *lncRNA:CR30009*, which was previously reported to be enriched in glia and co-localize with the glia marker gene *repo*^85^. Examining these lncRNA genes in our scRNA-seq data, we observed that the expression of *lncRNA:CR30009* and *lncRNA:CR45388* showed the highest correlation with neuroblast and glioblast marker genes **(Figure S9H)**. These observations implied that the two lncRNA genes may influence CNS morphometric changes through the regulation of neuroblasts. Therefore, by conducting morphometric analysis of the CNS, we were able to identify both known and potential regulators of CNS cell migration.

### Cell type and functional diversity of developing midgut

The *Drosophila* midgut, serving as the functional equivalent of the mammalian small intestine, fulfills versatile roles in food digestion, nutrient uptake, immunity, and endocrine regulation. The diverse functions of the midgut are carried out by distinct types of cells and the regions they form (reviewed in Refs^86,87^). Nevertheless, the timing of differentiation of these cell types remained elusive. Our prior investigations indicated that certain functional cell types began to emerge during late embryogenesis^13^.

Here, we delved deeper into the diversity of midgut cell types using our multi-omics data. The clustering resolution of our scRNA-seq data was adequate for distinguishing various midgut cell types. Consequently, we concentrated on the scRNA-seq data, combined endoderm and midgut cell clusters, and conducted high-resolution subclustering and annotation **(Figure 6A)**. The UMAP plot portrayed a multitude of intestinal cell types throughout the developmental and differentiation stages of the midgut **(Figure 6A-B)**. These included endoderm (marked by Notch signaling pathway genes *E(spl)m4-BFM* and *Brd*), adult midgut progenitors (AMPs, marked by *esg*)^26^, 6 types of entero-endocrine cells [EEs, marked by *pros* and distinguished by specific expression endocrine genes **(Figure 6C)**], and 6 types of enterocytes (ECs, marked and distinguished by digestive enzyme and metabolism-related genes)^88^. Cell clusters in transitional states between midgut primordium and functional ECs were denoted as “midgut chambers”, with each cluster distinguished by its top markers.

**Figure 6.**
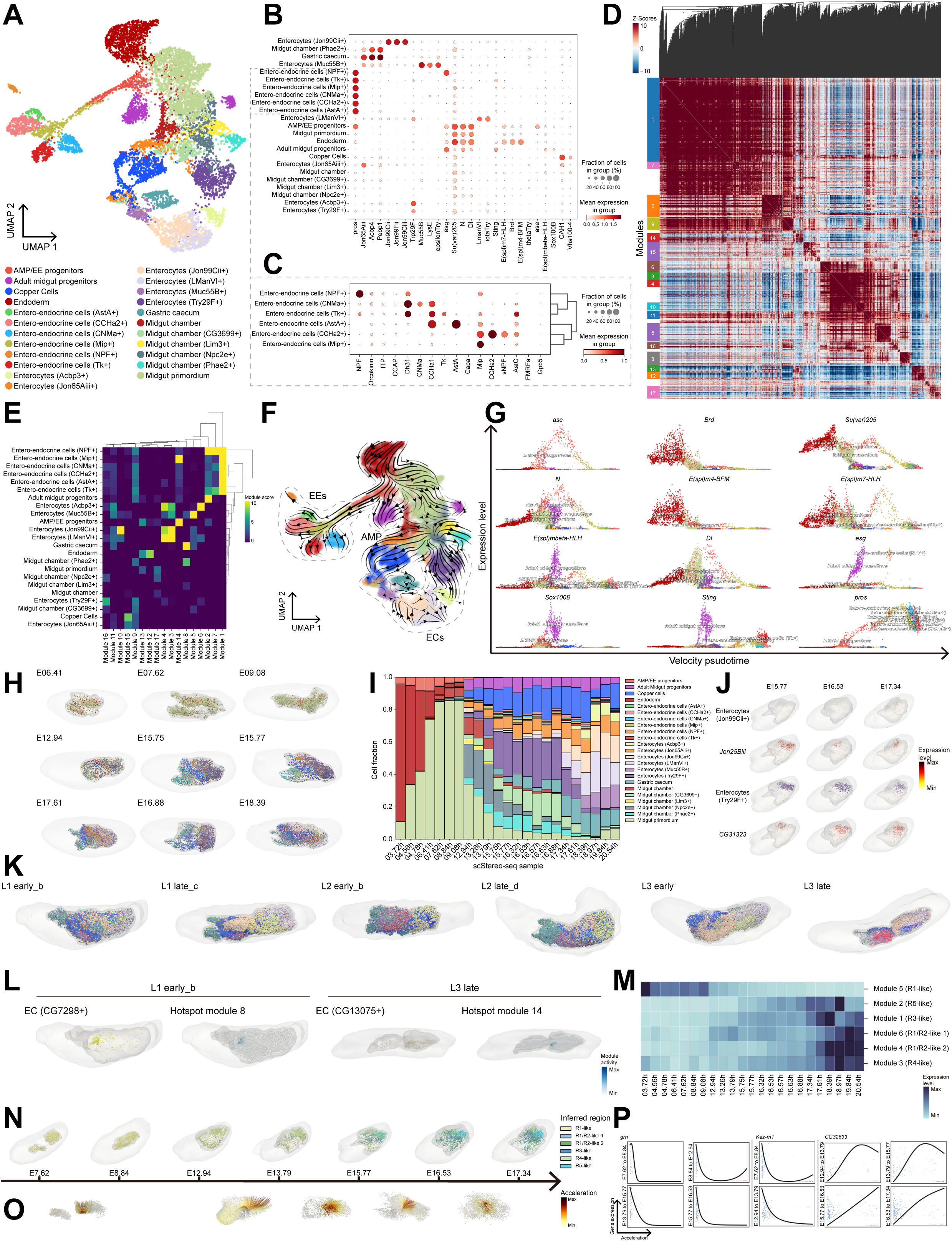
Cell type diversity and functional regionalization in midgut. **(A)** UMAP plot showing subclustering and annotation of endoderm and midgut cells from scRNA-seq data, derived from *Dynamo* analysis. **(B)** Bubble plot showing expression level and enrichment of top marker genes of cell types in **(A)**. **(C)** Same as **(B)** but within entero-endocrine cells. **(D)** Heat map showing correlation of functional gene modules identified by *Hotspot* in scRNA-seq data. Each row and each column represent a module marker gene, and *Z*-score indicates their correlation. **(E)** Heat map showing enrichment and clustering of *Hotspot* identified gene modules from **(D)** in midgut cell types in scRNA-seq data. **(F)** RNA velocity flow projected in UMAP space in **(A)**. Cell type color codes are the same as **(A)**. Dashed lines mark clusters representing adult midgut progenitors (AMPs), entero-endocrine cells (EEs), and enterocytes (ECs) discussed in the main text. **(G)** Dot plots showing relationship between velocity derived pseudotime and expression levels of genes of interest during differentiation of AMPs and EEs. Each dot represents one cell from midgut scRNA-seq data. **(H)** 3D midgut models across representative embryo scStereo-seq samples, showing spatial distribution of cell types, mesh models of midgut, and mesh models of the entire embryo. Cell type color codes are the same as **(A)**. **(I)** Bar plot showing cell type composition of midgut in scStereo-seq samples. Cell types are label transferred from scRNA-seq data. **(J)** Same as **(H)** but showing spatial distribution of copper cells, EC (*Jon99Cii+*), and EC (*Try29F*+) and their cell type marker genes in representative scStereo-seq samples. **(K)** Same as **(H)** but for larva scStereo-seq samples. Cell type color codes are the same as **Figure S11D**. Samples are not on the same scale. **(L)** 3D midgut models of L1 early and L3 late scStereo-seq samples, showing spatial distribution of representative ECs and their corresponding functional gene modules. **(M)** Heat map showing expression level of region-related gene modules across scStereo-seq samples. **(N)** Same as **(H)** but showing spatial distribution of inferred “adult midgut” regions. **(O)** 3D midgut cell migration trajectories and acceleration scores across 7 scStereo-seq samples of developmental age between 7 and 18 h. Sample viewpoints are different from **(N)** for better visualization of trajectories. **(P)** General linear model-based correlation between acceleration scores and expression levels of *grn*, *Kaz-m1*, and *CG32633* in transitions between representative scStereo-seq samples.

#### Differentiation and molecular characteristics of embryonic midgut cell typesV

We further investigated the molecular markers of distinct midgut cell types. Pathway enrichment analysis of cluster marker genes revealed that embryonic midgut cells are regulated by distinct signaling pathways, reflecting their versatile functions. Of note, the Notch signaling pathway was enriched in early endoderm, AMP/EE progenitors, and AMPs clusters, consistent with previous reports^27,89^. Additionally, the Wnt signaling pathway was enriched in multiple EE clusters^90^ **(Figure S10A)**. Interestingly, AMPs and several EE clusters showed high enrichment in pathways related to autophagy and apoptosis **(Figure S10A)**. Indeed, the expression of autophagy-related genes *Atg101* and *Atg9*^91^, as well as apoptosis-related genes *chrb* and *scyl*^92^ are specifically enriched in AMPs and EEs within midgut **(Figure S10B)**. It is possible that AMPs and EEs employ cell death-related mechanisms to maintain homeostasis during embryogenesis. To further characterize the functions of these diverse midgut cells, we used *Hotspot*^93^ to identify 17 gene modules from midgut scRNA-seq data **(Figure 6D)**, each with distinct functional GO and cell type-specific enrichment **(Figure 6E)**. For example, module 1, enriched in neuropeptide signaling pathways, was concentrated in all 6 EE clusters, aligning with their role in sensing stimuli and secreting neural signals for physiological regulation; Module 6, enriched in Cytochrome P450 family enzymes, was concentrated in EC (*Acbp3+*), suggesting a significant role in metabolism; Module 14, functionally enriched in genes regulating nervous system development and the Notch signaling pathway, was concentrated in AMPs and EE (*Mip+*) **(Figure S10C)**.

It is established that during metamorphosis, larval midgut cells undergo apoptosis, and adult midgut cells arise from AMPs to reconstitute the adult midgut^89,94^. Interestingly, in the UMAP plot, the differentiation trajectory of endoderm cells branched early on towards adult cell types (AMP/EE) and larval ones, which implied that the fates of these cell types were predetermined upon their differentiation from the endoderm primordium **(Figure 6A)**. We observed that AMPs and EEs originated from the same cluster of cells in the UMAP plot, marked by the expression of *esg* and *pros*. This is in line with previous reports indicating that AMPs and EEs derive from the same group of midgut cells^27^, which we denoted as “AMP/EE progenitors” in our data **(Figure 2E-F)**. To further track the kinetics of cell type emergence, we employed *Dynamo*^95^ to illustrate the transcriptomic vector fields of midgut development. *Dynamo* analysis revealed that the kinetics of cell state changes supported the notion that AMPs and EEs derived from the same group of progenitors **(Figure 6F)**. As previously reported, the Notch signaling pathway extensively participated in this differentiation process^27,89^, along with stem cell differentiation factors such as *esg*, *pros*, *ase*, and *Sox100B*^88^. Notably, we observed highly specific dynamics of the innate immune signaling gene *Sting*^96^ in AMPs, suggesting its role in the specification of AMPs **(Figure 6G)**. *Dynamo* analysis also facilitated the tracing of EC formation from their precursors in midgut chambers **(Figure 6F)** and suggested cell type-specific markers for their specification **(Figure S10D)**.

In summary, we categorized and examined the variety of cell types, each with unique functions, present in the embryonic midgut. This allowed us to reveal the differentiation trajectories of AMPs and EEs, as well as identify potential regulatory processes that govern their development and maintenance.

#### Spatial distribution of midgut cell types from embryonic to pupal stages

Next, we sought to map the midgut cell types we identified to their spatial locations. Using the cell type marker genes from scRNA-seq data as a reference, we located their counterparts in scStereo-seq data through label transfer **(Figure 6H)**. Co-embedding scRNA-seq and label transferred scStereo-seq data in the same UMAP space demonstrated high coherence **(Figure S10E)**. The top marker genes of label transferred scStereo-seq cell bins were also consistent with scRNA-seq cells **(Figure S10F)**, indicating precise mapping of cell types to their spatial locations. In the label transferred scStereo-seq 3D models, we observed the dynamics of changes in cell fraction throughout development, reflecting the different timings of emergence of these cell types. For example, EC (*Try29F+*) appeared around 13 h of development, while EC (*Acbp3+*) did not form until around 17 h **(Figure 6I)**. Neighborhood enrichment analysis suggested that although most cell types were sparse in their spatial distribution, EC (*Jon99Cii+*) and EC (*Try29F+*) were more aggregated compared to other cell types **(Figure S10G)**. Indeed, these cell types and the expression patterns of their marker genes occupied distinct spatial loci consistently across embryo scStereo-seq samples **(Figure 6J)**. Therefore, mapping cell types to our scStereo-seq data enabled the tracing of embryonic midgut cell type distribution within their spatiotemporal context. This approach provides a comprehensive understanding of how different cell types are spatially and temporally organized during embryonic midgut development.

We subsequently endeavored to identify midgut cell types in the larva and pupa scStereo-seq samples. We performed subclustering for larval and pupal midgut cell bins and used label transfer results as a reference for annotation. Given the substantial developmental changes and technical differences, we opted to use label transferred and re-annotated embryo scStereo-seq cell bins as the reference for label transfer of larva and pupa scStereo-seq samples, rather than embryo scRNA-seq cells. Cell clusters that demonstrated low confidence in label transfer were annotated as new larval or pupal cell types based on top marker genes **(Figure S11A-C and Table S4)**.

Compared to embryos, larvae displayed a more diverse array of intestinal epithelial cell types over development **(Figure 6K and Figure S11D)**. Notably, different ECs were densely clustered along the anterior-posterior axis of the midgut, as observed in the 3D models of their distribution **(Figure 6K)**. To understand their roles, we carried out *Hotspot* gene module analysis on each larva scStereo-seq sample. To fully leverage our scStereo-seq data, we took spatial location of cell bins into consideration during identification of gene modules. Examining the functional enrichment of gene modules, we noticed that various cell types performed unique functions, some of which were similar to those in the embryonic midgut, while others were new and specific to the larval midgut. For instance, the gene module concentrated in EC (*CG7298*+) in L1 early implied functional enrichment of chitin formation, suggesting that cell type specialized midgut chitinization (reviewed in Ref^97^) commenced in the early larval stages; the gene module concentrated in EC (*CG13075*+) in L3 late indicated functional enrichment of apoptosis and pattern specification, possibly related with midgut remodeling during late larval stages **(Figure 6L and Figure S11E)**.

During the L3 stage, substantial changes occurred among midgut cell types. The anterior gastric caecum and the posterior EC (*Acbp3+*) in the midgut contracted and decreased in number in L3 early sample, eventually vanishing completely in L3 late sample **(Figure 6K and Figure S11D)**. This suggested that significant remodeling and reorganization of the midgut takes place during the L3 stage, which coincided with the previously established timing of midgut cell death in this region before metamorphosis^98^. Accompanying this process, we observed a marked and widespread shift in the gene expression profiles of various cell types within the midgut, characterized by a considerable increase in the expression of ribosomal and mitochondrial genes and their representation in marker genes **(Table S4)**.

We subsequently examined the morphology of the midgut and the marker genes within the pupal midgut subclusters and identified a notable cluster of midgut cells that, rather than expressing digestion-related genes, exhibited strong expression of ecdysone-responsive genes, such as *Eig71Ek*, *Eig71Ea*, and *Edg78E* **(Table S2)**. These cells, which we designated as “midgut outer” in manual annotation, formed a sheath-like structure that enveloped the rest of midgut cells, which we labeled as “midgut inner” **(Figure S11F and Data S1)**. These structures, which receded after the P24 stage and re-emerged at the P72 stage, bore a strong resemblance to the yellow body and its surrounding midgut epithelium into which the midgut delaminates during metamorphosis^26^ (reviewed in Ref^99^).

In conclusion, the use of label transfer-assisted spatial mapping and annotation unveiled spatially restricted and cell type-specific functions of larval and pupal midgut cell types. The spatial transcriptomic data from our pupa scStereo-seq samples also offered valuable resources for studying the regulation of midgut morphogenesis.

#### Emergence and layout of embryonic midgut regions

The adult *Drosophila* midgut is conventionally divided into five regions (hereafter termed R1∼R5) based on morphological constraints, with each region performing distinct functions^100,101^. While functional regionalization of the midgut has been extensively studied at larva and adult stages, the precise timing of subregion emergence during embryonic development remains to be elucidated. In our previous study utilizing 3D spatial transcriptomic models, we observed the emergence of subregions with distinct digestive functions during late embryogenesis^13^.

Here, we further characterized the process of embryonic midgut regionalization. Referring to regional marker genes summarized in Ref^101^, we first identified 6 gene modules from adult midgut regional marker genes with *Hotspot* **(Figure S12A)** and established their correlation with expression profiles of adult midgut regions. Among them, modules 4 and 6 display similar correlation with both R1 and R2, so we denoted the two regions they corresponded to R1/R2-like 1 & 2, respectively **(Figure S12B)**. With these regional markers as references, we identified cell groups exhibiting transcriptomic similarity to adult R1 to R5 in scStereo-seq midgut cell bins, which we termed R1-like to R5-like. Each regional cell groups displayed distinct marker gene expression **(Figure S12C)** and increasing levels of spatial clustering over development, suggesting that they occupied distinct areas in the midgut **(Figure S12D)**. This allowed us to investigate the timing of their appearance and trace the spatial distribution of these regions. Upon inspecting the expression of gene modules in scStereo-seq samples, we noted that the expression of R1-like gene modules initiated at the very early stages of endoderm development and gradually declined over time. Modules corresponding to other regions began to actively express around 13 h of embryogenesis **(Figure 6M)**. Simultaneously, the spatial distribution of regions started to crystallize around the same time point, mirroring the spatial order as observed in the adult midgut (R1 to R5 from anterior to posterior) **(Figure 6N)**. This suggested that although midgut underwent lysis and reformation during metamorphosis, its regional organization was already patterned during embryogenesis.

To profile the biological functions each region undertook, we examined GO enrichment of marker genes for the identified embryonic midgut regions. The R1-like region is functionally enriched in protein metabolism; the R1/R2-like regions are functionally enriched in fatty acid metabolism; the R3-like region is functionally enriched in ion transport and pH regulation, consistent with the acidic nature of this region^102^; the R4-like region is functionally enriched in stimuli sensing, proteolysis, and nucleic acid metabolism; the R5-like region is functionally enriched in metal ion homeostasis **(Figure S12E)**. These functions aligned well with their counterparts in adult midgut regions^100,101^. We analyzed the cell type composition of each region and observed that over development, each region acquired its major cell types. For instance, R1/R2-like 2 mainly composed of EC (*Muc55B*+) and R3-like mainly composed of copper cells and EC (*Jon65Aiii*+) **(Figure S12F)**.

Together, our scStereo-seq data demonstrated distinctive cell compositions in the embryonic midgut regions, which determined the spatially localized sub-organ functions maintained through adulthood.

#### Morphometric regulators during embryonic midgut development

The *Drosophila* midgut experiences significant morphological transformations during development. It forms from the fusion of two separate rudiments at the anterior and posterior ends of the embryo, evolves into a closed chamber, and eventually establishes a highly convoluted tube-like morphology (reviewed in Ref^103^). We used *Spateo* to model the morphological changes during the fusion of the anterior and posterior midgut around stage 12 (∼8 h of development) and the convolution of midgut tubes during late embryogenesis **(Figure 6O and Movie S2)**. Morphometric analysis of cell migration modeled the fusion of early midgut and the torsion of late midgut, which revealed an association between midgut cell acceleration and multiple previously reported morphogenesis regulators across stages, including the GATA family TF *grn*, which is known to regulate the process of midgut fusion^104^, and Notch signaling pathway component *Kaz-m1*, which displays a highly restricted expression pattern at the fusion site of the midgut and has a potential regulatory role^105^ **(Table S8)**. The expression levels of both factors were negatively correlated with acceleration scores, suggesting that they were associated with inhibition of midgut cell migration **(Figure 6P)**. Starting from E15.77, gastric caecum-specific marker genes *Acbp4* and *Pebp1* ranked top in genes associated with all aspects of morphometric changes **(Table S8)**, in line with the timing of gastric caecum extrusion and formation from the midgut chamber (∼15 h of development)^41^. In addition, multiple Acyl-CoA binding protein (Acbp) family genes demonstrated high correlation with morphometric scores in late-stage midgut, which aligned with their known function of linking nutrient sensing and shaping tissue plasticity^106^. We also observed several potential regulators or effectors of midgut morphological changes, such as *CG32633*, which consistently displayed positive correlation with cell migration acceleration across samples **(Figure 6P)**. Thus, morphometric analysis provided clues for identifying potential regulators during the complex morphogenesis process of embryonic midgut.

## DISCUSSION

After our initial proof-of-principle application of Stereo-seq on *Drosophila*, we present here a single-cell 3D spatiotemporal multi-omics atlas spanning developmental lifespan of *Drosophila* from embryogenesis to metamorphosis. The current study builds upon our previous work by enhancing the Stereo-seq spatial transcriptomic dataset in several ways. Firstly, the sample collection window was expanded to include development from embryo to pupa. While ISH databases like BDGP and Fly-FISH have extensively probed spatial gene expression patterns in embryonic stages, there are still missing genes in these databases. Additionally, similar systematic databases are notably absent for the larval and pupal stages. Our scStereo-seq data effectively encapsulated the spatial gene expression patterns, unveiling the spatiotemporal gene expression dynamics for a list of over 300 genes in embryos, previously uncharted in ISH databases. Our data also serves as a valuable asset for delving into the spatial gene expression patterns in larvae and pupae. Secondly, our previous work using merged bins of a predetermined number of DNBs as units of analysis (e.g., bin 20 × 20 recognized 400 DNBs as a single “cell”) did not accurately capture the transcriptomes of individual cells. Here, we incorporated imaging data from nucleus staining with Stereo-seq to enable cell segmentation and established single-cell spatial transcriptomes. Finally, we integrated droplet-based scRNA-seq and scATAC-seq data with scStereo-seq data for embryo samples, which improved genome coverage and incorporated epigenomic information. The plethora of multi-omics data generated in this study provided many unique angles for dissecting the molecular underpinnings of various aspects of tissue development, as we have shown in this study.

The integration of multi-omics data has enriched our analysis, enabling a more nuanced portrayal of cell states. As an example, the transcriptomes of CNS mature neurons are remarkably uniform, as demonstrated by their intertwined distribution in the UMAP space of scRNA-seq data. However, the incorporation of scATAC-seq data, which shows a higher degree of heterogeneity among mature neurons, allowed us to identify detailed neuron subtypes and investigate the regulatory mechanisms driving their differentiation. Leveraging the high heterogeneity of scATAC-seq data, we were able to dissect the differentiation process of a group of neuron subtypes in detail and identified the TF *Kr* as a key regulator. The wealth of chromatin accessibility information allowed us to further uncover TFs that co-regulated gene expression with *Kr*. We also mapped cell clusters, derived from the integration of scRNA-seq and scATAC-seq data, to their spatial positions in the scStereo-seq data. This mapping enabled us to model single-cell transcriptomic and epigenomic profiles within tissue- and developmental stage-specific contexts. Therefore, this multi-omics data integration offered an unprecedented high-resolution spatiotemporal framework for analyzing cell state dynamics, such as TF regulons and signaling pathways.

There were also inconsistencies between data generated from different techniques. For example, late-stage epidermis and somatic muscle cells identified in the scATAC-seq data lacked corresponding scRNA-seq counterparts. We observed a similar lack of coherence in CNS mature neurons, which, in addition to missing scRNA-seq data, could also resulted from a temporal mismatch between chromatin accessibility and actual gene expression in these neurons. In certain instances, the chromatin regions of neuron subtype-specific genes were open, but gene expression was delayed. This discrepancy resulted in inaccurately imputed gene expression when integrating scATAC-seq and scRNA-seq data and mismatches in the co-embedded UMAP space. Similar phenomena have been observed in the mammalian nervous system, such as the process of epigenetic priming during normal or pathological development (reviewed in Refs^107,108^).

Compared to our previous study, the point cloud-based 3D modeling of developing tissues in this work provided significantly more detailed structural information that more accurately reflects organ anatomical features. Furthermore, by aligning 3D models between different time points with *Spateo*, we were able to simultaneously track cell migration paths and alterations in gene expression. This morphometric analysis provided a unique perspective, enabling the identification of potential regulators of cell migration and differentiation. It is recognized that the eventual shape and size of an organ can be influenced by physical interactions with neighboring organs and signaling molecules from distant organs during development^109–111^. In addition to the intra-organ morphometric analysis presented here, these models can also be used to investigate the impact of inter-organ physical or biochemical contact on local gene expression changes, which in turn affect the final boundaries of organs. With the establishment of a complete synapse-resolution connectome of the *Drosophila* larval brain^112^, our 3D transcriptomes have the potential to be spatially aligned with these synapse connectivity maps. By integrating spatial transcriptome and connectome data, we can simultaneously pinpoint the spatial locations of known and yet-to-be-identified neurons and deconvolute their molecular nature, leading to a deeper understanding of their physiological functions.

The extensive datasets we generated here can be leveraged in many ways. *Drosophila* larva and pupa have provided excellent models for studying the course of post-organogenesis development and metamorphosis, yet single-cell profiling of tissues at these stages remained scant. These stages of samples in our scStereo-seq data can be readily integrated with existing larva scRNA-seq datasets^6,7,113^ to complement them with spatial information or provide a spatial framework for future single-cell studies of larval or pupal tissues. The study of early stages of *Drosophila* pupal development has been challenging due to significant tissue lysis and reformation. Our pupa scStereo-seq data provided valuable insights for investigation of tissue-specific transcriptomic changes during metamorphosis.

Moreover, the unique organization of our datasets can serve as a source of inspiration for the development of multiple types of bioinformatic algorithms and methods and can serve as benchmarking resources for such algorithms. The organism-wide 3D high-resolution features of our previous Stereo-seq datasets have already facilitated the development of several approaches for various purposes, including quantitative spatiotemporal modeling of single-cell spatial transcriptomic datasets^14^, visualization and analysis of spatial omics data^114^, construction of databases and optimization of their accessibility^115^, alignment of 2D spatial transcriptomic sections for 3D modeling^116,117^, and more. The new dataset features in this study can further assist in the development of bioinformatic approaches in many other aspects, such as cell segmentation of spatial transcriptomic data, integration of multi-omics data, spatial mapping of cell types, machine learning-based cell type and age prediction, and cell lineage tracing, among others. With the rapid development of spatial transcriptomic techniques and consequently the mounting number of datasets, these methods will serve as invaluable tools to facilitate our interpretation of multi-omics datasets.

In order to make our data more accessible, we have incorporated our datasets into the Spateo Viewer platform. This platform is a versatile and scalable web application specifically designed for the exploration of spatial transcriptomics data. Accessible through our online data portal, Flysta3D, the Spateo Viewer provides user-friendly access to our 3D models. It enables interactive visualization of gene expression, activity of gene groups, and a variety of other customizable parameters within spatiotemporal contexts. We believe that our comprehensive multi-omics database will serve as a catalyst for systematic research into *Drosophila* development, facilitating a deeper understanding of organism-wide spatiotemporal dynamics.

## LIMITATIONS OF THIS STUDY

Our multi-modal analysis of the dataset revealed a wealth of information and demonstrated its potential for systematic spatiotemporal analysis of *Drosophila* development. However, there are still some areas that require improvement in future studies.

The scRNA-seq and scATAC-seq data in this study were obtained from separate samples. We aimed to align cells from the same tissues and same developmental stages between datasets for integrated multi-omics analysis through the control of collection window and inference of single cell developmental age. However, it is still possible that data used for integration were from different states of cells. Methods for simultaneous capture of transcriptomic and chromatin accessibility profiles from single cells have been developed lately^118,119^, which may provide better integration results, especially addressing the temporal mismatch issue between scRNA-seq and scATAC-seq data. With future technical improvements, spatial information of cells may also be captured simultaneously to generate actual multi-omics profiles for each single cell.

This multi-omics atlas was generated exclusively from the genetic background strain *w1118*. However, investigating developmental regulation or disease mechanisms often involves genetic perturbations, such as knockdown/knockout of key regulator genes, or changes in environmental conditions, such as pathogen infection and drug treatment. Therefore, in the future, we plan to expand our study to include *Drosophila* models with various genetic mutations or subjected to different infection and/or treatment conditions to establish organism-wide single-cell multi-omics atlases. This approach will be particularly beneficial for studying complex physiological processes that involve multiple tissues in response to genetic perturbations, such as the progression of neurodegenerative diseases, gut-brain axis communication, and multi-organ metabolic diseases. The pipeline established in this study can serve as a basis for such investigations, enabling the generation of comprehensive datasets that incorporate genetic and environmental variability.

## ACKNOWLEDGEMENTS

This work was supported by Shenzhen Key Laboratory of Gene Regulation and Systems Biology (Grant No. ZDSYS20200811144002008 to Y.H.), National Natural Science Foundation of China (Grant No. 32100684 to Q.H.), National Key R&D Program of China (Grant No. 2022YFC3400405 to X.X. and L.L.), National Key R&D Program of China (Grant No. 2022YFC3400300 to M.W.), Shenzhen Science and Technology Innovation Program (Grant No. KQTD20180411143432337, China) (to Y.H., W.C., and Q.H.), Guangdong Genomics Data Center (2021B1212100001 to T.Y. and J.C.). Part of data analysis work was performed on the STOmic Cloud (https://cloud.stomics.tech). Library sequencing was performed in China National GeneBank. This work was supported by Center for Computational Science and Engineering and Core Research Facilities at Southern University of Science and Technology. We thank Dr. Xiaojie Qiu and Dr. Martin Vingron for helpful suggestions on data analysis and Dr. Mariana Wolfner for helpful comments on the manuscript. We thank Biorender for graphical resources used in **Figure 1A** and Graphical Abstract.

## AUTHOR CONTRIBUTIONS

Q.H., L.L., X.X., and Y.H. conceived the idea; C.L., Y.G., W.C., L.L., X.X., and Y.H. supervised the work; Q.H. prepared the samples; M.W., Z.T., Y.H., N.Y., W.M., W.L., C.L., and X.L. prepared the sequencing library and performed sequencing; M.W., Q.H., Z.T., L.K., J.Y., R.X., Y.Z., Z.C., Z.J., K.O., X.W., Y.B., and M.L. performed computational analysis; Q.H., M.W., Z.T., L.K., and Y.H. interpreted the data; Q.H., Y.Z., T.Y, Y.W., and Z.Y. performed experimental validation; T.Y., and J.C. established the data portal and Flysta3D website; Q.H., M.W., W.C., L.L., X.X., and Y.H. acquired fundings; Q.H., M.W., Z.T., and Y.H. wrote the manuscript; All authors reviewed and edited the manuscript.

## DECLARATION OF INTERESTS

The chip, procedure, and application of Stereo-seq are covered in pending patents. Employees of BGI have stock holdings in BGI.

## METHODS

### RESOURCE AVAILABILITY

#### Lead contact

Further information and requests for the resources and reagents may be directed to the corresponding author Yuhui Hu.

### Materials availability

All materials used for Stereo-seq, MGI C4 scRNA-seq, and MGI C4 scATAC-seq are commercially available.

### Data and code availability

Raw data generated by Stereo-seq, scRNA-seq, and scATAC-seq in this study and associated analysis protocols and software can be accessed in our online database, Flysta3D. All data were analyzed with standard programs and packages, as detailed in Method details. Processed matrices can be accessed through Mendeley Data (https://doi.org/10.17632/tvvjfr3c6j.1, https://doi.org/10.17632/29695x8txs.1, and https://doi.org/10.17632/4zf847bxcd.1). All custom codes using open-source software to support this study are provided in a public GitHub repository. Any additional information required to re-analyze the data reported in this study is available from the lead contact upon request.

## EXPERIMENTAL MODEL AND SUBJECT DETAILS

### Fly strain maintenance

All Stereo-seq, scRNA-seq, and scATAC-seq samples were from *Drosophila* strain *w1118*. Flies were maintained on cornmeal-sucrose-agar media in a 25 ℃ incubator on a 12 h/12 h light/dark cycle.

### Fly sample preparation

Samples were prepared and embedded for cryosection and Stereo-seq as previously described^13^. Unless otherwise specified, the samples were sectioned along the left-right axis to represent sagittal planes.

For scRNA-seq, single cells were isolated and fixed following protocols described in Ref^120^ and stored at −20 ℃ until further use.

For scATAC-seq, embryos at the desired stages were collected from a population cage. The embryos were transferred to a 70 μm cell strainer, dechorionated in commercial bleach for 3 min, rinsed with ddH2O, and dried on a Kimwipe. Dechorionated embryos were then snap-frozen in liquid nitrogen and stored at −80 ℃ until further use.

## METHOD DETAILS

See method details in Supplemental Information.

## SUPPLEMENTAL INFORMATION

Supplemental Figures, Tables, Movies, and Data can be found in the Supplemental Information.

## Notes

https://doi.org/10.17632/tvvjfr3c6j.1

https://doi.org/10.17632/29695x8txs.1

https://doi.org/10.17632/4zf847bxcd.1

