## Supplemental Information for "A single-cell 3D spatiotemporal multi-omics atlas from *Drosophila* embryogenesis to metamorphosis"

### SUPPLEMENTAL FIGURE LEGENDS

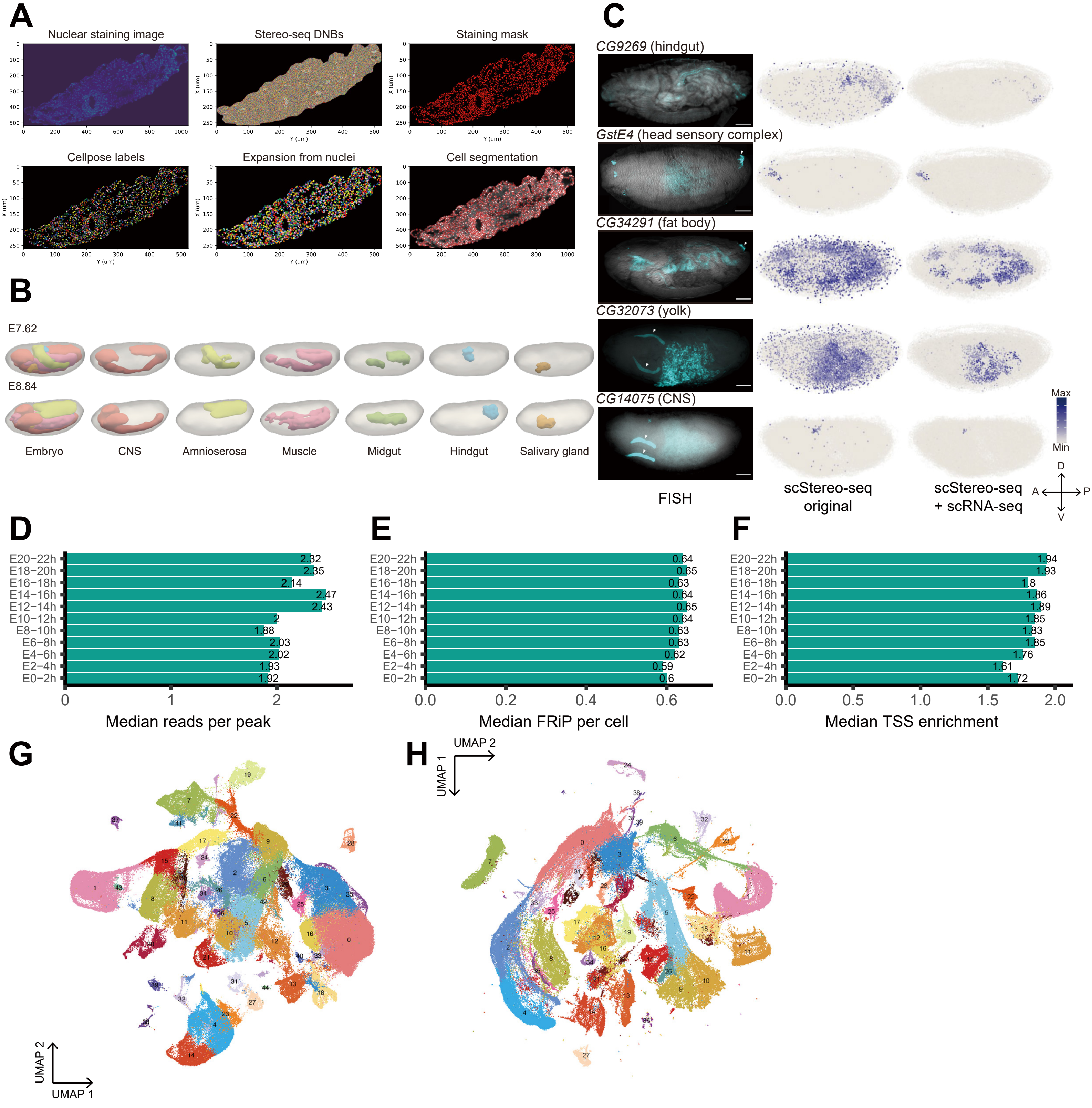
A
Nuclear staining image
Stereo-seq DNBs
Staining mask
Cellpose labels
Expansion from nuclei
Cell segmentation
X (um)
Y (um)
B
E7.62
E8.84
Embryo
CNS
Amnioserosa
Muscle
Midgut
Hindgut
Salivary gland
C
CG9269 (hindgut)
GstE4 (head sensory complex)
CG34291 (fat body)
CG32073 (yolk)
CG14075 (CNS)
FISH
scStereo-seq original
scStereo-seq + scRNA-seq
Max
Min
D
V
A
P
D
E20−22h 2.32
E18−20h 2.35
E16−18h 2.14
E14−16h 2.47
E12−14h 2.43
E10−12h 2
E8−10h 1.88
E6−8h 2.03
E4−6h 2.02
E2−4h 1.93
E0−2h 1.92
Median reads per peak
E
E20−22h 0.64
E18−20h 0.65
E16−18h 0.63
E14−16h 0.64
E12−14h 0.65
E10−12h 0.64
E8−10h 0.63
E6−8h 0.63
E4−6h 0.62
E2−4h 0.59
E0−2h 0.6
Median FRiP per cell
F
E20−22h 1.94
E18−20h 1.93
E16−18h 1.8
E14−16h 1.86
E12−14h 1.89
E10−12h 1.85
E8−10h 1.83
E6−8h 1.85
E4−6h 1.76
E2−4h 1.61
E0−2h 1.72
Median TSS enrichment
G
UMAP 2
UMAP 1
H
UMAP 2
UMAP 1

**Figure S1 Data quality control and pre-processing. Related to Figure 1.**

**(A)** Cell segmentation workflow of a representative L1 early section. **(B)** 3D modeling of individual tissues in representative embryo scStereo-seq samples, showing mesh models of tissues and the entire embryo. **(C)** Additional examples for **Figure 1C**. **(D-F)** Additional quality control statistics for scATAC-seq data, with bar plots showing **(D)** median reads per peak, **(E)** median fraction of reads in peaks (FRiP) per cell, and **(F)** median TSS enrichment scores across sample batches. **(G-H)** UMAP plots of aggregated **(G)** scRNA-seq and **(H)** scATAC-seq data after coarse unsupervised clustering, color coded and labeled by cluster number.

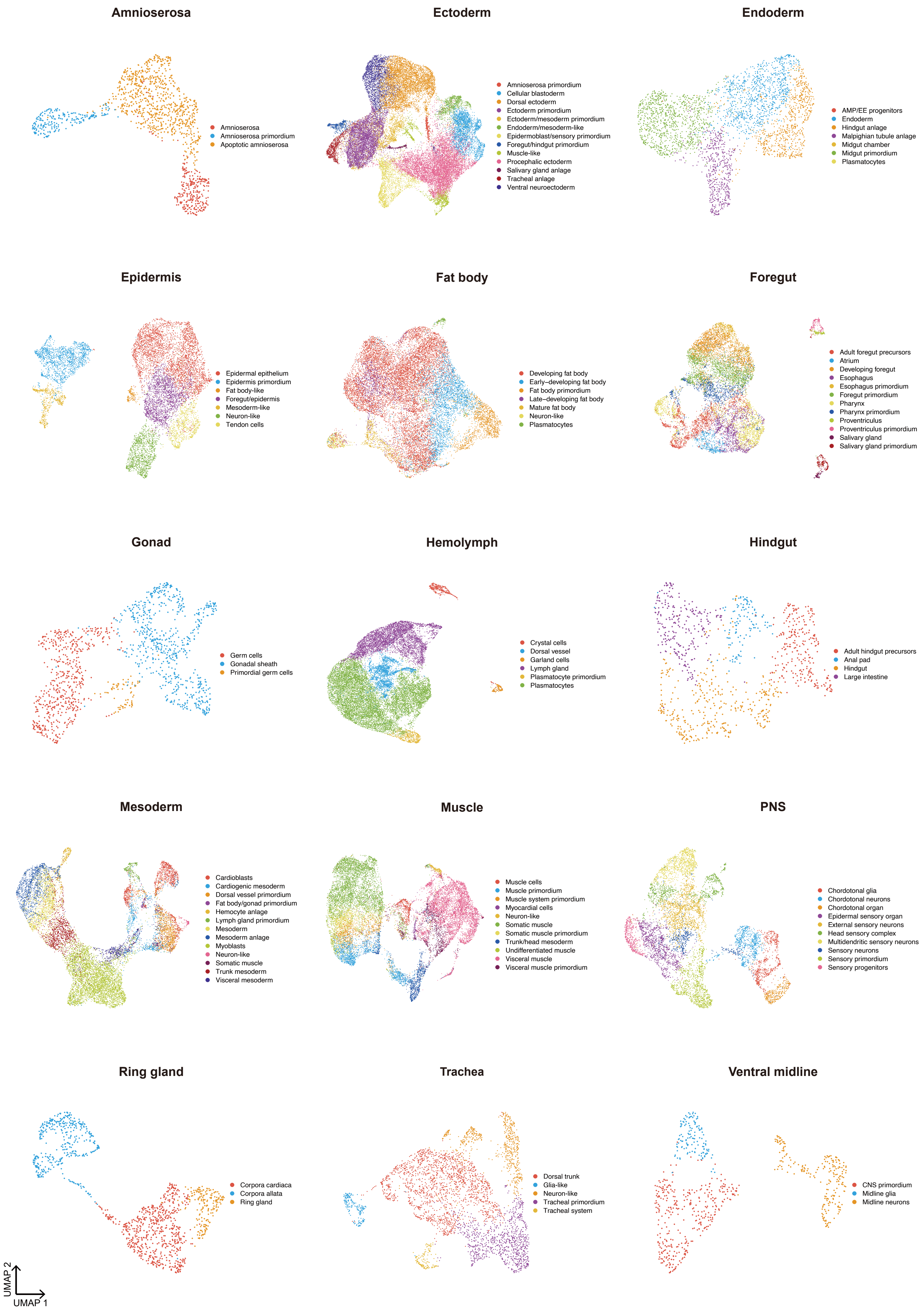

Amnioserosa
Amnioserosa
Amnioserosa primordium
Apoptotic amnioserosa
Ectoderm
Amnioserosa primordium
Cellular blastoderm
Dorsal ectoderm
Ectoderm primordium
Ectoderm/mesoderm primordium
Endoderm/mesoderm-like
Epidermoblast/sensory primordium
Foregut/hindgut primordium
Muscle-like
Procephalic ectoderm
Salivary gland anlage
Tracheal anlage
Ventral neuroectoderm
Endoderm
AMP/EE progenitors
Endoderm
Hindgut anlage
Malpighian tubule anlage
Midgut chamber
Midgut primordium
Plasmatocytes
Epidermis
Epidermal epithelium
Epidermis primordium
Fat body-like
Foregut/epidermis
Mesoderm-like
Neuron-like
Tendon cells
Fat body
Developing fat body
Early–developing fat body
Fat body primordium
Late–developing fat body
Mature fat body
Neuron-like
Plasmatocytes
Foregut
Adult foregut precursors
Atrium
Developing foregut
Esophagus
Esophagus primordium
Foregut primordium
Pharynx
Pharynx primordium
Proventriculus
Proventriculus primordium
Salivary gland
Salivary gland primordium
Gonad
Germ cells
Gonadal sheath
Primordial germ cells
Hemolymph
Crystal cells
Dorsal vessel
Garland cells
Lymph gland
Plasmatocyte primordium
Plasmatocytes
Hindgut
Adult hindgut precursors
Anal pad
Hindgut
Large intestine
Mesoderm
Cardioblasts
Cardiogenic mesoderm
Dorsal vessel primordium
Fat body/gonad primordium
Hemocyte anlage
Lymph gland primordium
Mesoderm
Mesoderm anlage
Myoblasts
Neuron-like
Somatic muscle
Trunk mesoderm
Visceral mesoderm
Muscle
Muscle cells
Muscle primordium
Muscle system primordium
Myocardial cells
Neuron-like
Somatic muscle
Somatic muscle primordium
Trunk/head mesoderm
Undifferentiated muscle
Visceral muscle
Visceral muscle primordium
PNS
Chordotonal glia
Chordotonal neurons
Chordotonal organ
Epidermal sensory organ
External sensory neurons
Head sensory complex
Multidendritic sensory neurons
Sensory neurons
Sensory primordium
Sensory progenitors
Ring gland
Corpora cardiaca
Corpora allata
Ring gland
Trachea
Dorsal trunk
Glia-like
Neuron-like
Tracheal primordium
Tracheal system
Ventral midline
CNS primordium
Midline glia
Midline neurons
UMAP 2
UMAP 1

**Figure S2 Subclustering and annotation of tissue clusters in scRNA-seq data. Related to Figure 2.**

See subcluster marker genes and notable markers of each subcluster in **Table S4**. Subclustering results of CNS and midgut are presented and discussed in **Figure 5** and **Figure 6**, respectively.

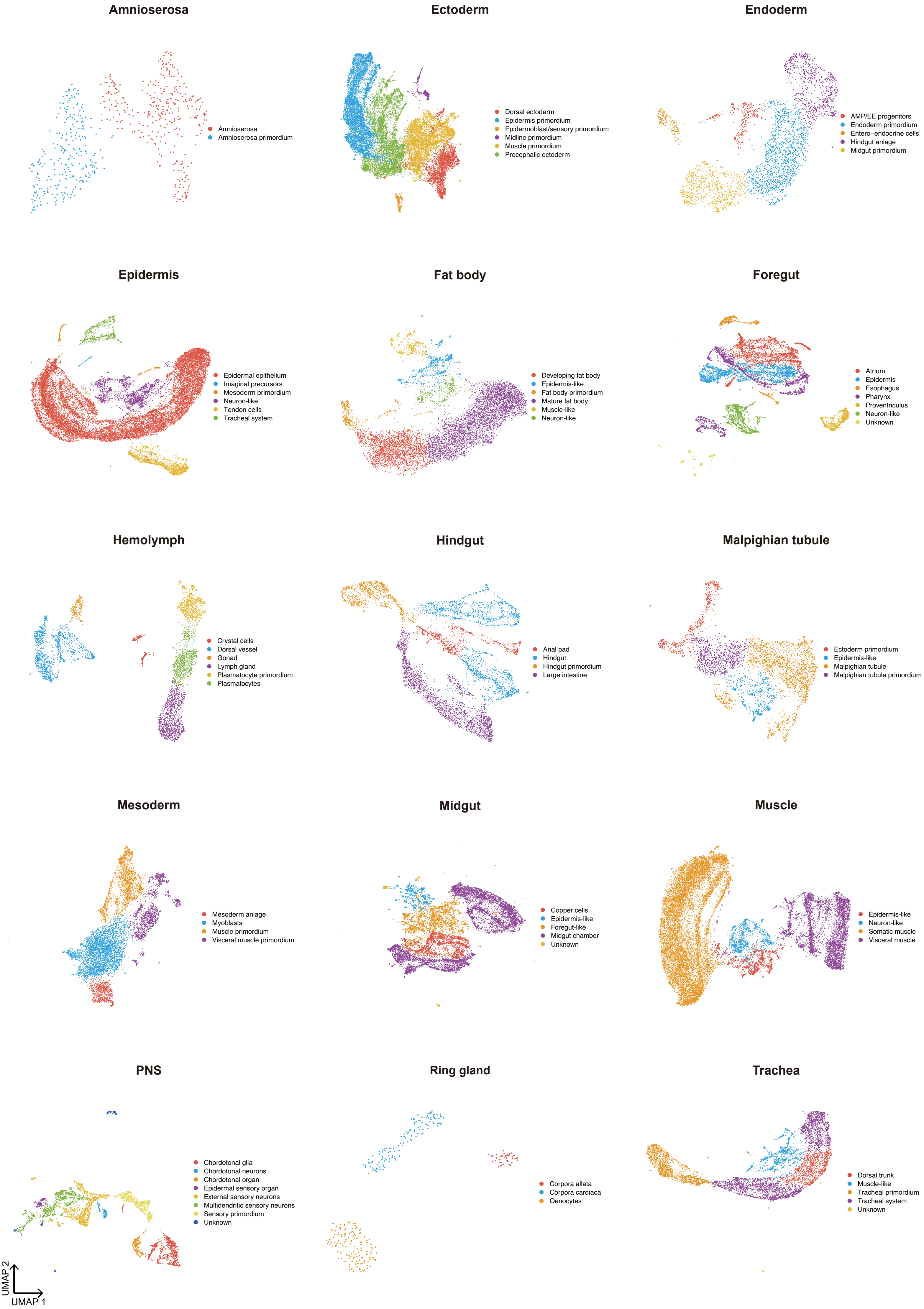

Amnioserosa
Amnioserosa
Amnioserosa primordium
Ectoderm
Dorsal ectoderm
Epidermis primordium
Epidermoblast/sensory primordium
Midline primordium
Muscle primordium
Procephalic ectoderm
Endoderm
AMP/EE progenitors
Endoderm primordium
Entero−endocrine cells
Hindgut anlage
Midgut primordium
Epidermis
Epidermal epithelium
Imaginal precursors
Mesoderm primordium
Neuron-like
Tendon cells
Tracheal system
Fat body
Developing fat body
Epidermis-like
Fat body primordium
Mature fat body
Muscle-like
Neuron-like
Foregut
Atrium
Epidermis
Esophagus
Pharynx
Proventriculus
Neuron-like
Unknown
Hemolymph
Crystal cells
Dorsal vessel
Gonad
Lymph gland
Plasmatocyte primordium
Plasmatocytes
Hindgut
Anal pad
Hindgut
Hindgut primordium
Large intestine
Malpighian tubule
Ectoderm primordium
Epidermis-like
Malpighian tubule
Malpighian tubule primordium
Mesoderm
Mesoderm anlage
Myoblasts
Muscle primordium
Visceral muscle primordium
Midgut
Copper cells
Epidermis-like
Foregut-like
Midgut chamber
Unknown
Muscle
Epidermis-like
Neuron-like
Somatic muscle
Visceral muscle
PNS
Chordotonal glia
Chordotonal neurons
Chordotonal organ
Epidermal sensory organ
External sensory neurons
Multidendritic sensory neurons
Sensory primordium
Unknown
Ring gland
Corpora allata
Corpora cardiaca
Oenocytes
Trachea
Dorsal trunk
Muscle-like
Tracheal primordium
Tracheal system
Unknown
UMAP 2
UMAP 1

**Figure S3 Subclustering and annotation of tissue clusters in scATAC-seq data. Related to Figure 2.**

See subcluster marker genes and notable markers of each subcluster in **Table S4**. Subclustering results of CNS and midgut are presented and discussed in **Figure 5** and **Figure 6**, respectively.

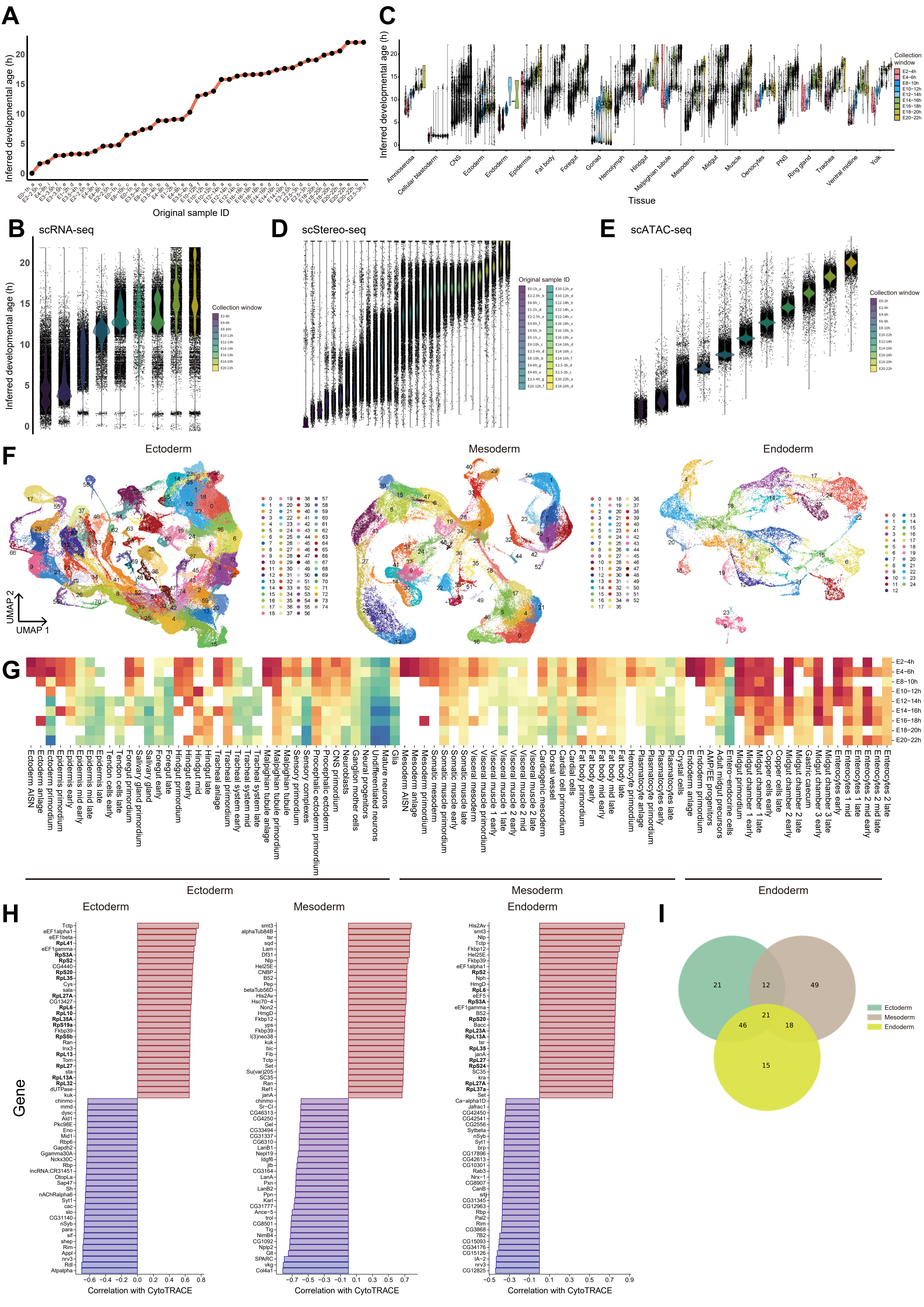
A
Inferred developmental age (h)
Original sample ID
C
Inferred developmental age (h)
Tissue
Amnioserosa
Cellular blastoderm
CNS
Ectoderm
Endoderm
Epidermis
Fat body
Foregut
Gonad
Hemolymph
Hindgut
Malpighian tubule
Mesoderm
Midgut
Muscle
Oenocytes
PNS
Ring gland
Trachea
Ventral midline
Yolk
Collection window
E2−4h
E4−6h
E8−10h
E10−12h
E12−14h
E14−16h
E16−18h
E18−20h
E20−22h
B
scRNA-seq
Collection window
D
scStereo-seq
Original sample ID
E
scATAC-seq
Collection window
F
Ectoderm
Mesoderm
Endoderm
UMAP 2
UMAP 1
G
E2−4h
E4−6h
E8−10h
E10−12h
E12−14h
E14−16h
E16−18h
E18−20h
E20−22h
Ectoderm AISN
Ectoderm anlage
Ectoderm primordium
Epidermis primordium
Epidermis early
Epidermis mid early
Epidermis mid late
Epidermis late
Tendon cells early
Tendon cells late
Foregut primordium
Salivary gland
Salivary gland primordium
Foregut early
Foregut late
Hindgut primordium
Hindgut early
Hindgut mid
Hindgut late
Tracheal anlage
Tracheal primordium
Tracheal system early
Tracheal system mid
Tracheal system late
Malpighian tubule primordium
Malpighian tubule anlage
Malpighian tubule
Sensory primordium
Sensory complexes
Procephalic ectoderm
Procephalic ectoderm primordium
CNS primordium
Neuroblasts
Ganglion mother cells
Neural progenitors
Undifferentiated neurons
Mature neurons
Glia
Mesoderm AISN
Mesoderm anlage
Mesoderm primordium
Somatic mesoderm
Somatic muscle primordium
Somatic muscle early
Somatic muscle late
Visceral mesoderm
Visceral muscle primordium
Visceral muscle 1 early
Visceral muscle 1 late
Visceral muscle 2 early
Visceral muscle 2 mid
Visceral muscle 2 late
Cardiogenic mesoderm
Dorsal vessel
Cardial cell primordium
Cardial cells
Fat body primordium
Fat body early
Fat body mid early
Fat body mid late
Fat body late
Hemocyte primordium
Plasmatocyte anlage
Plasmatocyte primordium
Plasmatocytes early
Plasmatocytes late
Crystal cells
Endoderm anlage
Endoderm primordium
AMP/EE progenitors
Adult midgut precursors
Entero-endocrine cells
Midgut primordium
Midgut chamber 1 early
Midgut chamber 1 late
Copper cells early
Copper cells late
Midgut chamber 2 early
Midgut chamber 2 late
Gastric caecum
Midgut chamber 3 early
Midgut chamber 3 late
Enterocytes early
Enterocytes 1 mid
Enterocytes 1 late
Enterocytes 2 mid early
Enterocytes 2 mid late
Enterocytes 2 late
Ectoderm
Mesoderm
Endoderm
H
Ectoderm
Mesoderm
Endoderm
Gene
Correlation with CytoTRACE
I
Ectoderm
Mesoderm
Endoderm
21
12
49
21
46
18
15

**Figure S4 Developmental age-matched integration of multi-omics data and dynamics of tissue development trajectories. Related to Figure 2 and Figure 3.**

**(A)** Line plot showing *RAPToR* inferred developmental age of embryo scStereo-seq samples. **(B)** Violin plot showing *RAPToR* inferred developmental age of scRNA-seq samples. **(C)** Same as **(B)** but showing tissues for each stage. **(D)** Same as **(B)** but for cell bins in individual scStereo-seq samples. **(E)** Violin plot showing neural network model inferred developmental age of scATAC-seq samples. **(F)** Co-embedded UMAP plots of integrated scRNA-seq scATAC-seq data of three germ layers after unsupervised clustering, color coded and labeled by cluster number. **(G)** Heatmap showing median *CytoTRACE* scores of tissue cell types based on scRNA-seq data along tissue development trajectories. Blank cells indicate absence of tissue types at corresponding stages. **(H)** Bar plots showing top 30 genes that are most positively/negatively correlated with *CytoTRACE* scores in three germ layers. Ribosome protein genes are in bold. **(I)** Venn diagram of top 100 genes most positively correlated with *CytoTRACE* scores in three germ layers.

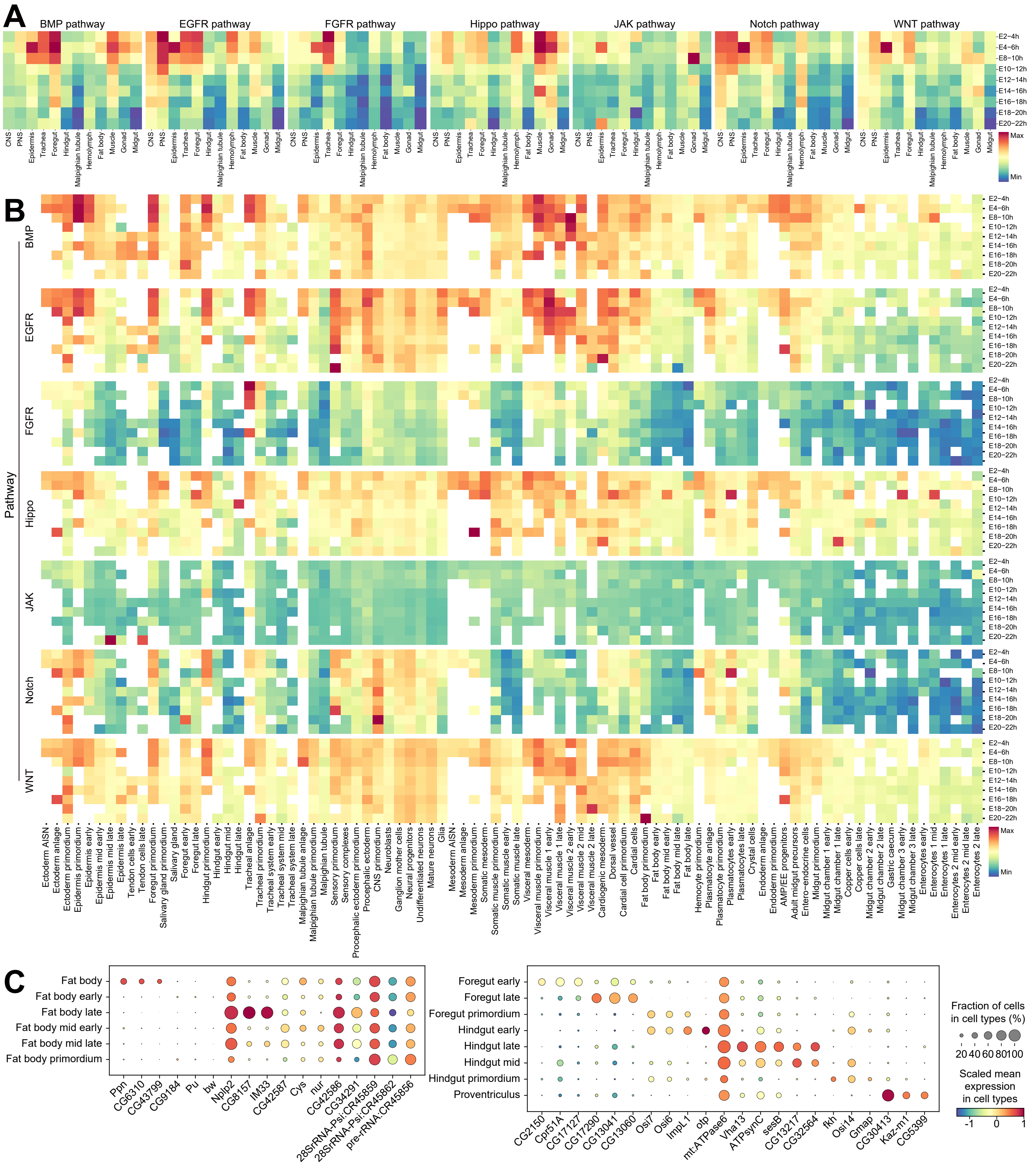
A
BMP pathway
EGFR pathway
FGFR pathway
Hippo pathway
JAK pathway
Notch pathway
WNT pathway
CNS
PNS
Epidermis
Trachea
Foregut
Hindgut
Malpighian tubule
Hemolymph
Fat body
Muscle
Gonad
Midgut
E2–4h
E4–6h
E8–10h
E10–12h
E12–14h
E14–16h
E16–18h
E18–20h
E20–22h
Max
Min
B
Pathway
BMP
EGFR
FGFR
Hippo
JAK
Notch
WNT
Ectoderm AISN
Ectoderm anlage
Ectoderm primordium
Epidermis primordium
Epidermis early
Epidermis mid early
Epidermis mid late
Epidermis late
Tendon cells early
Tendon cells late
Foregut primordium
Salivary gland primordium
Salivary gland
Foregut early
Foregut late
Hindgut primordium
Hindgut early
Hindgut mid
Hindgut late
Tracheal anlage
Tracheal primordium
Tracheal system early
Tracheal system mid
Tracheal system late
Malpighian tubule anlage
Malpighian tubule primordium
Malpighian tubule
Sensory primordium
Sensory complexes
Procephalic ectoderm primordium
Procephalic ectoderm
CNS primordium
Neuroblasts
Ganglion mother cells
Neural progenitors
Undifferentiated neurons
Mature neurons
Glia
Mesoderm AISN
Mesoderm anlage
Mesoderm primordium
Somatic mesoderm
Somatic muscle primordium
Somatic muscle early
Somatic muscle late
Visceral mesoderm
Visceral muscle primordium
Visceral muscle 1 early
Visceral muscle 1 late
Visceral muscle 2 early
Visceral muscle 2 mid
Visceral muscle 2 late
Cardiogenic mesoderm
Dorsal vessel
Cardial cell primordium
Cardial cells
Fat body primordium
Fat body early
Fat body mid early
Fat body mid late
Fat body late
Hemocyte primordium
Plasmatocyte anlage
Plasmatocyte primordium
Plasmatocytes early
Plasmatocytes late
Crystal cells
Endoderm anlage
Endoderm primordium
AMP/EE progenitors
Adult midgut precursors
Entero-endocrine cells
Midgut primordium
Midgut chamber 1 early
Midgut chamber 1 late
Copper cells early
Copper cells late
Midgut chamber 2 early
Midgut chamber 2 late
Gastric caecum
Midgut chamber 3 early
Midgut chamber 3 late
Enterocytes early
Enterocytes 1 mid
Enterocytes 1 late
Enterocytes 2 mid early
Enterocytes 2 mid late
Enterocytes 2 late
C
Fat body
Fat body early
Fat body late
Fat body mid early
Fat body mid late
Fat body primordium
Ppn
CG6310
CG43799
CG9184
Pu
bw
Nplp2
CG8157
IM33
CG42587
Cys
nur
CG42586
CG34291
28SrRNA-Psi:CR45859
28SrRNA-Psi:CR45862
pre-rRNA:CR45856
Foregut early
Foregut late
Foregut primordium
Hindgut early
Hindgut late
Hindgut mid
Hindgut primordium
Proventriculus
CG2150
Cpr51A
CG17127
CG17290
CG13041
CG13060
Osi7
Osi6
ImpL1
otp
mt:ATPase6
Vha13
ATPsynC
sesB
CG13217
CG32564
fkh
Osi14
Gmap
CG30413
Kaz-m1
CG5399
Fraction of cells in cell types (%)
20 40 60 80 100
Scaled mean expression in cell types
-1
0
1

**Figure S5 Spatiotemporal cell type dynamics along tissue development trajectories. Related to Figure 3.**

**(A)** Heatmap showing median gene activity scores of core components of signaling pathways based on scRNA-seq data along tissue development trajectories. **(B)** Same as **(A)**, but for tissue cell types along developmental trajectories. Blank cells indicate absence of tissue types at corresponding stages. **(C)** Bubble plots showing expression levels and enrichment of cell type top marker genes in label transferred fat body and foregut/hindgut cells in scStereo-seq data.

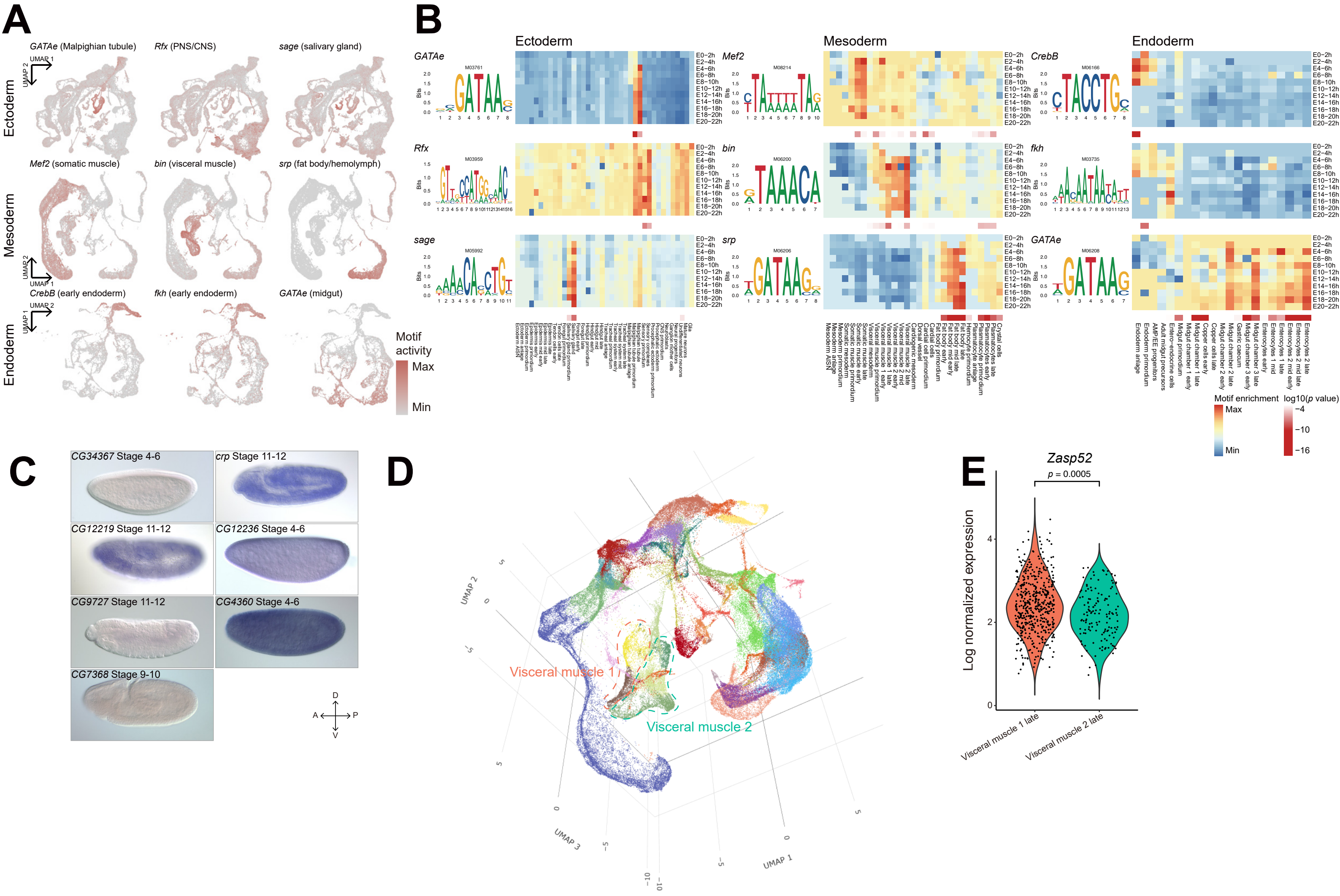

A
GATAe (Malpighian tubule)
Rfx (PNS/CNS)
sage (salivary gland)
Mef2 (somatic muscle)
bin (visceral muscle)
srp (fat body/hemolymph)
CrebB (early endoderm)
fkh (early endoderm)
GATAe (midgut)
Ectoderm
Mesoderm
Endoderm
Motif activity
Max
Min
B
Ectoderm
Mesoderm
Endoderm
Motif enrichment
log10(p value)
C
CG34367 Stage 4-6
crp Stage 11-12
CG12219 Stage 11-12
CG12236 Stage 4-6
CG9727 Stage 11-12
CG4360 Stage 4-6
CG7368 Stage 9-10
D
Visceral muscle 1
Visceral muscle 2
UMAP 1
UMAP 2
UMAP 3
E
Zasp52
p = 0.0005
Log normalized expression
Visceral muscle 1 late
Visceral muscle 2 late

**Figure S6 Transcription factor regulatory networks along tissue development trajectories. Related to Figure 4.**

**(A-B)** Same as **Figure 4A-B**, but for previously reported tissue-specific TFs. **(C)** BDGP *in situ* patterns of less characterized TFs in **Figure 4A-B**. A-P: anterior-posterior; D-V: dorsal-ventral. **(D)** 3D UMAP plot of integrated scRNA-seq and scATAC-seq data for mesoderm (also see **Data S2**). Dashed lines mark two visceral muscle groups. **(E)** Violin plot showing log normalized expression levels of *Zasp52* in scRNA-seq cells from visceral muscle 1 late and visceral muscle 2 late clusters. Wilcoxon test was used for statistical analysis.

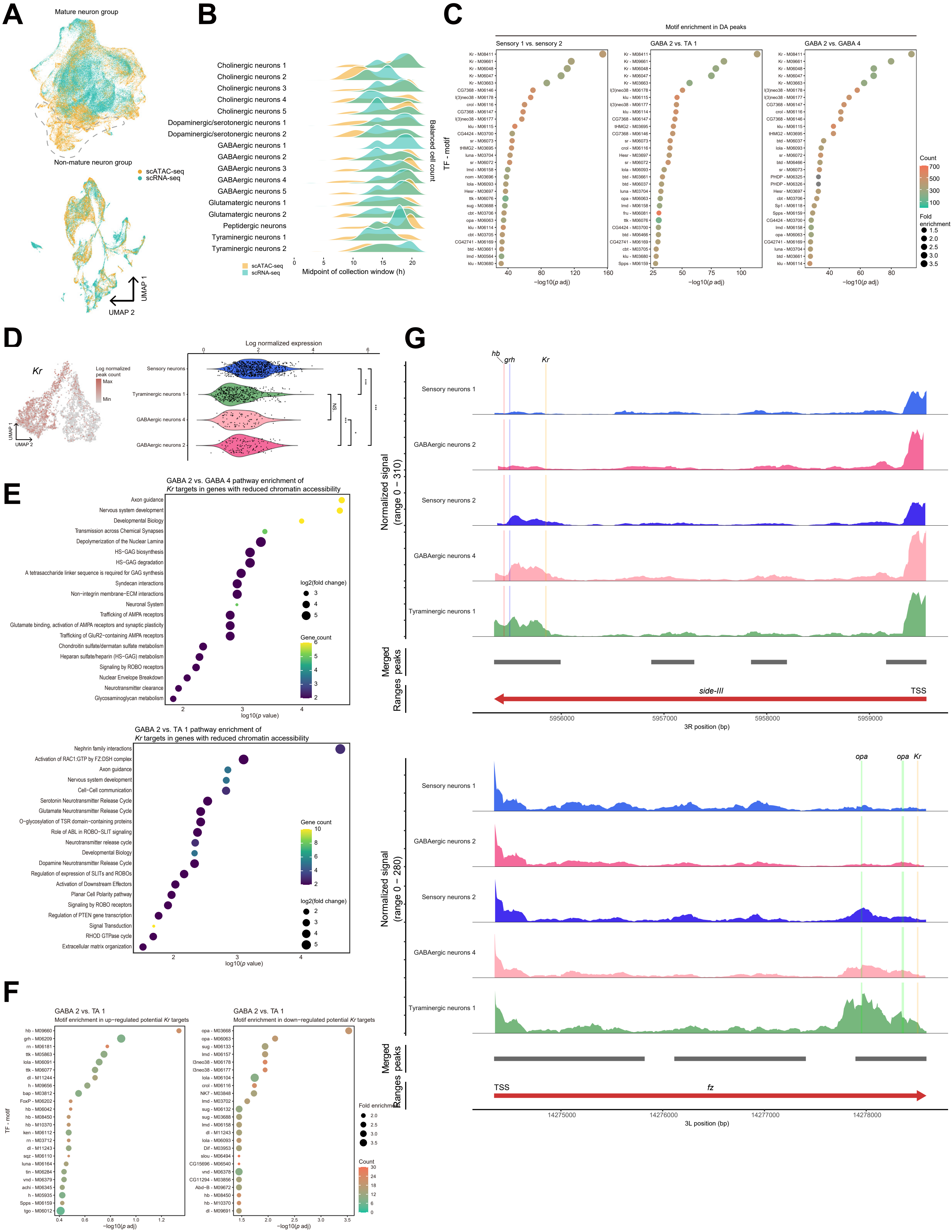
A
Mature neuron group
Non-mature neuron group
scATAC-seq
scRNA-seq
UMAP 1
UMAP 2
B
Cholinergic neurons 1
Cholinergic neurons 2
Cholinergic neurons 3
Cholinergic neurons 4
Cholinergic neurons 5
Dopaminergic/serotonergic neurons 1
Dopaminergic/serotonergic neurons 2
GABAergic neurons 1
GABAergic neurons 2
GABAergic neurons 3
GABAergic neurons 4
GABAergic neurons 5
Glutamatergic neurons 1
Glutamatergic neurons 2
Peptidergic neurons
Tyraminergic neurons 1
Tyraminergic neurons 2
Balanced cell count
Midpoint of collection window (h)
C
Motif enrichment in DA peaks
Sensory 1 vs. sensory 2
GABA 2 vs. TA 1
GABA 2 vs. GABA 4
TF - motif
−log10(p adj)
Count
Fold enrichment
D
Kr
Log normalized peak count
Max
Min
Log normalized expression
Sensory neurons
Tyraminergic neurons 1
GABAergic neurons 4
GABAergic neurons 2
NS
E
GABA 2 vs. GABA 4 pathway enrichment of Kr targets in genes with reduced chromatin accessibility
Axon guidance
Nervous system development
Developmental Biology
Transmission across Chemical Synapses
Depolymerization of the Nuclear Lamina
HS−GAG biosynthesis
HS−GAG degradation
A tetrasaccharide linker sequence is required for GAG synthesis
Syndecan interactions
Non−integrin membrane−ECM interactions
Neuronal System
Trafficking of AMPA receptors
Glutamate binding, activation of AMPA receptors and synaptic plasticity
Trafficking of GluR2−containing AMPA receptors
Chondroitin sulfate/dermatan sulfate metabolism
Heparan sulfate/heparin (HS−GAG) metabolism
Signaling by ROBO receptors
Nuclear Envelope Breakdown
Neurotransmitter clearance
Glycosaminoglycan metabolism
log2(fold change)
Gene count
log10(p value)
GABA 2 vs. TA 1 pathway enrichment of Kr targets in genes with reduced chromatin accessibility
Nephrin family interactions
Activation of RAC1:GTP by FZ:DSH complex
Axon guidance
Nervous system development
Cell−Cell communication
Serotonin Neurotransmitter Release Cycle
Glutamate Neurotransmitter Release Cycle
O−glycosylation of TSR domain−containing proteins
Role of ABL in ROBO−SLIT signaling
Neurotransmitter release cycle
Developmental Biology
Dopamine Neurotransmitter Release Cycle
Regulation of expression of SLITs and ROBOs
Activation of Downstream Effectors
Planar Cell Polarity pathway
Signaling by ROBO receptors
Regulation of PTEN gene transcription
Signal Transduction
RHOD GTPase cycle
Extracellular matrix organization
Gene count
log2(fold change)
log10(p value)
F
GABA 2 vs. TA 1
Motif enrichment in up-regulated potential Kr targets
GABA 2 vs. TA 1
Motif enrichment in down-regulated potential Kr targets
TF - motif
−log10(p adj)
Fold enrichment
Count
G
hb
grh
Kr
Sensory neurons 1
GABAergic neurons 2
Sensory neurons 2
GABAergic neurons 4
Tyraminergic neurons 1
Normalized signal (range 0 − 310)
Merged peaks
Ranges
side-III
TSS
3R position (bp)
opa
opa
Kr
Normalized signal (range 0 − 280)
TSS
fz
3L position (bp)

**Figure S7 Differentiation trajectories of mature neurons revealed by scATAC-seq. Related to Figure 5.**

**(A)** Co-embedding of neuron group and non-neuron group of CNS cells from scRNA-seq and scATAC-seq data in the same UMAP plot. Dashed lines mark cell clusters in scATAC-seq data that miss corresponding cells in scRNA-seq data. **(B)** Ridge plot showing count distribution of mature neuron cell types across collection windows of scRNA-seq and scATAC-seq data. Cells are down sampled and balanced across collection windows and cell types. **(C)** Bubble plots showing motif enrichment in top 1000 DA peaks among neuron subtypes. **(D)** Left: UMAP plot showing normalized peak count of *Kr* in scATAC-seq data; right: violin plot showing log normalized expression levels of *Kr* in scRNA-seq data. Wilcoxon test was used for statistical analysis. NS., not significant; *, $p < 0.05$; ***, $p < 0.001$. **(E)** Bubble plots showing REACTOME pathway enrichment of potential *Kr* target genes with reduced chromatin accessibility comparing GABAergic neurons 2 with GABAergic neurons 4, and GABAergic neurons 2 with tyraminergic neurons 1. **(F)** Bubble plots showing motif enrichment among *Kr* target genes with reduced chromatin accessibility and up- or down-regulated expression level, excluding *Kr* motifs. **(G)** Chromosomal regions around transcription start sites (TSS) of *side-III* and *fz*, showing detected peaks in neuron subtypes from scATAC-seq data and binding motifs of *Kr*, *hb*, *grh*, and *opa*.

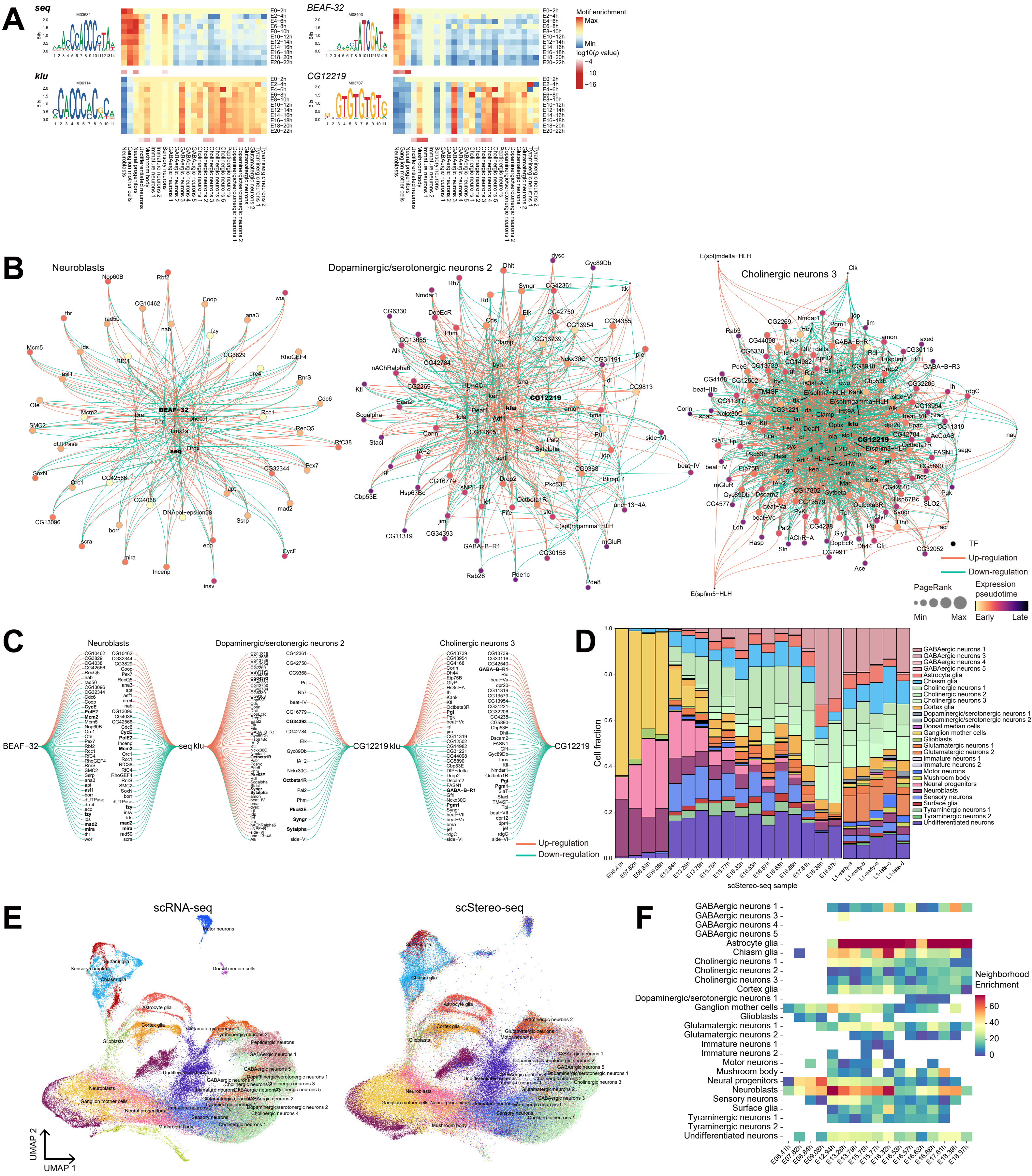

**Figure S8 Multi-omics dissection of gene regulation during embryo CNS development. Related to Figure 5.**

**(A)** TF motif enrichment during neuron development, showing TF genes, their corresponding motifs (left), motif enrichment heatmap (upper right) and enrichment *p* value heatmap (lower right) across cell types and developmental stages in CNS neuron cell types in scATAC-seq data. TFs with reported functions are in bold. **(B)** *Pando* identified GRNs of TFs (highlighted in bold) in representative cell types in **(A)**. **(C)** *Pando* identified regulons of TFs *BEAF-32* and *seq* in neuroblasts and those of TFs *klu* and *CG12219* in dopaminergic/serotonergic neurons 2 and cholinergic neurons 3. Genes in bold are discussed in detail in the main text. **(D)** Bar plot showing cell type composition of CNS in scStereo-seq samples. Cell types are label transferred from integrated scRNA-seq and scATAC-seq data. **(E)** Co-embedding of CNS cells from scRNA-seq and embryo scStereo-seq data in the same UMAP plots, labeled with original scRNA-seq annotations or transferred annotations. **(F)** Heatmap showing neighborhood enrichment scores of cell types from **(E)** across scStereo-seq samples. Blank cells indicate absence of label transferred cell types or lack of enrichment in corresponding samples.

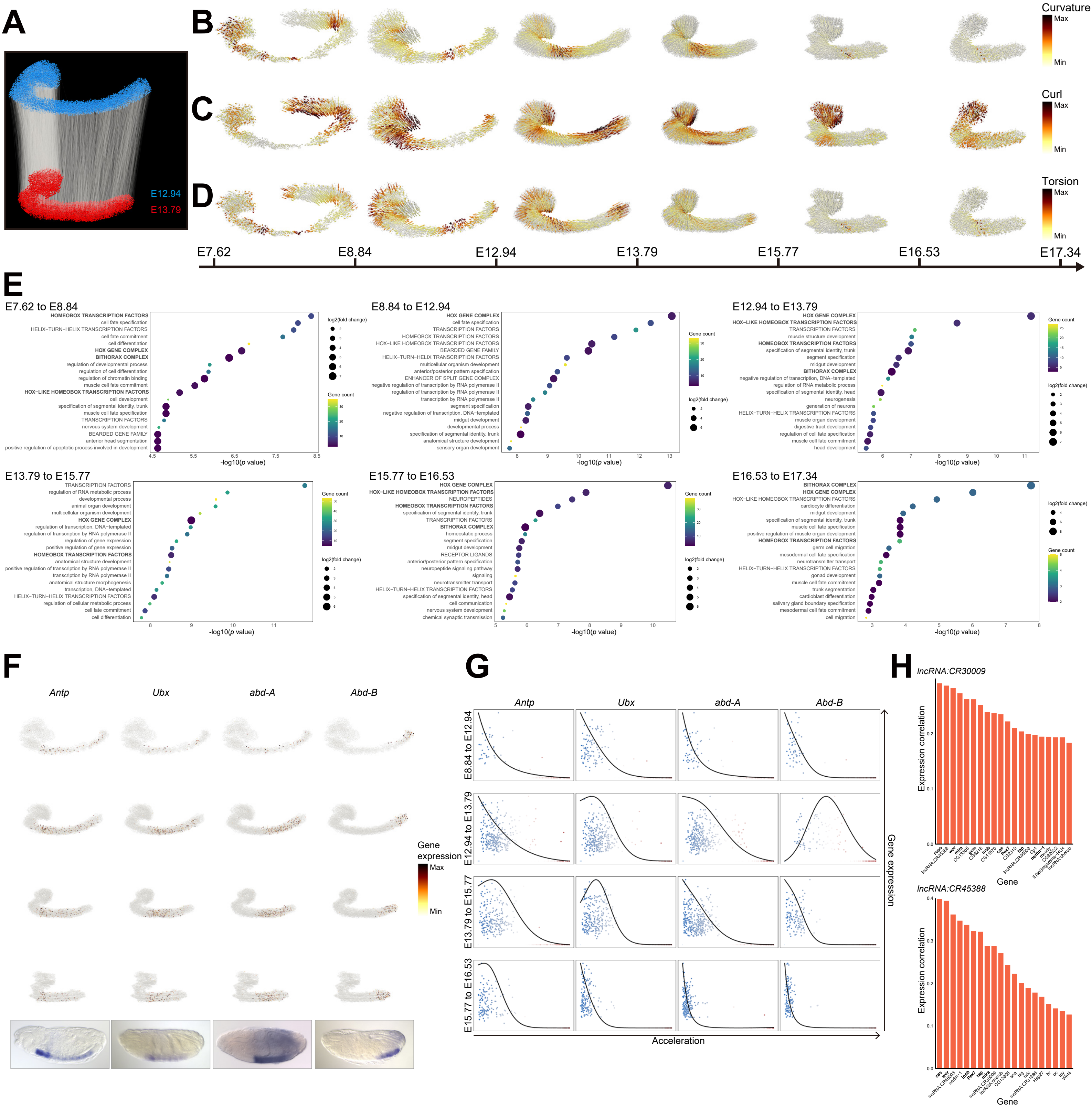
A
E12.94
E13.79
B
C
D
Curvature
Max
Min
Curl
Max
Min
Torsion
Max
Min
E7.62
E8.84
E12.94
E13.79
E15.77
E16.53
E17.34
E
E7.62 to E8.84
HOMEOBOX TRANSCRIPTION FACTORS
cell fate specification
HELIX-TURN-HELIX TRANSCRIPTION FACTORS
cell fate commitment
cell differentiation
HOX GENE COMPLEX
BITHORAX COMPLEX
regulation of developmental process
regulation of cell differentiation
regulation of chromatin binding
muscle cell fate commitment
HOX-LIKE HOMEOBOX TRANSCRIPTION FACTORS
cell development
specification of segmental identity, trunk
muscle cell fate specification
TRANSCRIPTION FACTORS
nervous system development
BEARDED GENE FAMILY
anterior head segmentation
positive regulation of apoptotic process involved in development
-log10(p value)
log2(fold change)
Gene count
E8.84 to E12.94
HOX GENE COMPLEX
cell fate specification
TRANSCRIPTION FACTORS
HOMEOBOX TRANSCRIPTION FACTORS
HOX-LIKE HOMEOBOX TRANSCRIPTION FACTORS
BEARDED GENE FAMILY
HELIX-TURN-HELIX TRANSCRIPTION FACTORS
multicellular organism development
anterior/posterior pattern specification
ENHANCER OF SPLIT GENE COMPLEX
negative regulation of transcription by RNA polymerase II
regulation of transcription by RNA polymerase II
transcription by RNA polymerase II
segment specification
negative regulation of transcription, DNA-templated
midgut development
developmental process
specification of segmental identity, trunk
anatomical structure development
sensory organ development
-log10(p value)
Gene count
log2(fold change)
E12.94 to E13.79
HOX GENE COMPLEX
HOX-LIKE HOMEOBOX TRANSCRIPTION FACTORS
TRANSCRIPTION FACTORS
muscle structure development
HOMEOBOX TRANSCRIPTION FACTORS
specification of segmental identity, trunk
segment specification
midgut development
BITHORAX COMPLEX
negative regulation of transcription, DNA-templated
regulation of RNA metabolic process
specification of segmental identity, head
neurogenesis
generation of neurons
HELIX-TURN-HELIX TRANSCRIPTION FACTORS
muscle organ development
digestive tract development
regulation of cell fate specification
muscle cell fate commitment
head development
-log10(p value)
Gene count
log2(fold change)
E13.79 to E15.77
TRANSCRIPTION FACTORS
regulation of RNA metabolic process
developmental process
animal organ development
multicellular organism development
HOX GENE COMPLEX
regulation of transcription, DNA-templated
regulation of transcription by RNA polymerase II
regulation of gene expression
positive regulation of gene expression
HOMEOBOX TRANSCRIPTION FACTORS
anatomical structure development
positive regulation of transcription by RNA polymerase II
transcription by RNA polymerase II
anatomical structure morphogenesis
transcription, DNA-templated
HELIX-TURN-HELIX TRANSCRIPTION FACTORS
regulation of cellular metabolic process
cell fate commitment
cell differentiation
-log10(p value)
Gene count
log2(fold change)
E15.77 to E16.53
HOX GENE COMPLEX
HOX-LIKE HOMEOBOX TRANSCRIPTION FACTORS
NEUROPEPTIDES
HOMEOBOX TRANSCRIPTION FACTORS
specification of segmental identity, trunk
TRANSCRIPTION FACTORS
BITHORAX COMPLEX
homeostatic process
segment specification
midgut development
RECEPTOR LIGANDS
anterior/posterior pattern specification
neuropeptide signaling pathway
signaling
neurotransmitter transport
HELIX-TURN-HELIX TRANSCRIPTION FACTORS
specification of segmental identity, head
cell communication
nervous system development
chemical synaptic transmission
-log10(p value)
Gene count
log2(fold change)
E16.53 to E17.34
BITHORAX COMPLEX
HOX GENE COMPLEX
HOX-LIKE HOMEOBOX TRANSCRIPTION FACTORS
cardiocyte differentiation
midgut development
specification of segmental identity, trunk
muscle cell fate specification
positive regulation of muscle organ development
HOMEOBOX TRANSCRIPTION FACTORS
germ cell migration
mesodermal cell fate specification
neurotransmitter transport
HELIX-TURN-HELIX TRANSCRIPTION FACTORS
gonad development
muscle cell fate commitment
trunk segmentation
cardioblast differentiation
salivary gland boundary specification
mesodermal cell fate commitment
cell migration
-log10(p value)
log2(fold change)
Gene count
F
Antp
Ubx
abd-A
Abd-B
Gene expression
Max
Min
G
Antp
Ubx
abd-A
Abd-B
E8.84 to E12.94
E12.94 to E13.79
E13.79 to E15.77
E15.77 to E16.53
Gene expression
Acceleration
H
lncRNA:CR30009
Expression correlation
Gene
lncRNA:CR45388
Expression correlation
Gene

**Figure S9 Gene expression dynamics during CNS morphometric changes. Related to Figure 5.**

**(A)** Alignment and connection of CNS cell bins between 2 representative scStereo-seq samples (E12.94 and E13.79). **(B-D)** Visualization of **(B)** curvature, **(C)** curl, and **D)** torsion scores of morphometric changes in CNS across the 7 scStereo-seq samples in **Figure 5K**. **(E)** Bubble plots showing GO (lower case terms) and FlyBase Gene Group (upper case terms) enrichment of genes associated with changes in four CNS morphometric scores. Gene terms in bold are discussed in the main text. **(F)** Spatial expression patterns of Hox family genes *Antp*, *Ubx*, *and-A*, and *Abd-B* in CNS 3D models and their BDGP *in situ* patterns from corresponding stages. **(G)** General linear model-based correlation between acceleration scores and expression levels of Hox family genes in **(F)** across scStereo-seq samples. **(H)** Bar plots showing genes that display highest expression correlation with *lncRNA:CR30009* and *lncRNA:CR45388* in scRNA-seq data. Gene names in bold indicate neuroblast or glioblast markers.

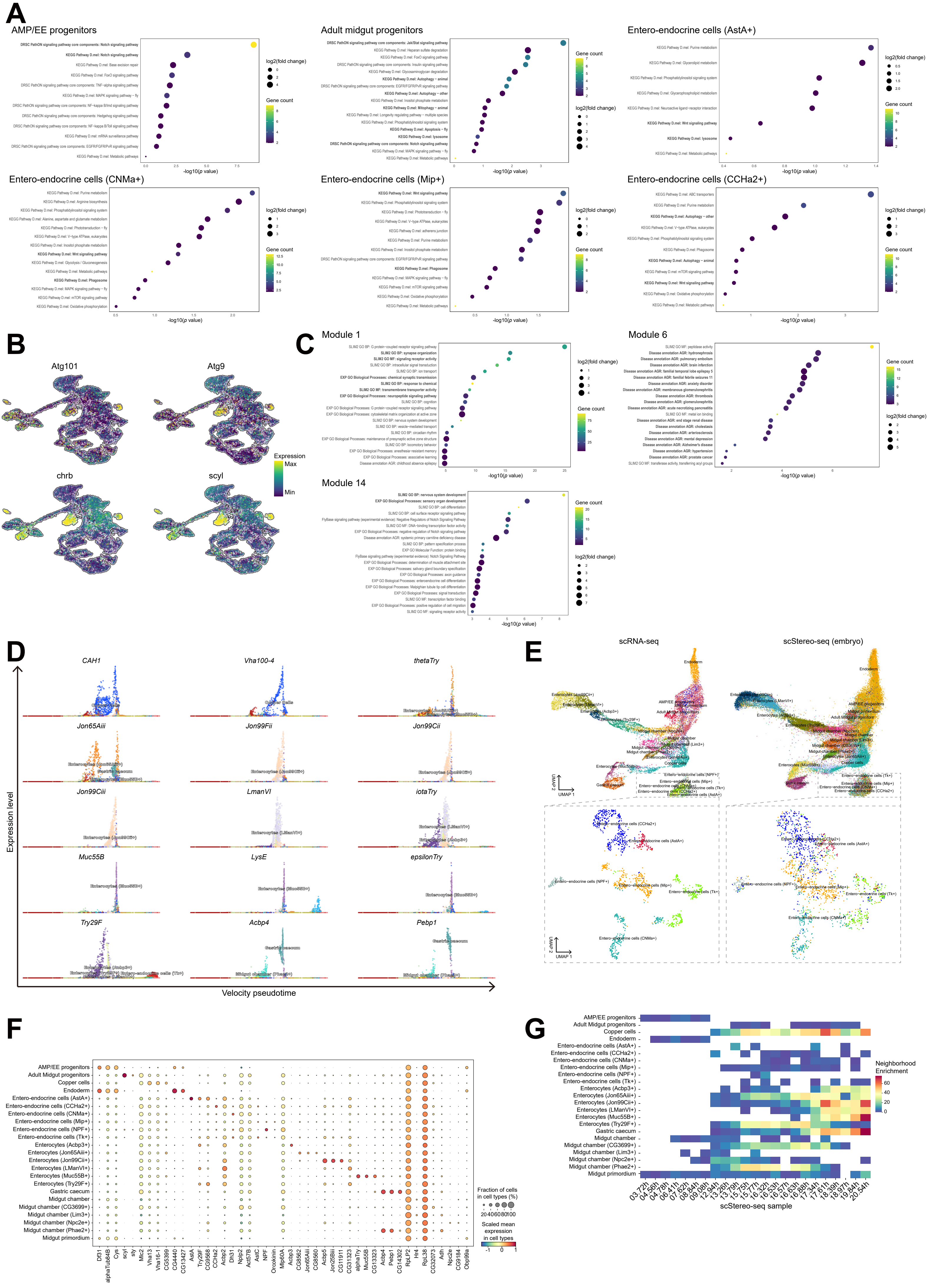
A
AMP/EE progenitors
Adult midgut progenitors
Entero-endocrine cells (AstA+)
Entero-endocrine cells (CNMa+)
Entero-endocrine cells (Mip+)
Entero-endocrine cells (CCHa2+)
log2(fold change)
Gene count
-log10(p value)
B
Atg101
Atg9
chrb
scyl
Expression
Max
Min
C
Module 1
Module 6
Module 14
D
CAH1
Vha100-4
thetaTry
Jon65Aiii
Jon99Fii
Jon99Cii
Jon99Ciii
LmanVI
iotaTry
Muc55B
LysE
epsilonTry
Try29F
Acbp4
Pebp1
Expression level
Velocity pseudotime
E
scRNA-seq
scStereo-seq (embryo)
UMAP 1
UMAP 2
F
Fraction of cells in cell types (%)
Scaled mean expression in cell types
G
Neighborhood Enrichment
scStereo-seq sample

**Figure S10 Diversity of cell types and functions in embryonic midgut. Related to Figure 6.**

**(A)** Bubble plots showing KEGG and DRSC PathON pathway enrichment of midgut cell type marker genes from representative clusters. Pathways in bold are discussed in the main text. **(B)** UMAP plots of midgut scRNA-seq data showing expression level of cell death related genes *Atg101*, *Atg*9, *chrb*, and *scyl*. **(C)** Bubble plots showing gene group enrichment of representative *Hotspot* identified gene modules in midgut scRNA-seq data. Gene group terms in bold are discussed in the main text. **(D)** Dot plots showing relationship between velocity derived pseudotime and expression levels of genes of interest during differentiation of ECs. Each dot represents one cell from midgut scRNA-seq data. **(E)** Co-embedding of midgut (upper) and EE (lower) cells from scRNA-seq and scStereo-seq data in the same UMAP plots. Original scRNA-seq annotations or transferred annotations are labeled. **(F)** Bubble plots showing expression levels and enrichment of cell type top marker genes in label transferred midgut cells in scStereo-seq data. **(G)** Heatmap showing neighborhood enrichment scores of cell types across scStereo-seq samples. Blank cells indicate absence of label transferred cell types or lack of enrichment in corresponding samples.

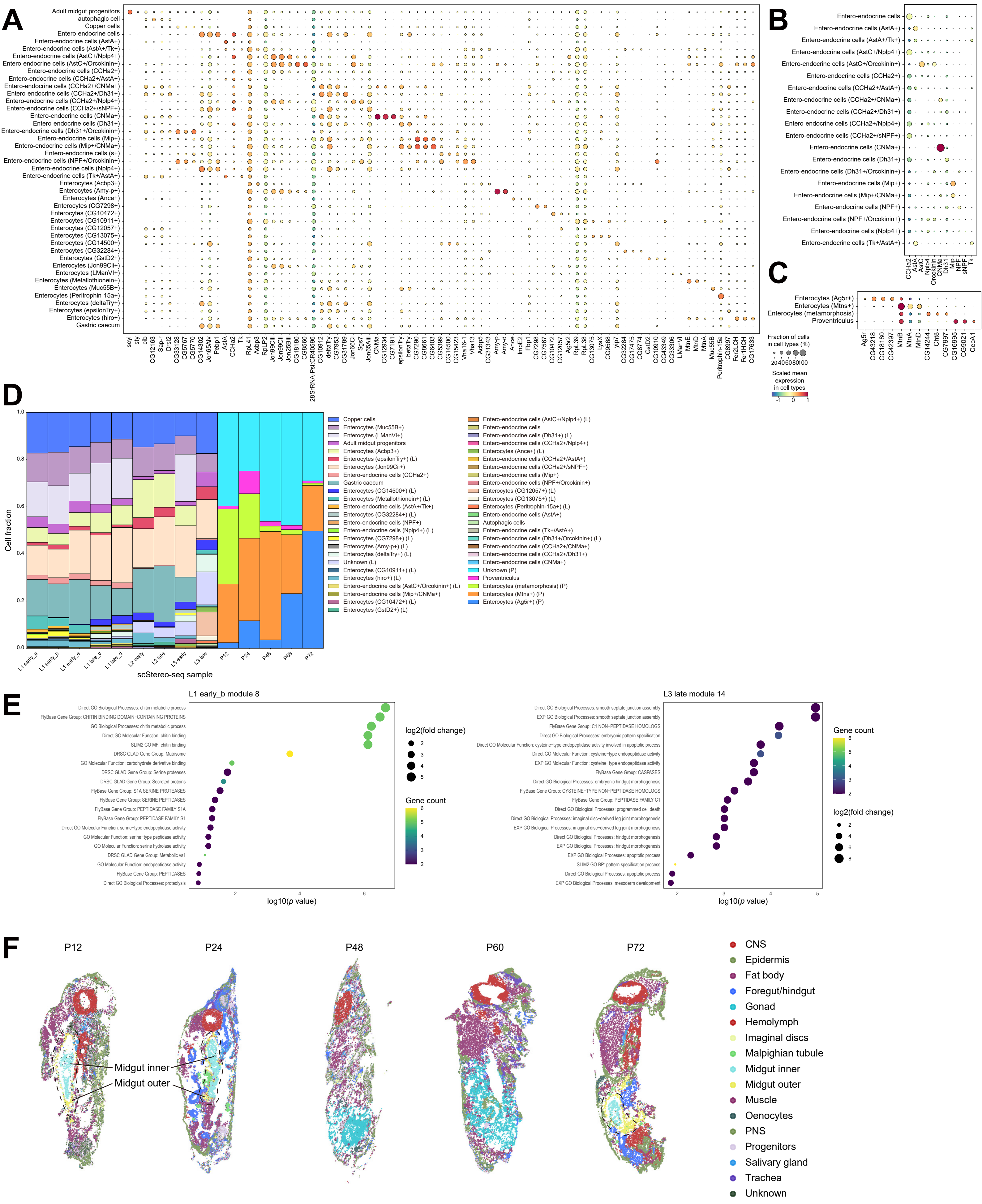
A
Adult midgut progenitors
autophagic cell
Copper cells
Entero-endocrine cells
Entero-endocrine cells (AstA+)
Entero-endocrine cells (AstA+/Tk+)
Entero-endocrine cells (AstC+/Nplp4+)
Entero-endocrine cells (AstC+/Orcokinin+)
Entero-endocrine cells (CCHa2+)
Entero-endocrine cells (CCHa2+/AstA+)
Entero-endocrine cells (CCHa2+/CNMa+)
Entero-endocrine cells (CCHa2+/Dh31+)
Entero-endocrine cells (CCHa2+/Nplp4+)
Entero-endocrine cells (CCHa2+/sNPF+)
Entero-endocrine cells (CNMa+)
Entero-endocrine cells (Dh31+)
Entero-endocrine cells (Dh31+/Orcokinin+)
Entero-endocrine cells (Mip+)
Entero-endocrine cells (Mip+/CNMa+)
Entero-endocrine cells (s+)
Entero-endocrine cells (NPF+/Orcokinin+)
Entero-endocrine cells (Nplp4+)
Entero-endocrine cells (Tk+/AstA+)
Enterocytes (Acbp3+)
Enterocytes (Amy-p+)
Enterocytes (Ance+)
Enterocytes (CG7298+)
Enterocytes (CG10472+)
Enterocytes (CG10911+)
Enterocytes (CG12057+)
Enterocytes (CG13075+)
Enterocytes (CG14500+)
Enterocytes (CG32284+)
Enterocytes (GstD2+)
Enterocytes (Jon99Cii+)
Enterocytes (LManVI+)
Enterocytes (Metallothionein+)
Enterocytes (Muc55B+)
Enterocytes (Peritrophin-15a+)
Enterocytes (deltaTry+)
Enterocytes (epsilonTry+)
Enterocytes (hiro+)
Gastric caecum
scyl sty cib CG12163 Sap-r Drsl2 CG33128 CG5767 CG5770 CG14302 Jon65Aiv Pebp1 AstA CCHa2 Tk RpL41 Acbp3 RpLP2 Jon99Cii Jon99Ciii Jon25Biii CG18180 CG8560 28SrRNA-Psi:CR40596 CG10912 deltaTry CG7953 CG31789 Jon66Ci Sgs7 Jon65Aiii CNMa CG12934 CG7191 epsilonTry betaTry CG7290 CG8661 CG6403 CG5399 CG14500 CG15423 Vha16-1 Vha13 Acbp5 CG31343 Amy-p Amy-d Ance ImpE2 Fbp1 CG7298 CG7567 CG10472 CG12057 Ag5r2 RpL39 RpL38 CG13075 LysX CG9568 yip7 CG32284 CG17475 CG8774 GstD2 CG10910 CG43349 CG33306 LManVI MtnE MtnD MtnA Muc55B Peritrophin-15a CG8997 Fer2LCH Fer1HCH CG17633
B
Entero-endocrine cells
Entero-endocrine cells (AstA+)
Entero-endocrine cells (AstA+/Tk+)
Entero-endocrine cells (AstC+/Nplp4+)
Entero-endocrine cells (AstC+/Orcokinin+)
Entero-endocrine cells (CCHa2+)
Entero-endocrine cells (CCHa2+/AstA+)
Entero-endocrine cells (CCHa2+/CNMa+)
Entero-endocrine cells (CCHa2+/Dh31+)
Entero-endocrine cells (CCHa2+/Nplp4+)
Entero-endocrine cells (CCHa2+/sNPF+)
Entero-endocrine cells (CNMa+)
Entero-endocrine cells (Dh31+)
Entero-endocrine cells (Dh31+/Orcokinin+)
Entero-endocrine cells (Mip+)
Entero-endocrine cells (Mip+/CNMa+)
Entero-endocrine cells (NPF+)
Entero-endocrine cells (NPF+/Orcokinin+)
Entero-endocrine cells (Nplp4+)
Entero-endocrine cells (Tk+/AstA+)
CCHa2 AstA AstC Nplp4 Orcokinin CNMa Dh31 Mip NPF sNPF Tk
C
Enterocytes (Ag5r+)
Enterocytes (Mtns+)
Enterocytes (metamorphosis)
Proventriculus
Ag5r CG43218 CG18180 CG42397 MtnB MtnA MtnD CG14244 Cht8 CG7997 CG16995 CG9021 CecA1
Fraction of cells in cell types (%)
20 40 60 80 100
Scaled mean expression in cell types
-1 0 1
D
Cell fraction
1.0 0.8 0.6 0.4 0.2 0.0
L1 early_a L1 early_b L1 early_e L1 late_c L1 late_d L2 early L2 late L3 early L3 late P12 P24 P48 P68 P72
scStereo-seq sample
Copper cells
Enterocytes (Muc55B+)
Enterocytes (LManVI+)
Adult midgut progenitors
Enterocytes (epsilonTry+) (L)
Enterocytes (Jon99Cii+)
Entero-endocrine cells (CCHa2+)
Gastric caecum
Enterocytes (CG14500+) (L)
Enterocytes (Metallothionein+) (L)
Entero-endocrine cells (AstA+/Tk+) (L)
Enterocytes (CG32284+) (L)
Entero-endocrine cells (NPF+)
Entero-endocrine cells (Nplp4+) (L)
Enterocytes (CG7298+) (L)
Enterocytes (Amy-p+) (L)
Enterocytes (deltaTry+) (L)
Unknown (L)
Enterocytes (CG10911+) (L)
Enterocytes (hiro+) (L)
Entero-endocrine cells (AstC+/Orcokinin+) (L)
Entero-endocrine cells (Mip+/CNMa+) (L)
Enterocytes (CG10472+) (L)
Enterocytes (GstD2+) (L)
Entero-endocrine cells (AstC+/Nplp4+) (L)
Entero-endocrine cells
Entero-endocrine cells (Dh31+) (L)
Entero-endocrine cells (CCHa2+/Nplp4+)
Enterocytes (Ance+) (L)
Entero-endocrine cells (CCHa2+/AstA+)
Entero-endocrine cells (CCHa2+/sNPF+)
Entero-endocrine cells (Mip+)
Entero-endocrine cells (NPF+/Orcokinin+)
Enterocytes (CG12057+) (L)
Enterocytes (CG13075+) (L)
Enterocytes (Peritrophin-15a+) (L)
Entero-endocrine cells (AstA+)
Autophagic cells
Entero-endocrine cells (Tk+/AstA+)
Entero-endocrine cells (Dh31+/Orcokinin+) (L)
Entero-endocrine cells (CCHa2+/CNMa+)
Entero-endocrine cells (CCHa2+/Dh31+)
Entero-endocrine cells (CNMa+)
Unknown (P)
Proventriculus
Enterocytes (metamorphosis) (P)
Enterocytes (Mtns+) (P)
Enterocytes (Ag5r+) (P)
E
L1 early_b module 8
Direct GO Biological Processes: chitin metabolic process
FlyBase Gene Group: CHITIN BINDING DOMAIN-CONTAINING PROTEINS
GO Biological Processes: chitin metabolic process
Direct GO Molecular Function: chitin binding
SLIM2 GO MF: chitin binding
DRSC GLAD Gene Group: Matrisome
GO Molecular Function: carbohydrate derivative binding
DRSC GLAD Gene Group: Serine proteases
DRSC GLAD Gene Group: Secreted proteins
FlyBase Gene Group: S1A SERINE PROTEASES
FlyBase Gene Group: SERINE PEPTIDASES
FlyBase Gene Group: PEPTIDASE FAMILY S1A
FlyBase Gene Group: PEPTIDASE FAMILY S1
Direct GO Molecular Function: serine-type endopeptidase activity
GO Molecular Function: serine-type peptidase activity
GO Molecular Function: serine hydrolase activity
DRSC GLAD Gene Group: Metabolic vs1
GO Molecular Function: endopeptidase activity
FlyBase Gene Group: PEPTIDASES
Direct GO Biological Processes: proteolysis
2 4 6
log10(p value)
log2(fold change)
2 3 4 5
Gene count
6 5 4 3 2
L3 late module 14
Direct GO Biological Processes: smooth septate junction assembly
EXP GO Biological Processes: smooth septate junction assembly
FlyBase Gene Group: C1 NON-PEPTIDASE HOMOLOGS
Direct GO Biological Processes: embryonic pattern specification
Direct GO Molecular Function: cysteine-type endopeptidase activity involved in apoptotic process
Direct GO Molecular Function: cysteine-type endopeptidase activity
EXP GO Molecular Function: cysteine-type endopeptidase activity
FlyBase Gene Group: CASPASES
Direct GO Biological Processes: embryonic hindgut morphogenesis
FlyBase Gene Group: CYSTEINE-TYPE NON-PEPTIDASE HOMOLOGS
FlyBase Gene Group: PEPTIDASE FAMILY C1
Direct GO Biological Processes: programmed cell death
Direct GO Biological Processes: imaginal disc-derived leg joint morphogenesis
EXP GO Biological Processes: imaginal disc-derived leg joint morphogenesis
Direct GO Biological Processes: hindgut morphogenesis
EXP GO Biological Processes: hindgut morphogenesis
EXP GO Biological Processes: apoptotic process
SLIM2 GO BP: pattern specification process
Direct GO Biological Processes: apoptotic process
EXP GO Biological Processes: mesoderm development
2 3 4 5
log10(p value)
Gene count
6 5 4 3 2
log2(fold change)
2 4 6 8
F
P12 P24 P48 P60 P72
Midgut inner
Midgut outer
CNS
Epidermis
Fat body
Foregut/hindgut
Gonad
Hemolymph
Imaginal discs
Malpighian tubule
Midgut inner
Midgut outer
Muscle
Oenocytes
PNS
Progenitors
Salivary gland
Trachea
Unknown

**Figure S11 Diversity of cell types and functions in larval and pupal midgut. Related to Figure 6.**

**(A-C)** Bubble plots showing expression level and enrichment of top marker genes of **(A)** all larval midgut, **(B)** larval entero-endocrine cells, and **(C)** all pupal midgut cell types in scStereo-seq data. **(D)** Bar plot showing cell type composition of larval and pupal midgut in scStereo-seq samples. Annotations with (L) or (P) indicate clusters identified only in larva or pupa samples, respectively. **(E)** Bubble plots showing gene group enrichment of representative *Hotspot* identified gene modules in larval midgut scStereo-seq data. **(F)** Clustering and annotation results of 2D spatial transcriptomes of representative pupa scStereo-seq sample sections. Clusters annotated as “midgut inner” and “midgut outer” are highlighted. Samples are not on the same scale.

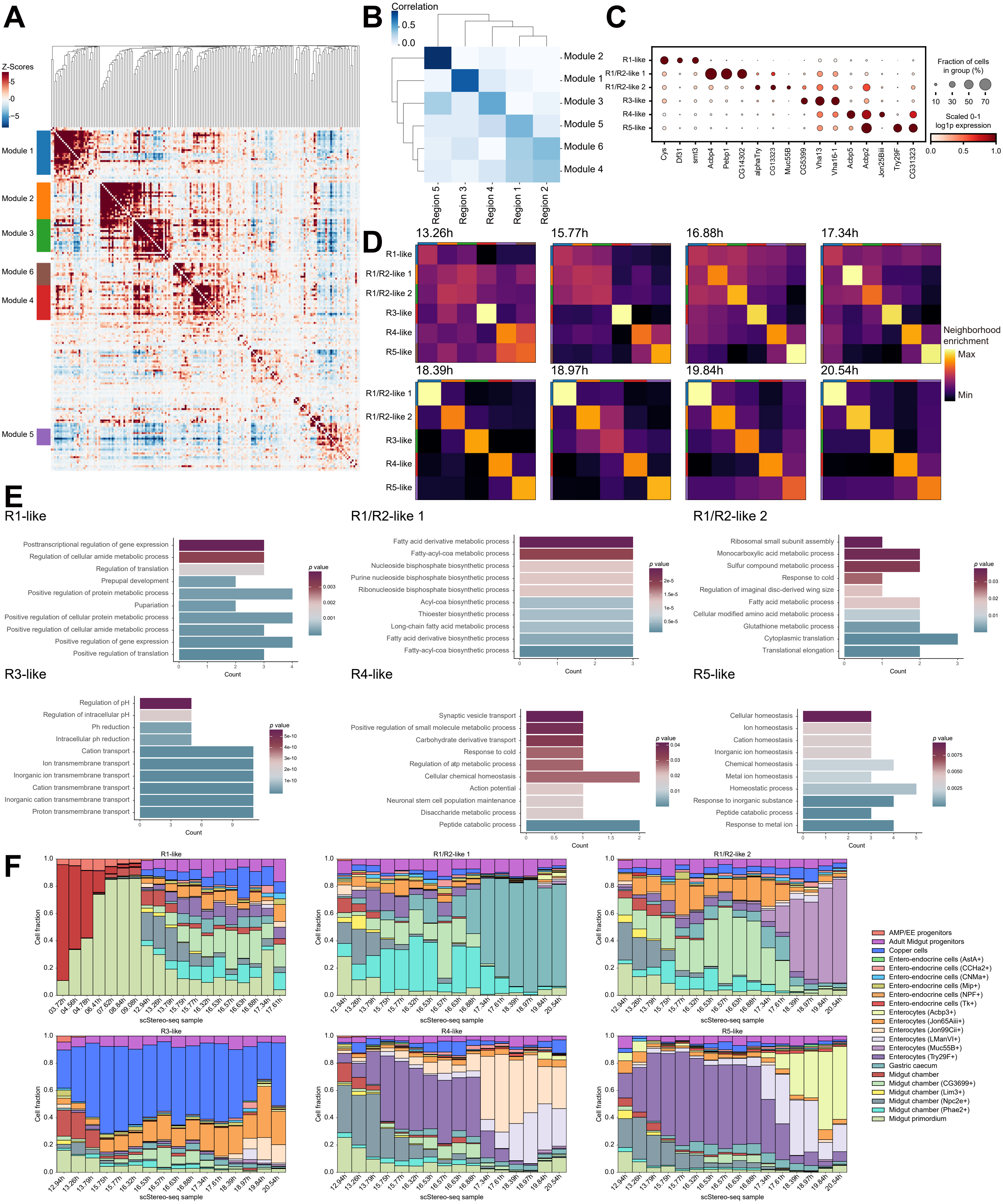
A
Z-Scores
Module 1
Module 2
Module 3
Module 6
Module 4
Module 5
B
Correlation
Module 2
Module 1
Module 3
Module 5
Module 6
Module 4
Region 5
Region 3
Region 4
Region 1
Region 2
C
R1-like
R1/R2-like 1
R1/R2-like 2
R3-like
R4-like
R5-like
Cys
Df31
smt3
Acbp4
Pebp1
CG14302
alphaTry
CG13323
Muc55B
CG5399
Vha13
Vha16-1
Acbp5
Acbp2
Jon25Biii
Try29F
CG31323
Fraction of cells in group (%)
Scaled 0-1 log1p expression
D
13.26h
15.77h
16.88h
17.34h
18.39h
18.97h
19.84h
20.54h
Neighborhood enrichment
Max
Min
E
R1-like
Posttranscriptional regulation of gene expression
Regulation of cellular amide metabolic process
Regulation of translation
Prepupal development
Positive regulation of protein metabolic process
Pupariation
Positive regulation of cellular protein metabolic process
Positive regulation of cellular amide metabolic process
Positive regulation of gene expression
Positive regulation of translation
R1/R2-like 1
Fatty acid derivative metabolic process
Fatty-acyl-coa metabolic process
Nucleoside bisphosphate biosynthetic process
Purine nucleoside bisphosphate biosynthetic process
Ribonucleoside bisphosphate biosynthetic process
Acyl-coa biosynthetic process
Thioester biosynthetic process
Long-chain fatty acid metabolic process
Fatty acid derivative biosynthetic process
Fatty-acyl-coa biosynthetic process
R1/R2-like 2
Ribosomal small subunit assembly
Monocarboxylic acid metabolic process
Sulfur compound metabolic process
Response to cold
Regulation of imaginal disc-derived wing size
Fatty acid metabolic process
Cellular modified amino acid metabolic process
Glutathione metabolic process
Cytoplasmic translation
Translational elongation
R3-like
Regulation of pH
Regulation of intracellular pH
Ph reduction
Intracellular ph reduction
Cation transport
Ion transmembrane transport
Inorganic ion transmembrane transport
Cation transmembrane transport
Inorganic cation transmembrane transport
Proton transmembrane transport
R4-like
Synaptic vesicle transport
Positive regulation of small molecule metabolic process
Carbohydrate derivative transport
Response to cold
Regulation of atp metabolic process
Cellular chemical homeostasis
Action potential
Neuronal stem cell population maintenance
Disaccharide metabolic process
Peptide catabolic process
R5-like
Cellular homeostasis
Ion homeostasis
Cation homeostasis
Inorganic ion homeostasis
Chemical homeostasis
Metal ion homeostasis
Homeostatic process
Response to inorganic substance
Peptide catabolic process
Response to metal ion
Count
p value
F
Cell fraction
scStereo-seq sample
AMP/EE progenitors
Adult Midgut progenitors
Copper cells
Entero-endocrine cells (AstA+)
Entero-endocrine cells (CCHa2+)
Entero-endocrine cells (CNMa+)
Entero-endocrine cells (Mip+)
Entero-endocrine cells (NPF+)
Entero-endocrine cells (Tk+)
Enterocytes (Acbp3+)
Enterocytes (Jon65Aiii+)
Enterocytes (Jon99Cii+)
Enterocytes (LManVI+)
Enterocytes (Muc55B+)
Enterocytes (Try29F+)
Gastric caecum
Midgut chamber
Midgut chamber (CG3699+)
Midgut chamber (Lim3+)
Midgut chamber (Npc2e+)
Midgut chamber (Phae2+)
Midgut primordium

**Figure S12 Functional regionalization of embryonic midgut. Related to Figure 6.**

**(A)** Heatmap showing correlation of functional gene modules identified by *Hotspot* from adult midgut region marker genes in scStereo-seq data. Each row and each column represent a module marker gene, and *Z*-score indicates their correlation. **(B)** Heatmap showing correlation between gene modules in **(A)** and adult midgut region markers. **(C)** Bubble plot showing expression level and enrichment of top marker genes of embryonic midgut regions. **(D)** Heatmaps showing neighborhood enrichment scores of midgut regions across scStereo-seq samples. **(E)** Bar plots showing GO enrichment of marker genes in regions identified in embryonic midgut. **(F)** Bar plots showing cell type composition of midgut regions in scStereo-seq samples. Cell types are label transferred from scRNA-seq data.

### SUPPLEMENTAL TABLES

**Table S1 scStereo-seq, scRNA-seq, and scATAC-seq samples covered in this study and data quality control (3 worksheets). Related to Figure 1 (separate spreadsheet).**

Number ranges in original embryo sample names indicate collection time window, e.g., E8-10h indicates embryos collected 8 to 10 h from egg laying. We did not obtain quality P36 scStereo-seq and E6-8h scRNA-seq data. For the latter, based on UMAP plot and *RAPToR* inference, we believe that the rest of the samples have achieved extensive coverage of cells at all developmental ages.

Worksheet 1: quality control stats for scStereo-seq data.

Worksheet 2: quality control stats for scRNA-seq data.

Worksheet 3: quality control stats for scATAC-seq data.

**Table S2 Lists of top marker genes and annotations of clusters generated in scStereo-seq, scRNA-seq, scATAC-seq, and integrated scRNA-seq and scATAC-seq data (8 worksheets). Related to Figures 1 and 2 (separate spreadsheet).**

Worksheet 1: Top marker genes for clustering of scStereo-seq data.

Worksheet 2: Cluster annotations of scStereo-seq data.

Worksheet 3: Top marker genes for coarse clustering of scRNA-seq data.

Worksheet 4: Coarse cluster annotations of scRNA-seq data.

Worksheet 5: Top marker genes for coarse clustering of scATAC-seq data.

Worksheet 6: Coarse cluster annotations of scATAC-seq data.

Worksheet 7: Top marker genes for clustering of integrated scRNA-seq and scATAC-seq data.

Worksheet 8: Cluster annotations of integrated scRNA-seq and scATAC-seq data.

**Table S3 A list of genes without reported spatial expression patterns and their scStereo-seq inferred spatial expression patterns. Related to Figure 1 (separate spreadsheet).**

A list of 338 genes without reported spatial expression patterns. Cell type and/or tissue enrichment inferred from scStereo-seq data are listed if available, with number of occurrences among Stereo-seq samples labeled in brackets. Spatial gene expression patterns generated from integrated scStereo-seq and scRNA-seq data are presented as projections along the Z-axis. All Stereo-seq samples are shown in lateral view with the anterior side facing left.

**Table S4 Lists of marker genes and annotations of subclusters generated from scRNA-seq tissue subclusters, scATAC-seq tissue subclusters, integrated scRNA-seq and scATAC-seq CNS subclusters, and scStereo-seq midgut subclusters (9 worksheets). Related to Figures 2, 5, and 6s (separate spreadsheet).**

Worksheet 1: Top marker genes for tissue subclustering of scRNA-seq data.

Worksheet 2: Tissue subcluster annotations of scRNA-seq data.

Worksheet 3: Top marker genes for tissue subclustering of scATAC-seq data.

Worksheet 4: Tissue subcluster annotations of scATAC-seq data.

Worksheet 5: Top marker genes for CNS subclustering of integrated scRNA-seq and scATAC-seq data.

Worksheet 6: CNS subcluster annotations of integrated scRNA-seq and scATAC-seq data.

Worksheet 7: Top pairwise marker genes for CNS mature neuron subtypes of integrated scRNA-seq and scATAC-seq data.

Worksheet 8: Top marker genes for larval and pupal midgut subclustering of scStereo-seq data.

Worksheet 9: Larval and pupal midgut subcluster annotations of scStereo-seq data.

**Table S5 A list of tissue substructure/cell type marker genes identified in both scRNA-seq and scATAC-seq datasets. Related to Figure 2 (separate spreadsheet).**

**Table S6 Top 100 *CytoTRACE* genes positively correlated with *CytoTRACE* score and their overlap in three germ layers. Related to Figure 3.**

| Ectoderm | Mesoderm | Endoderm | Gene count | CytoTRACE genes |
|---|---|---|---|---|
| True | True | True | 21 | Tctp, eEF1alpha1, eEF1gamma, RpS3A, RpS2, RpL6, Fkbp39, Ran, dUTPase, Set, akirin, RpL23A, Hel25E, RpS24, Nlp, janA, His2Av, smt3, Hsc70-4, Nph, Nop60B |
| False | True | True | 18 | tsr, Df31, CNBP, B52, Non2, HmgD, Fkbp12, yps, bic, Fib, SC35, D1, Cyp1, Bacc, SmB, geminin, Mapmodulin, Roc1a |
| True | False | True | 46 | eEF1beta, RpL41, RpS20, RpL35, RpL27A, RpL10, RpL35A, RpS19a, Inx3, RpL13, RpL27, sta, RpL13A, RpL32, RpS30, RpL37A, RpLP1, RpL37a, Inx2, RpS16, RpS27, RpL23, RpL30, RpL14, eEF5, RpL4, RpL7, RpL34b, RpL18A, RpS26, RpL3, RpS11, RpS7, RpS10b, RpS8, RpS18, RpL22, RpS9, RpL18, RpS17, RpS25, Rack1, RpL19, RpL31, RpS28b, RpL36 |
| False | False | True | 15 | kra, RpL15, RpL24, chic, Rbp1, RpS6, lolal, RpS15, Ip259, SNRPG, RpL17, His4r, RpS4, RpS13, arm |
| True | True | False | 12 | RpS5b, Tom, kuk, CycB, CycA, borr, polo, pre-lola-G, msb1l, fzy, CG13096, Su(var)205 |
| False | True | False | 49 | alphaTub84B, sqd, Lam, Pep, betaTub56D, l(3)neo38, Ref1, Hrb87F, Hrb98DE, His3.3A, mod, Rm62, tsu, nop5, cass, pzg, lark, SelD, Nop56, HnRNP-K, Ama, rump, CG1943, mago, lwr, kis, SF2, x16, rin, lncRNA:CR45184, Chrac-16, CG7483, NHP2, SmD2, HmgZ, SmE, RnpS1, Ntf-2, Ndf, CG30122, Cdk1, CCT4, hth, Eb1, alphaTub84D, Nopp140, Top2, PIG-B, lig |
| True | False | False | 21 | CG4440, Cys, sala, CG13427, link, RpL36A, CG8960, RpL9, RpL21, eEF2, PCNA, RpS27A, RpLP2, RpL7A, CG10035, RpS21, Mcm7, CycB3, RnrS, RpL26, RpL12 |

**Table S7 A list of transcription factor motifs enriched in tissue types in three germ layers and CNS. Related to Figures 4 and 5 (separate spreadsheet).**

**Table S8 Lists of genes associated with CNS and midgut morphological changes based on *Spateo* morphometric analysis (2 worksheets). Related to Figures 5 and 6 (separate spreadsheet).**

For parameters describing vector fields in *Spateo* morphometric analysis: acceleration refers to the cell migration velocity at a certain point, which can serve as an indicator of cell migration distance during cell state transition; Curvature describes how a spatial path of cell migration bends, which can reflect the smoothness of the path during cell state transition; Curl and torsion are two additional measurements of vector fields, describing the degree of rotation and curve twisting in the field, respectively. They can provide additional insights into gene functions on cell migration paths.

**Table S9 A list of primers for generating FISH probes in this study. Related to Figures 1, 2, 4, and 5 (separate spreadsheet).**

**Table S10 Hex codes in palettes for color coding of tissue and cell type clusters (2 worksheets). Related to all figures (separate spreadsheet).**

#### SUPPLEMENTAL MOVIES

**Movie S1 Vector fields of embryonic CNS morphometric changes across 7 scStereo-seq samples. Related to Figure 5.**

Color scales indicate acceleration scores of CNS cell bins.

**Movie S2 Vector fields of embryonic midgut morphometric changes across 7 scStereo-seq samples. Related to Figure 6.**

Color scales indicate acceleration scores of midgut cell bins.

### SUPPLEMENTAL DATA

**Data S1 Unsupervised clustering results of scStereo-seq samples covered in this study. Related To Figure 1.**

2D representation of cell bin clusters of all available sections are shown. Cell bins from all available sections were combined for clustering. Available through Mendeley Data (https://doi.org/10.17632/4zf847bxcd.1).

**Data S2 Interactive 3D UMAP plots of integrated scRNA-seq and scATAC-seq data of down sampled all cells and separated three germ layers. Related to Figure 2.**

**Data S3 Interactive 3D UMAP plots of CNS scRNA-seq, scATAC-seq, and integrated data. Related to Figure 5.**

### METHODS

#### METHOD DETAILS

##### Stereo-seq library preparation and sequencing

**Tissue processing and imaging.** Stereo-seq library was prepared following the protocol in Ref[1] with modification. Unless otherwise mentioned, buffer recipes and primer sequences used in this study were the same as Ref[1]. Briefly, embedded samples were balanced at -20 ℃ for 30 min and sectioned using a Leica CM1950 cryostat at a thickness of 7 or 8 μm. All available resulting slices were collected for sequencing. To minimize batch effects, six embryo/L1 slices, or four L2 slices, or two L3/pupa slices were mounted on one 1 cm × 1 cm Stereo-seq chip simultaneously. The mounted chips were then incubated at 37 ℃ for 3 min and subsequently fixed in cold methanol (Sigma, 34860) at -20 ℃ for 30 min. After fixation, chips were treated with staining solution consisted of 0.1 × SSC (Ambion, AM9770), 1/200 nucleic acid dye (Invitrogen, Q10212), and 2 U/μl RNase inhibitor (NEB, M0314L). Sections were incubated in staining solution for 3 min and then rinsed with 0.1 × saline sodium citrate (SSC) buffer supplemented with 2 U/μl RNase inhibitor. Sections were then imaged using a Motic Custom PA53 FS6 microscope (10 × objective) before RNA capture.

**RNA capture and *in situ* reverse transcription.** Tissue slices, mounted on Stereo-seq chips, were subjected to treatment with 0.1% pepsin (Sigma, P7000) in a 0.01 M HCl solution, followed by incubation at 37 ℃ for 6 min for permeabilization. Permeabilized slices were washed twice using 0.1 × SSC buffer supplemented with 0.05 U/ml RNase inhibitor. After permeabilization, RNAs were released from the tissue and captured by DNA nanoballs (DNBs) on Stereo-seq chips. Chips were incubated in reverse transcription mixture at 42 ℃ overnight. After reverse transcription, chips were washed twice with 0.1 × SSC buffer and subsequently incubated in tissue removal buffer at 37 ℃ for 30 min. cDNAs were collected by treating the chip with Exonuclease I (NEB, M0293L) for 1 h at 37 ℃. Residual cDNAs were collected through a final rinse of the chip with 0.1× SSC buffer. The flowthrough, combined with the Exonuclease I treated product, was purified using 0.8 × VAHTS DNA clean beads (Vazyme, N411-03).

**Library construction and sequencing.** Purified cDNAs were amplified using the

KAPA HiFi Hotstart Ready Mix (Roche, KK2602) with 0.8 μM cDNA-PCR primer. PCR amplification was performed with the following steps: pre-heating at 95 ℃ for 5 min; 15 amplification cycles with 98 ℃ for 20 s, 58 ℃ for 20 s, and 72 ℃ for 3 min; final incubation at 72 ℃ for 5 min. Amplified cDNAs were purified using 1 × VAHTS DNA clean beads. For library construction, 20 ng cDNAs were fragmented using in-house Tn5 transposase at 55 ℃ for 10 min. 0.02% SDS was added to terminate the reaction. Fragmented cDNAs were then subjected to amplification using the KAPA HiFi Hotstart Ready Mix with the addition of 3 μL each 10 μM Stereo-seq-Library-F primer and Stereo-seq-Library-R primer. PCR amplification was performed with the following steps: pre-heating at 95℃ for 5 min; 13 amplification cycles with 98 ℃ for 20 s, 58 ℃ for 20 s, and 72 ℃ for 30 s; final incubation at 72 ℃ for 5 min. Amplified PCR products were purified using VAHTS DNA clean beads. Construction of sequencing libraries and sequencing with MGI DNBSEQ-T10 sequencer were performed following manufacturer's protocols.

**scRNA-seq library construction and sequencing**

Sequencing libraries were prepared using the DNBelab C Series High-throughput Single-Cell RNA Library Preparation Kit (MGI, 940000004700) following manufacturer's protocol. Briefly, fixed single cell suspensions, preserved in methanol, were balanced at 4 ℃ for 5 min and centrifuged at 2,000 g at 4℃ for 5 min. Cell pellets were washed twice with wash buffer [PBS with 1% RNAse inhibitor and 0.04% bovine serum albumin (BSA)] before resuspension in 100 μL wash buffer. Cells were counted and 20,000 cells were aliquoted for droplet generation, emulsion breakage, beads collection, reverse transcription, second strand synthesis, cDNA amplification, and droplet index product amplification to generate barcoded libraries. Sequencing libraries were quantified using Qubit ssDNA Assay Kit (Invitrogen, Q10212) and subsequently sequenced with MGI DNBSEQ-T10 sequencer.

**scATAC-seq library construction and sequencing**

Sequencing libraries were prepared using the DNBelab C Series Single-Cell ATAC Library Prep set (MGI, 1000021878) following the manufacturer's protocol. Briefly, single nuclei were extracted by grinding flash-frozen embryos in the lysis buffer using a 2 mL homogenizer (Sigma-Aldrich, D8938). Nuclei were washed twice with wash buffer and resuspended in 100 μL wash buffer. Nuclei were counted before 50,000 nuclei were aliquoted and subjected to in-house Tn5 transposase treatment. Treated single-nucleus suspension was then converted to barcoded scATAC-seq libraries through droplet encapsulation, pre-amplification, emulsion breakage, beads collection, DNA amplification, and purification. Indexed sequencing libraries were constructed following the manufacturer's protocol and quantified using Qubit ssDNA Assay kit. Sequencing libraries were subsequently sequenced with MGI DNBSEQ-T10 sequencer.

**scStereo-seq data processing**

**Cell segmentation.** Manual registration of the DNB image with the nucleic acid staining image was performed as described in Ref[1]. Nucleus identification and cell segmentation was performed on the aligned images using *Cellpose*[2]. Additionally, *spateo.cs.expand_labels* function from *Spateo*[3] was applied to augment each segmented nucleus with an additional 10 DNBs, approximating the actual size of *Drosophila* cells. Furthermore, *spateo.io.read_bgi* function was employed to perform cell segmentation on the aligned bin1 matrix. Within each segmented cell, UMI counts from all DNBs corresponding to the segmentation were aggregated, preserving the counts on a per-gene basis. These aggregated counts were then summed to generate a gene/cell matrix for downstream analysis. To accurately determine the centroid of each cell, *rearrr* package[4] was used to facilitate the identification of the central position within the segmented cell.

**Section alignment.** *spateo.align.morpho_align* function was used to perform registration on all slices of each sample, which involved aligning and co-registering all slices within a sample to ensure accurate spatial alignment throughout the sample volume.

**Data quality control.** Cell bins were first filtered to retain those with mitochondrial gene content ≤ 10%. Cell bins were further filtered with *filter_cells* function from the *spateo.pp.filter* module (*min_area=20*, *min_expr_genes=20*). Gene filtering was applied with *filter_genes* function from the same module (*min_cells=3*, *min_counts*=1). *spateo.tl.pearson_residuals* function was then used to normalize data from all slices from the same sample. Normalized data were combined to create an integrated matrix for each sample.

**Dimension reduction and clustering.** *dynamo.tl.compute_neighbors* function was first used to compute a neighbor graph with parameters *n_neighbors=6* and *n_pca_components=50*. Dimensionality reduction was performed using *spateo.tl.pca_spateo* function with parameter *n_pca_components=50*, along with the *spateo.tl.reduceDimension* function from the *dyn.tl* module with parameter *n_components*=2. Cluster assignment was performed using *spateo.tl.scc* function. Marker genes for each cluster were determined using *scanpy.tl.rank_genes_groups* function from *SCANPY*[5] with *t-test* method. Based on marker genes and spatial morphology, clusters were manually curated and annotated into cell types, tissues, and germ layers.

**3D model reconstruction and alignment.** *Spateo* was employed to reconstruct 3D models of spatial transcriptomic data. Specifically, 3D point-cloud models of scStereo-seq samples were generated using *spateo.tdr.construct_pc* function with default parameters. To align spatial coordinates for continuous morphometric analysis, the scStereo-seq sample of the first time point was set as the reference by transforming the coordinates of the sample through *spateo.tdr.rotate_model* function, such that the centroid of the embryo was located at origin. The anterior-posterior axis and dorsal-ventral axis corresponded to the x and y axis in the coordinate system, respectively. *spateo.tl.model_align* function was used with default parameters to align all other sample models to the same coordinate system. Finally, 3D mesh models of samples were created through *spateo.tdr.construct_surface* function with parameters adjusted for each sample. 3D models of individual tissues were generated with the same methods as those for the entire embryo.

**scRNA-seq data processing**

**Construction of gene/cell matrix.** Raw reads generated by DNBelab C4-based scRNA-seq were processed as previously described[6]. Briefly, reads were pre-processed with *PISA*[7] and aligned to the *Drosophila melanogaster* genome (dm6) using *STAR*[8]. Gene expression levels were quantified with *PISA* to generate gene/cell matrix, and genes present in < 3 cells were filtered.

**Data quality control.** Cells were filtered by the following criteria: feature number > 500 and mitochondrial gene content ≤ 10%. *DoubletFinder*[9] was used to identify and remove doublets, with a doublet anticipation of 3%. Batch effects were corrected with *scVI*[10].

**Dimension reduction, clustering, and annotation.** *Seurat*[11] was used for data normalization and dimension reduction. Louvain method was used for cell clustering. Based on marker genes, clusters were manually curated and annotated into cell types, tissues, and germ layers. For tissue subclustering, annotated tissues were divided into their respective matrices and subjected to *RunUMAP* function, followed by re-clustering at different resolutions (see **Table S4**) to identify detailed cell subtypes. Based on marker genes, subclusters were manually curated and annotated into cell subtypes. For all plots, hex color codes of tissues can be found in **Table S10**.

**scATAC-seq data processing**

**Construction of peak/cell matrix.** Raw reads generated by DNBelab C4-based scATAC-seq were pre-processed with *PISA*. Retained reads were then aligned to the *Drosophila melanogaster* genome (dm6) using *BWA*[12] with default parameters to generate BAM files. Subsequently, fragment files from the same scATAC-seq library were created using *bap2*[13]. Processed fragments were used for peak calling with *MACS2*[14] for each sample, including only autosome peaks. To minimize batch effects, samples within the same time window were matched pairwise and peak integration was performed with *IDR*[15]. IDR integrated peaks were then merged using *BEDTools*[16]. The resulting merged peak list per time window was used to construct the peak/cell matrix for each time window. Peaks present in < 1% cells were filtered. Further analysis was carried out using *Signac*[17].

**Data quality control.** Cells were filtered by the following criteria: peak region fragment number > 4000 and < 40000; reads percentage in peaks ≥ 15%; blacklist ratio < 0.05; TSS enrichment score > ~1.7 (adjusted for each time window); reads per peak (reads per cell divided by peak number per cell) > 1.5 - 2.1 (adjusted for each time window); nucleosome signal < 4. Doublets were removed through a modified *scrublet* algorithm[18]. After calculation of doublet scores, cells above the 95th percentile were retained. To assess the quality of our data, peaks from our data were juxtaposed with annotated TSS sites (2 kb upstream and 200 bp downstream of TSS)[19], peaks from previous *Drosophila* embryo scATAC-seq datasets[20], curated sets of known embryonic enhancers[21–23], and bulk DHS peaks[24]. For each pair of datasets, the percentage of elements in one set that overlapped with elements in the other set was calculated, permitting a minimum overlap of 1 bp, and reciprocally.

**Dimension reduction and clustering.** Dimension reduction and clustering were performed following pipelines described in Ref[18]. Briefly, peak/cell matrix was normalized using latent semantic indexing with log-scaled term frequency. The 2$^{nd}$ to the 50$^{th}$ principal components were retained after running *RunPCA* function from *Seurat* and L2-normalization was applied to these components. To minimize batch effects, *Harmony*[25] was applied on the PCA matrix prior to running *RunUMAP* function from *Seurat* (*min.dist = 0.3*) for clustering. Louvain clustering algorithm was used and cluster resolution for each time window was set to 0.3. Two rounds of dimension reduction and clustering were performed for each time window. In both rounds, clusters with a cell count < 5% of total cell count and clusters dominated by cells from a single batch (> 80%) were discarded. After the first round of clustering, the retained clusters were used in cluster-specific peak re-calling to generate a new peak/cell matrix, including only autosome peaks. The resulting clusters were then re-clustered and selected for the final whole embedding.

**Integrative UMAP embedding.** To create an integrative peak list encompassing samples from all time windows, peak profiles from all 11 time windows after the peak re-calling were merged using *BEDTools*. With this peak list, 11 chromatin accessibility objects were created, containing retained cells after two rounds of clustering. These objects were then merged to generate a single chromatin accessibility profile. Dimension reduction and clustering were performed as described above with some modifications, including removal of peak representation filter. Clusters with < 100 cells, instead of clusters with a cell count < 5% of total cell count, were excluded. Additionally, *Runharmony* function from *Harmony* was not run on the integrative matrix to preserve differences between time windows, which may reflect epigenetic changes during development.

**Tissue and cell type annotation.** To annotate cell types at the tissue level, a gene activity matrix was created using *GeneActivity* function from *Signac*, which computed the counts in both gene body and promoter region (2 kb upstream of TSS) to generate a gene/cell matrix. Marker genes for each cluster were identified using the *FindAllMarkers* function from *Seurat* with the parameters *min.pct = 0.1* and *logfc.threshold = 0.1*. Based on marker genes, clusters were manually curated and annotated into cell types, tissues, and germ layers **(Table S4)**. Tissue subclustering was performed as described for scRNA-seq data.

**Developmental age inference**

Inference of developmental age for each cell in scRNA-seq and scStereo-seq data was performed by *RAPToR Drosophila* reference. For scRNA-seq data, normalized gene expression matrices of single cells were used as input; for scStereo-seq data, pseudo bulk expression matrices of entire embryo samples were used as input. Overlapping genes between our data and reference data were used for age inference. *ae* function was used to generate age predictions from input.

Inference of developmental age for each cell in scATAC-seq data was performed following a neural network-based algorithm described in Ref[18]. The peak/cell matrix described above was used to train a new model from our own dataset. In brief, peak/cell matrix was divided into 11 partitions and 10 of them was used for model training, reserving 1 partition for testing. The midpoint of each time window was considered as the developmental age. The constructed model was then used to infer developmental age of all cells.

**Integration of scRNA-seq and scStereo-seq data**

**Gene expression imputation.** *NovoSpaRc*[26] was used for gene expression imputation in scStereo-seq data based on scRNA-seq data. Single cells within 1 h difference of inferred age in scRNA-seq data were used for imputation of each scStereo-seq sample. Gene expression matrices from scRNA-seq data were used as the primary input for *NovoSpaRc* and scStereo-seq gene expression matrices were used as reference atlas. Optimal transport of cells to their spatial locations was computed with the following parameters: *alpha_linear = 0.8, epsilon = 5e-3*.

**Label transfer of scRNA-seq annotation.** After dimensionality reduction, top 20 dimensions of scRNA-seq data were used for label transfer. Canonical correlation analysis (CCA) was performed using *FindTransferAnchors* function from *Seurat* to identify corresponding anchors between scRNA-seq and scStereo-seq data. *TransferData* function was used to transfer annotations from scRNA-seq data to scStereo-seq data. *MapQuery* function was used to integrate embeddings of scRNA-seq and scStereo-seq data and project them in the same UMAP space for visualization. For midgut of larval samples (except L3 late), label transferred and re-annotated embryo scStereo-seq midgut cell bins, instead of scRNA-seq midgut cells, were used as the reference for label transfer. Cell clusters with low transfer confidence scores were manually re-annotated. Label transfer was performed on L3 late midgut with similar methods, with label transferred and re-annotated L3 early midgut as the reference.

**Neighborhood enrichment.** To calculate spatial enrichment scores of each cell type within each scStereo-seq sample and generate heatmap and bar plot visualization, *squidpy.pl.nhood_enrichment* function from *Squidpy*[27] was used with the mode parameter *zscore*.

**Integration of scRNA-seq and scATAC-seq Data**

**CCA data integration.** scRNA-seq data and scATAC-seq data of CNS and three germ layers (ectoderm, mesoderm, and endoderm) were integrated for analysis. The gene activity matrix for the scATAC-seq data was calculated using *GeneActivity* function from *Signac*. Both scRNA-seq expression matrix and scATAC-seq gene activity matrix were normalized using *NormalizeData* function from *Seurat*. To mitigate biases introduced by different sequencing techniques, CCA was performed using *FindTransferAnchors* function from *Seurat*. CCA was conducted with scATAC-seq gene activity matrix as the query and scRNA-seq expression matrix as the reference to identify transfer anchors. Leveraging these transfer anchors, *TransferData* function from *Seurat* was employed to obtain the imputed scATAC-seq matrix (*refdata* = *`scRNA data matrix`, weight.reduction* = *`scATAC pca.l2 matrix`, dim* = *2:50*). Subsequently, imputed scATAC-seq matrix was merged with scRNA-seq data matrix. Dimension reduction was performed to co-embed them into the same UMAP space. After running *RunPCA* function, the $1^{st}$ to the $50^{th}$ principal components were used to run *RunUMAP* function to obtain the co-embedded UMAP matrix.

**Annotation of integrated data.** Cell type marker genes were generated using *FindAllMarkers* function from *Seurat* with the thresholds minimum cell fraction = 0.1 and logFC > 0.1. Top 50 markers were selected for manual annotation. Top 200 marker genes generated by the *COSG*[28] were also examined for additional reference, where the minimum cell fraction was also set to 0.1. Based on marker genes identified by both methods, clusters were manually curated and annotated **(Table S4)**. For all plots, hex color codes of tissue cell types can be found in **Table S10**.

**Generation of vector fields with *PhyloVelo***

*Phylovelo*[29] was used to generate velocity fields and infer trajectories using monotonically expressed genes. First, duplicate genes were filtered using *drop_duplicate_genes* function. Log-normalization was then performed to filter genes with < 10 read counts. Subsequently, velocities were inferred using the normalized data and the top 5% of genes with the highest Spearman correlations. *k=15* was set for the k-nearest-neighbor (kNN) graph for the final velocity embedding.

**Construction of tissue development trajectory graph**

Based on cell differentiation Alta, Silhouette indexes of adjacent cell types were calculated to evaluate the divergence time between adjacent cell types using *scikit-learn*[30]. The adjacent cell types and their corresponding Silhouette indexes were used as nodes and edges, respectively, to construct an undirected graph. The layout of graph was arranged by using the Yifan Hu proportional algorithm[31] and rendered in *Gephi*[32].

##### Cell differentiation potential analysis

*CytoTRACE*[33] was utilized to estimate the differentiation potential of single cells with default parameters. A sample size of 3,000 were used. *plotCytoTRACE* and *plotCytoGenes* functions were used to visualize the results.

##### Signal pathway activity analysis

Gene components of signaling pathways were obtained from *FlyphoneDB*[34]. scRNA-seq data were filtered by the following criteria: cells with ≥ 200 genes, and genes present in ≥ 3 cells. Filtered data were then subjected to preliminary normalization using *scanpy.pp.normalize_per_cell* function, followed by log transformation with *scanpy.pp.log1p* function. *scanpy.pp.scale* function was applied to perform further data standardization. *scanpy.tl.score_genes* function was used to assess the expression scores of major genes from each signaling pathway in individual cells. *sns.clustermap* function was used to compute the mean expression scores of each pathway across cell types and collection windows and generate heatmaps.

##### TF motif enrichment analysis

Overlapping genes between top 200 markers from *Seurat* and top 200 markers from *COSG* in each cell type in integrated scRNA-seq and scATAC-seq dataC were selected as potential TF target genes. Genomic regions of potential target genes were overlapped with differentially accessible peaks (DA peaks, identified by *FindAllMarker* function of *Seurat*, using test *"LR"* and parameters *minimum cell fraction = 0.1* and *logFC > 0.1*) specific to each cell type. Only peaks annotated as promoter-TSS peaks by the *annotatePeak.pl* program from *Homer*[35] were used for motif enrichment. Motifs from the *CIS-BP* database[36] were enriched in the selected peak regions using the *FindMotifs* function from *Signac*. A *p*-value cutoff of $< 10^{-4}$ was used to filter significant motifs. Activity scores were then generated for these significant motifs. After applying *AddMotifs* function in *Signac*, motif activity score matrix was created using *RunChromVar* function from *Signac*.

##### GRN analysis with *Pando*

*Pando*[37] was used to generate GRN in desired cell groups. For each cell type, motifs retained after enrichment analysis were used as input for the motif position weight matrix (PWM) and overlapping genes between top 200 markers from *Seurat* and top 200 markers from *COSG* in each cell type were used as target genes input. To infer GRNs, TF correlation threshold was set to be ≥ 0.1 and *Signac* method was used for the peak-to-gene assignment. To identify TF-target gene modules, *p*-value threshold was set to 0.05. The number of variables in the model and *R-square* threshold were adjusted based on the total number of modules found (*nvar = 10*, *rsq = 0.2*, or *nvar = 1*, *rsq = 0.05*). After module inference, modules with *p*-values > 0.05 or *p*-values > $10^{-4}$ were discarded, depending on the total number of modules identified. GRN and TF graphs were then visualized.

**Marker gene identification in CNS mature neurons**

To search for distinctive marker genes among mature neuron subtypes, cell subtypes were compared in pairs using *FindMarkers* functions from *Seurat*. Identified marker genes of each cell subtype were then ordered by their occurrences in pairwise comparisons. For each cell subtype, the top 3 marker genes with the most occurrences were retained **(Table S4)**. Such procedure was performed both between groups (e.g., all GABAergic neurons vs. all cholinergic neurons) and within groups (e.g., GABAergic neurons 1 vs. GABAergic neurons 2).

**Trajectory analysis with *STREAM***

*STREAM*[38] was employed for trajectory inference of differentiation from sensory neurons cluster. Dimension reduction was performed as described above in scATAC-seq data processing. Trajectory inference was initiated by setting the initial node as 4 using *seed_elastic_principal_graph* function and performed with *elastic_principal_graph* function with the following parameter settings: *epg_alpha=0.01*, *epg_mu=0.1*, *epg_lambda=0.02*, and *epg_trimmingradius=3*. To ensure the inclusion of additional cells, leaf branches were extended with *extend_elastic_principal_graph* function. Specifically, *epg_ext_mode='WeightedCentroid'* and *epg_ext_par=0.8* settings were used for this purpose.

**Tissue morphometric analysis**

Morphometric trajectories between two scStereo-seq samples were modeled with *spateo.tdr.cell_directions* from *Spateo* with default parameters. Each cell from the earlier time point was assigned to their most likely counterpart from the later time point to generate morphometric vector fields with *SparseVFC* algorithm through *spateo.tdr.morphofiled* function. Subsequently, geometry quantities were calculated with *spateo.tdr.morphofield_acceleration*, *spateo.tdr.morphofield_curvature*, *spateo.tdr.morphofield_curl*, and *spateo.tdr.morphofield_torsion* functions. A general linear model (GLM) regression was performed with *spateo.tl.glm_degs* function to identify genes with highest correlation with morphometric property changes. Gene set enrichment was then performed with *PANGEA*[39].

##### Sub-clustering and re-annotation of endoderm scRNA-seq data

Endoderm cells were isolated from the scRNA-seq *Seurat* object using *subset* function. Cell clusters with germ layer annotation of endoderm were extracted, and the cluster annotated as "hindgut anlage" was excluded. The dimensionality of the data was reduced first by PCA (30 components) on the top 3,000 most variable genes, and then further by UMAP (*dims = 1:30*). Cell clusters were identified first with the *Seurat FindNeighbors* function (*dims = 1:30*), and then *FindClusters* function with a resolution of 1.4. This process resulted in a total of 37 subclusters. Marker genes for each cluster were identified using *Seurat FindAllMarkers* function (*only.pos = TRUE, min.pct = 0.25, logfc.threshold = 0.5*). Clusters were annotated based on these markers. Clusters annotated as "visceral muscle", "neurons", "plasmatocytes", and "proventriculus" were then excluded **(Table S4)**.

##### Midgut marker gene enrichment

Marker genes of each cell subtype were generated using *Findmarker* function of *Seurat* package with parameters *logFC > 0.5* and *pValue < 0.05*. Top 100 markers were used. To screen for more cell type-specific markers, *COSG* package was also used to generate marker genes of each cell type. Common marker genes identified with both methods were kept for the following analysis. Gene ontology and pathway enrichment were identified with *PANGEA*.

##### Gene module identification in midgut cell types

*Hotspot*[40] was utilized to identify co-expressed gene modules in endoderm cells. A weighted kNN graph with 30 neighbors was computed and genes were ranked in descending order based on their *Z* scores. Top 2,000 genes were selected and genes with an auto-correlation FDR > 0.05 were discarded. Gene modules were generated through agglomerative clustering, with the minimum number of genes per module set at 40. 17 modules were identified, and 376 genes were not assigned to a module. *Hotspot* eigengene module scores were determined by *calculate_module_scores* function. Mean scores of each module were calculated based on cell types and visualized using *clustermap* function in *seaborn*[41].

##### RNA velocity field analysis

Based on re-annotated endoderm cells in scRNA-seq data, *Velocyto* command line interface[42] was used to calculate spliced and unspliced transcripts of captured genes. *Dynamo*[43] was used to model differentiation dynamics of cell trajectories. Raw count matrices of spliced and unspliced transcripts were processed with *recipe_monocle* function in *Dynamo*. Top 2,000 highly variable genes were used as feature genes for dimension reduction using UMAP and default parameters. Kinetic parameters were calculated with *dynamo.tl.dynamics* function. Velocity vector flows were projected to 2D UMAP space and visualized with *dynamo.tl.cell_velocities* function. Continuous vector fields were generated in the UMAP space using *vf.VectorField* function. Velocity pseudotime of single cells were determined with *ddhodge* function. Expression patterns of genes of interest were plotted against single-cell pseudotime. Using the vector fields, speed, acceleration, and curvature scores for single cells were calculated with *vf.speed*, *vf.acceleration*, and *vf.curvature* functions, respectively.

##### Gene module analysis in midgut regions

A set of 1,500 regional marker genes from Ref[44] were used for midgut region analysis. These markers were overlapped with genes captured in scStereo-seq data and 1,452 of them were retained to create a kNN graph using *hs.create_knn_graph* function (*weighted_graph=True, n_neighbors=100*) from *Hotspot*. A local correlation matrix was constructed to assess the correlation among cells within each midgut region. Top 50 marker genes of each region were selected for local correlation calculation with *hs.local_correlation_z* function. Gene modules were filtered to retain the ones that contained ≥ 8 genes and had a false discovery rate (FDR) < 0.05. The similarity between gene modules and midgut regions were evaluated by similarity scores, which calculated the ratio of region-specific genes in each module. Identified midgut regions were filtered to retain the ones that contained ≥ 20 cells.

##### Fluorescence *in situ* hybridization

FISH was performed following protocols described in Fly-FISH database[45] with the following modification: Alexa Fluor 488 Tyramide SuperBoost Kit, goat anti-mouse (Invitrogen, B40922) or Alexa Fluor 594 Tyramide SuperBoost Kit, streptavidin (Invitrogen, B40935) was used in the signal developmental step after primary antibody incubation following vendor protocols. Primers to generate RNA probes from embryo cDNA are listed in **Table S9**. Samples were imaged with a Zeiss LSM 980 NLO confocal microscope with a 10 × objective. Imaging results were processed and analyzed with ImageJ[46].
